# A coevolved EDS1-SAG101-NRG1 module mediates cell death signaling by TIR-domain immune receptors

**DOI:** 10.1101/572826

**Authors:** Dmitry Lapin, Viera Kovacova, Xinhua Sun, Joram Dongus, Deepak D. Bhandari, Patrick von Born, Jaqueline Bautor, Nina Guarneri, Johannes Stuttmann, Andreas Beyer, Jane E. Parker

**Author notes:** **Corresponding author:** Jane E. Parker. these authors contributed equally (Dmitry Lapin, Viera Kovacova, Xinhua Sun). Laboratory of Nematology, Wageningen University, Droevendaalsesteeg 1, 6708 PB, Wageningen, the Netherlands. Material distribution policy: The authors responsible for distribution of materials integral to the findings presented in this study in accordance with the policy described in the Instructions for Authors (www.plantcell.org) are: Jane E. Parker and Dmitry Lapin.

## Abstract

Plant intracellular nucleotide-binding/leucine-rich repeat (NLR) immune receptors are activated by pathogen effectors to trigger host defenses and cell death. Toll-Interleukin1-receptor (TIR)-domain NLRs (TNLs) converge on the Enhanced Disease Susceptibility1 (EDS1) family of lipase-like proteins for all resistance outputs. In Arabidopsis TNL immunity, *At*EDS1 heterodimers with Phytoalexin Deficient4 (*At*PAD4) transcriptionally boost basal defense pathways. *At*EDS1 uses the same surface to interact with PAD4-related Senescence-Associated Gene101 (*At*SAG101), but the role of *At*EDS1-*At*SAG101 heterodimers was unclear. We show that *At*EDS1-*At*SAG101 function together with *At*NRG1 coiled-coil domain helper NLRs as a coevolved TNL cell death signaling module. *At*EDS1-*At*SAG101-*At*NRG1 cell death activity is transferable to the solanaceous species, *Nicotiana benthamiana*, and cannot be substituted by *At*EDS1-*At*PAD4 with *At*NRG1 or *At*EDS1-*At*SAG101 with endogenous *Nb*NRG1. Analysis of EDS1-family evolutionary rate variation and heterodimer structure-guided phenotyping of *At*EDS1 variants or *At*PAD4-*At*SAG101 chimeras identify closely aligned α-helical coil surfaces in the *At*EDS1-*At*SAG101 partner C-terminal domains that are necessary for TNL cell death signaling. Our data suggest that TNL-triggered cell death and pathogen growth restriction are determined by distinctive features of EDS1-SAG101 and EDS1-PAD4 complexes and that these signaling machineries coevolved with further components within plant species or clades to regulate downstream pathways in TNL immunity.

## Introduction

In plants, immunity to host-adapted pathogens is mediated by large, diversified families of intracellular nucleotide-binding/leucine-rich repeat (NLR) receptors whose members recognize specific pathogen virulence factors (effectors) that are delivered into host cells to promote infection (Baggs et al., 2017). NLRs are ATP/ADP-binding molecular switches and their activation by effectors involves intra-and intermolecular conformational changes, which lead to rapid host gene expression changes, induction of antimicrobial pathways and, often, localized cell death called a hypersensitive response (HR) (Cui et al., 2015; Jones et al., 2016). A signature of plant NLR immunity is the induction of multiple transcriptional sectors which can buffer the host against pathogen interference (Tsuda et al., 2013; Cui et al., 2018; Mine et al., 2018; Bhandari et al., 2019). How NLR receptors initiate downstream resistance pathways in effector-triggered immunity (ETI) remains unclear.

Two major pathogen-sensing NLR receptor classes, TIR-NLRs (also called TNLs) and CC-NLRs (CNLs), are broadly defined by their N-terminal Toll/interleukin-1 receptor (TIR) or coil-coiled (CC) domains. Evidence suggests that these domains serve in receptor activation and signaling (Cui et al., 2015; Zhang et al., 2016a).Different NLR protein families characterized in Arabidopsis and *Solanaceae* species function together with pathogen-detecting (sensor) NLRs in ETI and are thus considered ‘helper’ NLRs (Wu et al., 2018a) that might bridge between sensor NLRs and other immunity factors. Members of the *NRC* (*NLR required for HR-associated cell death*) gene family, which expanded in Asterids, signal in ETI conferred by partially overlapping sets of phylogenetically related sensor CNLs (Wu et al., 2017). Two sequence-related NLR groups, N Required Gene1 (NRG1, (Peart et al., 2005; Qi et al., 2018)) and Accelerated Disease Resistance1 (ADR1, (Bonardi et al., 2011)) signaling NLRs, were originally classified by a distinct CC domain sequence (referred to as CCR) shared with the Arabidopsis Resistance to Powdery Mildew8 (RPW8) family of immunity proteins (Collier et al., 2011). Subsequent analysis revealed that although monocot ADR1s lack the CCR (Zhong and Cheng, 2016), these NLRs share a phylogenetically-distinct nucleotide-binding domain (Shao et al., 2016; Zhong and Cheng, 2016). Studies of *NRG1* and *ADR1* mutants in Arabidopsis and *Nicotiana benthamiana* revealed important roles of the genes in ETI (Peart et al., 2005; Bonardi et al., 2011; Dong et al., 2016; Castel et al., 2018; Qi et al., 2018; Wu et al., 2018b).

Notably, *NRG1* genes are necessary for eliciting host cell death in several TNL but not CNL receptor responses (Castel et al., 2018; Qi et al., 2018). By contrast, three Arabidopsis *ADR1* genes (*AtADR1, AtADR1-L1* and *AtADR1-L2*) act redundantly in signaling downstream of CNL and TNL receptors (Bonardi et al., 2011; Dong et al., 2016; Wu et al., 2018b). In resistance to bacteria triggered by the Arabidopsis sensor CNL Resistant to *Pseudomonas syringae*2 (RPS2), *ADR1*-family genes stimulated accumulation of the disease resistance hormone salicylic acid (SA) and cell death (Bonardi et al., 2011). Analysis of flowering plant (angiosperm) genomes indicated presence of *NRG1* and *TNL* genes in eudicot lineages and loss of these genes from monocots and several eudicots (Collier et al., 2011; Shao et al., 2016; Zhang et al., 2016b). By contrast, Arabidopsis *ADR1* orthologs are present in eudicot and monocot species (Collier et al., 2011; Shao et al., 2016). So far, there was no evidence of molecular interactions between sensor and helper NLRs.

All studied TNL receptors, activated by pathogen effectors in ETI or as auto-active molecules (producing autoimmunity), signal via the non-NLR protein, Enhanced Disease Susceptibility1 (EDS1) for transcriptional defense reprograming and cell death (Wiermer et al., 2005; Wirthmueller et al., 2007; García et al., 2010; Xu et al., 2015; Adlung et al., 2016; Ariga et al., 2017; Qi et al., 2018). EDS1 is therefore a key link between TNL activation and resistance pathway induction. Consistent with an early regulatory role in TNL signaling, *At*EDS1 interacts with a number of nuclear TNLs (Bhattacharjee et al., 2011; Heidrich et al., 2011; Kim et al., 2012). Also, an interaction between *Nb*EDS1a and *Nb*NRG1 was recently reported (Qi et al., 2018). Together with Phytoalexin Deficient4 (PAD4) and Senescence-Associated Gene101 (SAG101), EDS1 constitutes a small family which was found in angiosperms but not non-seed species, post-dating the origin of *NLR* genes in plants (Wagner et al., 2013; Gao et al., 2018). Phylogenetic sampling of 16 angiosperm species indicated that *EDS1* and *PAD4* are present in eudicots and monocots, whereas *SAG101* (like *NRG1* and *TNLs*) was not detected in monocot genomes (Collier et al., 2011; Wagner et al., 2013).

The three EDS1-family proteins possess an N-terminal α/β-hydrolase fold domain with similarity to eukaryotic class-3 lipases and a unique C-terminal α-helical bundle, referred to as the ‘EP’ domain (pfam id: PF18117) (Wagner et al., 2013). *At*EDS1 forms exclusive heterodimers with *At*PAD4 and *At*SAG101 through N-and C-terminal contacts between the partner domains (Feys et al., 2001; Feys et al., 2005; Rietz et al., 2011; Wagner et al., 2013). Genetic, molecular and protein structural evidence from Arabidopsis revealed a function of *At*EDS1 heterodimers with *At*PAD4 in basal immunity that is boosted by TNLs in ETI via an unknown mechanism (Feys et al., 2005; Rietz et al., 2011; Bhandari et al., 2019). EDS1-PAD4 basal immunity limits the growth of infectious (virulent) pathogens without host cell death and is thus thought to reflect a core EDS1-PAD4 immunity function (Zhou et al., 1998; Rietz et al., 2011; Cui et al., 2017). In Arabidopsis accession Col-0 (Col), ETI conferred by the nuclear TNL pair Resistant to *Ralstonia solanacearum* 1S - Resistant to *Pseudomonas syringae* 4 (RRS1S-RPS4) recognizing *Pseudomonas syringae* effector AvrRps4 has been used extensively to investigate *At*EDS1-*At*PAD4 signaling (Heidrich et al., 2011; Saucet et al., 2015). Col *RRS1S-RPS4* ETI is associated with a weak cell death response (Heidrich et al., 2011), and to bolster basal immunity, *At*EDS1-*At*PAD4 complexes steer host transcriptional programs towards SA-induced defenses and away from SA-antagonizing jasmonic acid (JA) pathways (Zheng et al., 2012; Cui et al., 2018; Bhandari et al., 2019). This signaling involves positively charged amino acids at an *At*EDS1 EP domain surface lining a cavity formed by the EDS1-family heterodimers (Bhandari et al., 2019). The function of EDS1-SAG101 complexes in TNL ETI was not determined, although *At*SAG101 but not *At*PAD4 was required for autoimmunity conditioned by the TNL pair Chilling Sensitive 3/Constitutive Shade Avoidance 1 (CHS3/CSA1) (Xu et al., 2015). Also, TNL ETI but not basal immunity was retained in Arabidopsis accession Ws-2 expressing an *At*EDS1 variant (EDS1^L262P^) which formed stable complexes with SAG101 but not PAD4 (Rietz et al., 2011). These data suggest that *At*EDS1-*At*PAD4 and *At*EDS1-*At*SAG101 heterodimers have distinctive roles in TNL ETI.

Here we examine EDS1-family sequence variation across seed plant lineages and test whether EDS1-PAD4 and EDS1-SAG101 complexes are functionally transferable between different plant groups. Despite high levels of conservation, we find there are barriers to EDS1 heterodimer functionality between plant lineages. By measuring TNL immunity resistance and cell death outputs in Arabidopsis and tobacco (*Nicotiana benthamiana*) ETI pathway mutants, we establish that *At*EDS1 and *At*SAG101 cooperate with *At*NRG1 but not with tobacco NRG1 (*Nb*NRG1) in TNL cell death signaling. We provide evidence that *At*EDS1 and *At*PAD4 have a different immunity role that limits bacterial pathogen growth. A structure-guided analysis of *At*EDS1 and *At*PAD4/*At*SAG101 variants indicates decision-making between cell death and bacterial growth inhibition branches in TNL (*RRS1S-RPS4*) immunity is determined by distinctive features of the EDS1-SAG101 and EDS1-PAD4 complexes. Our data suggest that signaling machineries co-evolved within plant species and clades for regulating downstream pathways in TNL immunity.

## Results

### Dicot plants from the order *Caryophyllales* lack predicted SAG101 orthologs

A previous study showed that EDS1 and PAD4 encoding genes are present in flowering plants (angiosperms) (Wagner et al., 2013). Here, we investigated the distribution of EDS1-family members using recent genomic information. Analysis of protein-sequence orthogroups from genomes of 52 green plants shows that EDS1 and PAD4 are present in 46 seed plant species, including conifers (Supplemental Table 1, Supplemental Dataset 5, 6, 7), suggesting that the EDS1 family arose in a common ancestor of gymno-and angiosperms. We did not detect EDS1-family orthologs in the aquatic monocot *Spirodela polyrhiza* (duckweed). As reported (Wagner et al., 2013), *At*SAG101 orthologs are absent from monocots and the basal eudicot *Aquilegia* and *Erythranthe guttata* (order *Lamiales*, formerly *Mimulus guttatus*). Here, SAG101 was also not found in conifers or the eudicot species *Beta vulgaris* (sugar beet) from the order *Caryophyllales* (Supplemental Table 1). Reciprocal BLAST searches failed to identify putative *At*SAG101 orthologs in genomes and transcriptomes of nine additional *Caryophyllales* genomes (quinoa, amaranth, six species from *Silene* genus and spinach). We concluded that loss of *SAG101* is likely common not only to monocots but also *Caryophyllales* eudicot species.

Next, we used 256 sequences of EDS1-family orthologs identified with OrthoMCL, and additional BLAST searches to infer fine-graded phylogenetic relationships (Figure 1, Supplemental Figure 1, see Methods). On a maximum likelihood phylogenetic tree (Figure 1A), EDS1, PAD4 and SAG101 predicted proteins of flowering plants form clearly separated nodes. Conifer EDS1 and PAD4 belong to distinct clades that do not fall into the EDS1 and PAD4 of flowering plant groups. Therefore, functions of EDS1-family proteins might have diverged significantly between conifers and flowering plants. Conifer EDS1 further separated into two well-supported branches. Analyzed *Solanaceae* genomes (with the exception of pepper, *Capsicum annuum*) encode SAG101 proteins in two well-supported groups, we refer to as A and B (Figure 1A), suggesting SAG101 diversification within *Solanaceae*. Because the EDS1-family tree topology is not known, we also performed a Bayesian inference of phylogeny (MrBayes phylotree; Supplemental Figure 1A), which supported conclusions made from the maximum likelihood tree analysis (Figure 1A).

**Figure 1.**
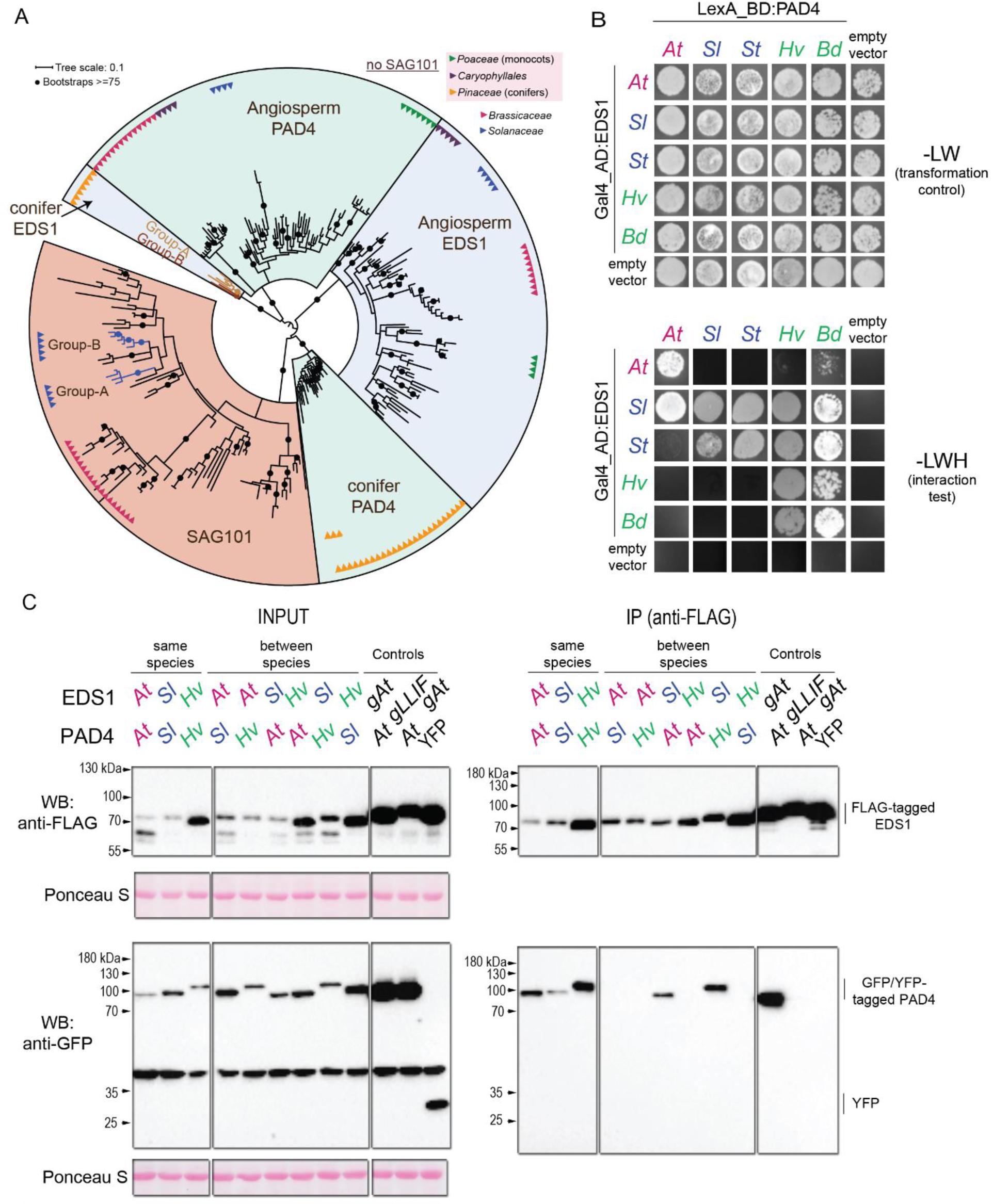
Phylogeny of EDS1 family proteins in seed plants and conservation of EDS1-PAD4 interactions in angiosperms. (A) Maximum-likeiihood tiee of 256 sequences from predicted EDSI-family proteins. Branches with Feteenstein’s bootstrap support values ≥ 75 are given as black dots EDSI PALM and SAGI01 orthologs form separate groups. Solanaceae SAG101 fall into two groups - A and B. Similarly, conifers have two EDSI groups labeled A and B (B) A yeast-two-hybnd assay testing interactions between EDS1 and PAD4 from Arabidopsis (*At*). tomato (*SI*). potato (*St*). barley (*Hv*) and Brachypodium (*Bd*). -LW drop out selection medium without leucine and tryptophan-LWH medium without leucine, tryptophan and histidine. Each comb-nation shown was tested n 2-4 independent experiments with similar results (C) Western blot analysis showing proteins before (INPUT) and after immunoprecipitation (IP) assay to test *in planta* interactions between Arabidopsis (*At*). tomato (*SI*) and barley (*HV*) EDS1 and PAD4 orthologs as indicated, and detected with a-FLAG or a-GFP antibodies. Proteins were transiently co-expressed in ***N. benthamiai*** IPs using Arabidopsis *(At) pEDS1:gEDS1-3xFLAG* (*gAt)* or (At) pEDS1:gEDSU3xFLAG (gAt) or p£DS1:gEDSl_LUF/AAAA-3xFLAG (gLLIF) with Arabidopsis (At) 35SPAD4-YFP served as positive and negative controls, respectively Ponceau S staining of the membrane shows equal loading o’ input samples Comb-nations tested in four independent experiments with the exception of FLAG-S€DST’YFP-WvPAD4 and FLAG-Hw€DS1i’YFP-S^5AD4. which were repeatec twice.

Although *Brassicaceae, Caryophyllales* and *Poaceae* EDS1 and PAD4 form well supported clades (Figure 1A, Supplemental Figure 1A), generally the EDS1-family does not provide sufficient resolution to separate other groups within flowering plants. This might be explained by conservation of the proteins and negative selection. Indeed, EDS1 sequences in *Brassicaceae, Solanaceae* and *Poaceae* appear to have evolved mainly under purifying selection constraints (62.0-88.4% of sites, Supplemental Table 2). Mapping of evolutionary rates obtained with the refined EDS1, PAD4 and SAG101 phylogenetic trees (see Methods) showed that slowly evolving (conserved) amino acids are present in the core lipase-like domain α/β-hydrolase folds and EP domain α-helical bundles likely to preserve structural stability, but also on the partner EP domain surfaces lining the heterodimer cavity (Supplemental Figure 1B). Several conserved amino acids on this cavity surface of *At*EDS1 are essential for TNL immunity (Bhandari et al., 2019). The hydrophobic character of ‘LLIF’ α-helix (H) in the *At*EDS1 lipase-like domain, which contacts hydrophobic pockets in corresponding *At*PAD4 and *At*SAG101 domains (Wagner et al., 2013), is also conserved across species (Supplemental Figure 1C). While EDS1 sequences in three flowering plant families appear to have evolved mainly under purifying selection (Supplemental Table 2), further analysis of evolutionary constraints indicated positive selection in *Brassicaceae* EDS1 sequences at five positions with multinucleotide mutations: R16, K215, Q223, R231 and K487 (Col *At*EDS1 AT3G48090 coordinates; Supplemental Figure 1D, 1E, Supplemental Table 2). These amino acids are surface-exposed on the crystal structure of *At*EDS1, the first four being located in the lipase-like domain, and K487 in the EP domain (Supplemental Figure 1E). Whether this variation has adaptive significance is unclear since an *At*EDS1^K487R^variant retained TNL immunity function (Bhandari et al., 2019).

In summary, we find that EDS1 and PAD4 orthologs are present in conifers as well as flowering plants and form phylogenetically distinct sequence groups in these lineages. This suggests an origin of the EDS1 family in a common ancestor of seed plants. Also, multiple species of the eudicot lineage *Caryophyllales* lack SAG101 orthologs, suggesting that loss of *SAG101* is not a sporadic event in the evolution of eudicots.

### Interactions between EDS1-family proteins from eudicot and monocot species

High conservation at the lipase-like EP domain interfaces of EDS1-SAG101 and EDS1-PAD4 (Supplemental Figure 1B) would be in line with heterodimer formation between partners in species other than Arabidopsis and, potentially, interactions between EDS1 and PAD4 or SAG101 originating from different phylogenetic groups. We therefore tested EDS1-family proteins interactions within and between representative species of eudicot (*Brassicaeae, Solanaceae*) and monocot (*Poaceae*) families. In yeast-2-hybrid (Y2H) assays, EDS1 and PAD4 from the same species or family formed a complex (Figure 1B). Notably, tomato (*Solanum lycopersicum*) EDS1 (*Sl*EDS1) also interacted with *At*PAD4 and barley (*Hordeum vulgare*) and *Brachypodium distachyon* PAD4 proteins (*Hv*PAD4 and *Bd*PAD4) (Figure 1B). By contrast, *At*EDS1 and monocot *Hv*EDS1 or *Bd*EDS1 did not interact with either *Sl*PAD4 or potato (*Solanum tuberosum*) *St*PAD4. These data show that EDS1 and PAD4 partners in species that are distant from Arabidopsis also interact physically and there are some between-clade associations.

We selected Arabidopsis, tomato and barley EDS1-PAD4 combinations for *in planta* co-immunoprecipitation (IP) assays of epitope-tagged proteins transiently expressed after *Agrobacterium tumefaciens* (agroinfiltration) of *N. benthamiana* leaves. As expected, *At*PAD4-YFP immunoprecipitated (IPed) with coexpressed *At*EDS1-FLAG. This interaction was strongly reduced when the *At*EDS1 ‘LLIF’ N-terminal heterodimer contact was mutated ((Wagner et al., 2013), Figure 1C). In accordance with the Y2H data, YFP-*Sl*PAD4 and YFP-*Hv*PAD4 IPed with FLAG-EDS1 from the same species (*Sl*EDS1 and *Hv*EDS1). Also, FLAG-*Sl*EDS1 interacted with YFP-*At*PAD4 and YFP-*Hv*PAD4, but FLAG-*At*EDS1 did not interact with either YFP-*Sl*PAD4 or YFP-*Hv*PAD4 (Figure 1C). Similarly, FLAG-*Hv*EDS1 failed to interact with YFP-*At*PAD4 or YFP-*Sl*PAD4 (Figure 1C). Hence, EDS1 and PAD4 from the same eudicot or monocot species form stable complexes *in planta* like *At*EDS1-*At*PAD4, suggesting that EDS1-PAD4 heterodimer formation is a conserved feature across angiosperms. In Y2H and *in planta*, between-clade complex formation is not universal, indicating that there are barriers to certain EDS1-PAD4 partner interactions between distant lineages.

We also tested in *N. benthamiana* transient assays whether *Sl*EDS1 or *At*EDS1 can form complexes with SAG101 proteins from *Solanaceae* (*N. benthamiana*) and Arabidopsis (Supplemental Figure 2). *S. lycopersicum* has two *SAG101* genes, which fall respectively into *Solanaceae* SAG101 groups A and B (Figure 1A, Supplemental Figure 1A) and are most sequence-related to *N. benthamiana Nb*SAG101a and *Nb*SAG101b (81.52 and 72.44% sequence identity; Supplemental Dataset 1). As expected, *At*EDS1-FLAG interacted with *At*SAG101-YFP in IP assays (Supplemental Figure 2A). Also, FLAG-*Sl*EDS1 interacted with *Nb*SAG101a-GFP and *Nb*SAG101b-GFP, consistent with the close phylogenetic relationship between cultivated tomato and *N. benthamiana*. Notably, FLAG-*Sl*EDS1 IPed *At*SAG101-YFP, but *At*EDS1-FLAG did not IP *Nb*SAG101a or *Nb*SAG101b (Supplemental Figure 2A), similar to the *At*EDS1/*Sl*PAD4 combinations (Figure 1B and 1C). As shown previously (Feys et al., 2005), *At*SAG101-YFP localized to the nucleus, whereas *Nb*SAG101a-GFP and *Nb*SAG101b-GFP had a nucleocytoplasmic distribution in *N. benthamiana* (Supplemental Figure 2B). Together, the data suggest that EDS1-partner interactions are conserved across angiosperms, but there are some restrictions to protein interactions between different taxonomic groups.

### Tomato EDS1-PAD4 are functional in Arabidopsis *TNL RPP4* immunity

The above data show that tomato EDS1 (*Sl*EDS1) forms a stable complex with tomato and Arabidopsis PAD4 proteins (Figure 1B and 1C). We therefore tested whether *Sl*EDS1-*Sl*PAD4 signal together or with respective *At*EDS1 and *At*PAD4 partners in Arabidopsis TNL immunity (Figure 2). For this, four EDS1-PAD4 co-expression constructs (*At*EDS1-*At*PAD4, *Sl*EDS1-*Sl*PAD4, *Sl*EDS1-*At*PAD4, *At*EDS1-*Sl*PAD4; Figure 2A) were transformed into a triple *eds1-2 pad4-1 sag101-3* mutant line in accession Col-0 (Col). Arabidopsis or tomato EDS1 and PAD4 coding sequences were fused to N-terminal FLAG and YFP tags, respectively. Expression of the genes was driven by Arabidopsis *EDS1* and *PAD4* promoters (Gantner et al., 2018). Three independent transgenic lines expressing the tagged proteins (Figure 2A) were selected and spray-inoculated with the downy mildew pathogen *Hyaloperonospora arabidopsidis* (*Hpa*) isolate Emwa1, which is recognized in Col by TNL receptor RPP4 (Van Der Biezen et al., 2002). Pathogen spores were quantified on leaves at 7 d post inoculation (dpi). Col expressing StrepII-3xHA-YFP was resistant to *Hpa* Emwa1 while *eds1-2 pad4-1 sag101-1* and accession Ws-2 (which lacks *RPP4* (Holub, 1994)) were susceptible (Figure 2B). The *At*EDS1-*At*PAD4 pair fully restored *Hpa* resistance in *eds1-2 pad4-1 sag101-3* (Figure 2B), consistent with an EDS1-PAD4 heterodimer being necessary and sufficient for TNL immunity in Arabidopsis (Glazebrook et al., 1997; Feys et al., 2001; Rietz et al., 2011; Wagner et al., 2013). The *Sl*EDS1-*Sl*PAD4 pair also conferred full *RPP4* immunity (Figure 2B). Thus, *Sl*EDS1-*Sl*PAD4 is functionally transferable from tomato to Arabidopsis. By contrast, between-species EDS1-PAD4 combinations *At*EDS1-*Sl*PAD4 and *Sl*EDS1-*At*PAD4 did not fully prevent *Hpa* sporulation. While no *RPP4* resistance was detected in Arabidopsis lines expressing *At*EDS1-*Sl*PAD4 (which did not interact in Y2H and IP assays (Figures 1B and 1C), there was a partial resistance response in plants expressing *Sl*EDS1-*At*PAD4 (Figure 2B), which did interact (Figure 1B and 1C). We reasoned that the between-clade *Sl*EDS1-*At*PAD4 combination likely retains some TNL resistance signaling function because it can form a heterodimer (Figure 1B and 1C), but incompatibility with Arabidopsis factors might prevent it from functioning fully in Arabidopsis TNL (*RPP4*) signaling.

**Figure 2.**
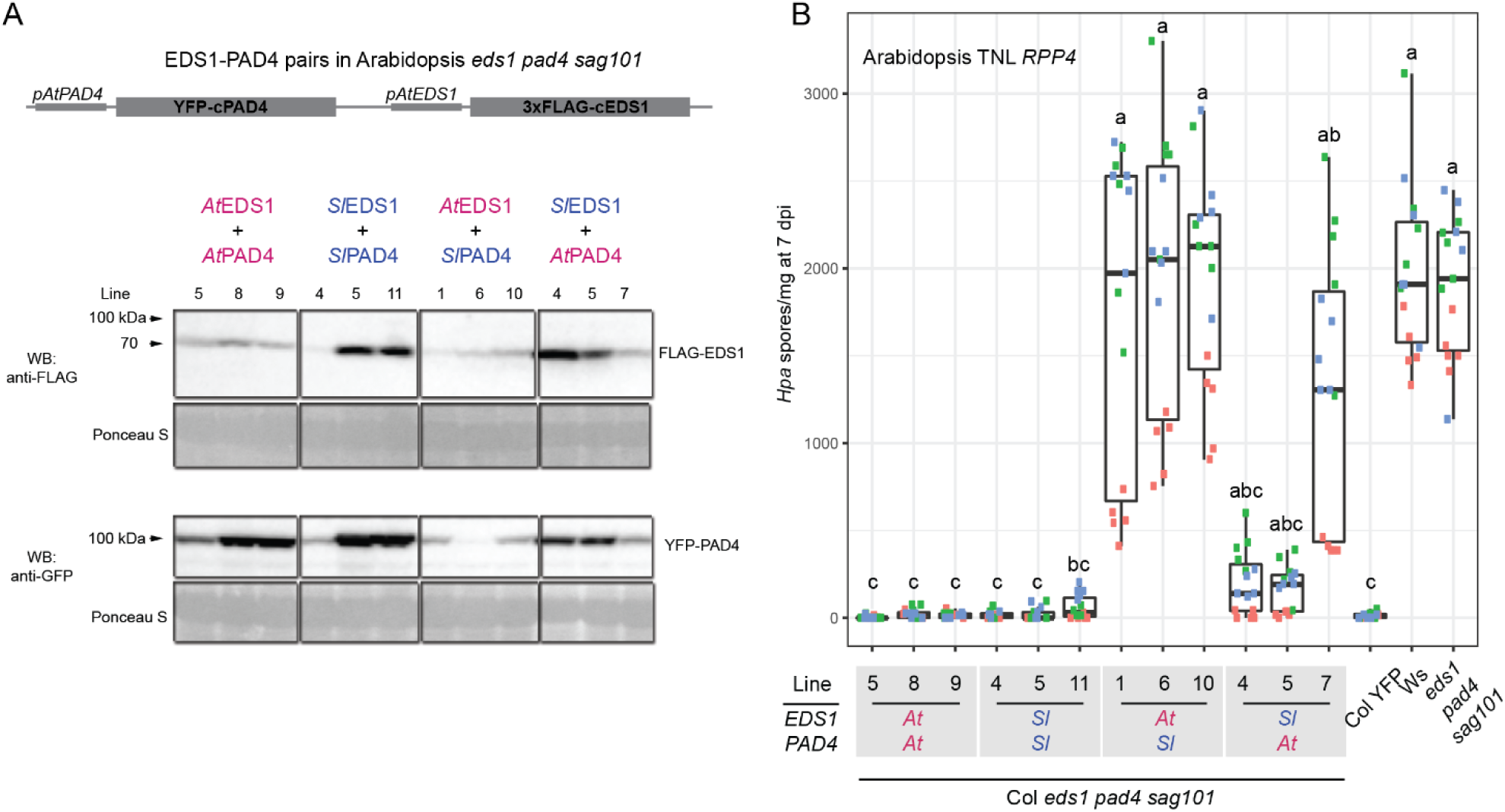
A tomato EDS1-PAD4 pair functions in Arabidopsis TNL *RPP4* immunity. (A) Upper panel: schematic representation of a Golden Gate assembled fragment to express coding sequences of Arabidopsis (*At*) and tomato (*SI*) EDS1 and PAD4 under the control of corresponding Arabidopsis promoters in Arabidopsis Col *eds1-2 pad4*-*1 sag101-3.* Lower panel: Western blot analysis FLAG-EDS1 and YFP-PAD4 proteins in Arabidopsis T3 independent transgenic lines, as indicated, after *Hpa* Emwal infection (3 dpi. panel B). Ponceau S staining of the membrane served as a loading control. The analysis was performed twice with similar results. (B) A TNL (*RPP4*) resistance assay in the same T3 independent transgenic lines as shown in (A. lower panel). *Hpa* Emwal comdiospores on leaves were quantified at 7 dpi. An Arabidopsis Col YFP (resistant). Ws-2 (*rpp4*. susceptible) and non transformed *edsl-2 pad4-1 swg101-1* (susceptible) served as controls. Data from three independent experiments (biological replicates) are represented on a box-plot with dots in the same color corresponding to technical replicates (individual normalized spore counts) from one independent experiment. Genotypes sharing letters above box-whiskers on the plot do not show statistically significant differences (Nemenyi test with Bonferrom correction for multiple testing. a=0.01, n=l5).

### *Sl*EDS1 with *Sl*SAG101b confer *N. benthaniama* TNL *Roq1* immunity

To explore further whether EDS1-family members are functionally transferable between eudicot species for TNL immunity, we exploited the *N. benthamiana TNL Recognition of XopQ1* (*Roq1*) resistance system. Roq1 recognizes the Type III-secreted effector XopQ delivered from leaf-infecting *Xanthomonas campestris* pv. *vesicatoria* (*Xcv*) bacteria (Adlung et al., 2016; Schultink et al., 2017). This recognition induces *NbEDS1a*-dependent cell death and resistance to pathogen growth (Adlung et al., 2016; Schultink et al., 2017; Qi et al., 2018). We used Agrobacteria-mediated transient expression of proteins in *N. benthamiana* in combination with simultaneous *Xcv* infiltration or XopQ-myc agroinfiltration of leaf sectors to monitor TNL *Roq1* resistance and cell death, respectively (Figure 3, Supplemental Figure 3). For the assays (setup outlined on Figure 3A), WT *N. benthamiana* and reported *Nbeds1a, Nbpad4* single and *Nbeds1a pad4* double mutants (Ordon et al., 2017), as well as *N. benthamiana pad4 sag101a sag101b* triple mutant (abbreviated to *Nb-pss*, (Gantner et al., 2019)) were used. We first tested whether agroinfiltration interferes with *Roq1* resistance to *Xcv* growth (Supplemental Figure 3A). Streptomycin (at 150-200 ug/l) allowed selective growth on agar plates of *Xcv* but not *A. tumefaciens* strain GV3101 bacteria extracted from leaves. *A. tumefaciens* infiltrated at two densities (OD_600_ 0.1 and 0.4, strain to express YFP) did not affect *Xcv* growth in susceptible *Nbeds1a* but further reduced low *Xcv* proliferation in resistant WT *N. benthamiana* leaves (Supplemental Figure 3A). We took the 2.5-3 log_10_ difference in *Xcv* titers between WT and *Nbeds1a* upon agroinfiltration as a measure of *EDS1*-dependent *TNL* resistance to *Xcv* growth.

**Figure 3.**
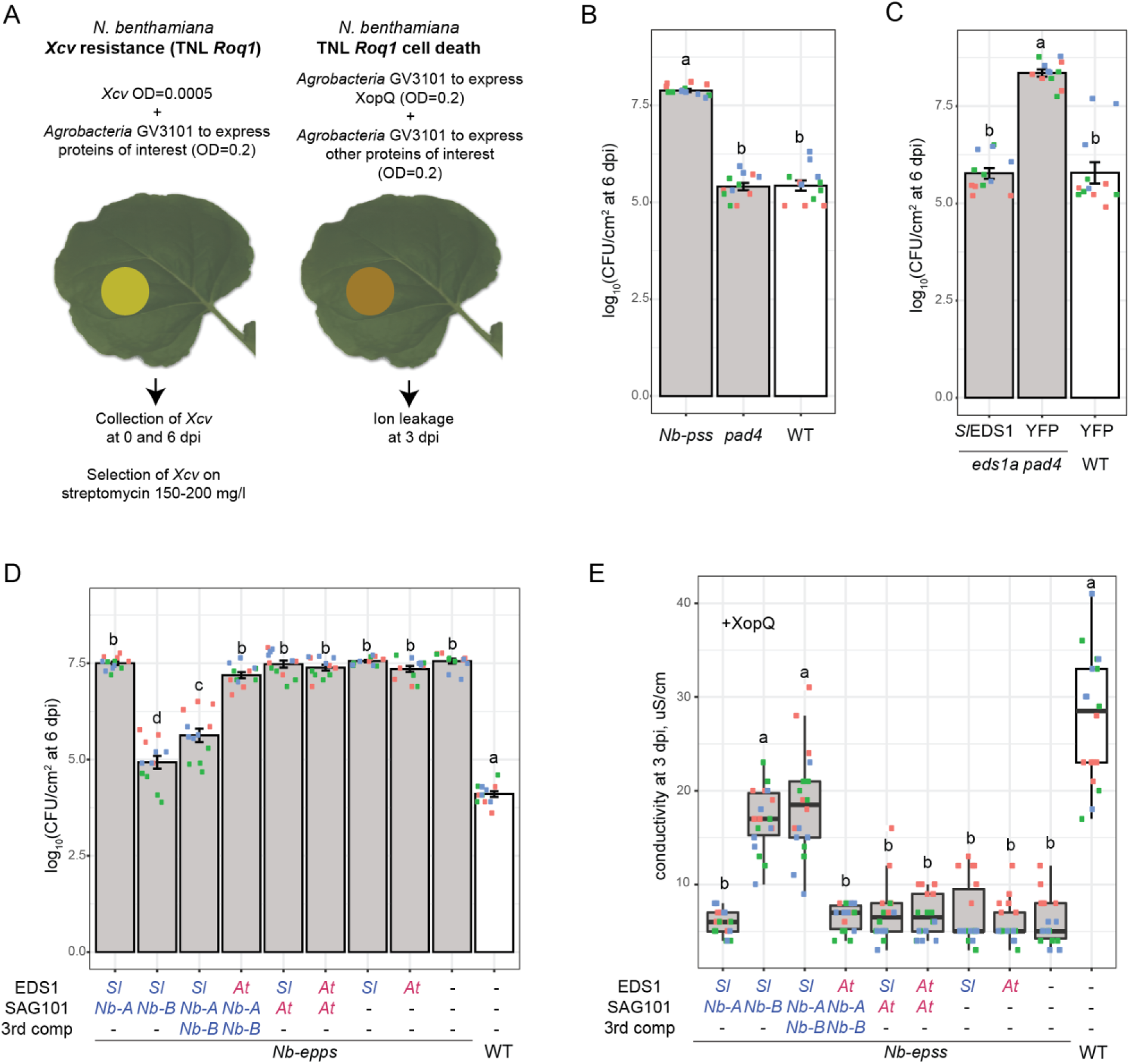
Tomato or *N. benthamiana* EDS1 signal with AtoSAGIOlb in TNL *Roql* immunity and cell death. (A) Schematic showing *N. benthamiana* transient complementation assays to test protein functionalities in *Roq1* immunity at the level of *Xarthomonas campestris* pv. *vesicatoria (Xcv)* growth inhibition and XopQ-tnggered cell death (see Methods for details). (B) *N benthamiana pad4* plants display similar resistance as WT to *Xcv* bactena at 6 dpi (00600=0 0005 after coinfiltration with *A tumefociens* expressing YFP). By contrast. *Xcv* growth is ∼2.5 log_10_ higher in an *N. benthamiana pad4 sag101a sag101b (Nb-pss)* mutant Dots of the same color on box-plots correspond to technical replicates (individual bacterial extractions) from one of three independent experiments (biological replicates). Genotypes shanng letters above bars do not show statistically significant differences (Tukey’s HSD. α=0.001. n=12). Error bars represent ±SEM. (C) Transiont oxprossion of *pAtEDS 1 FLAG-SIEDS1* fully complements susceptibility of *N. benthomiana eds 1a pod4* plants at the level of *Xcv* growth Overlapping letter codes above bars show that differences between genotypes are not statistically significant (Tukey’s HSD, α=0.001, n=12 from three independent experiments used as biological replicates). Error bars represent ±SEM (D) Complementation of susceptibility in *N benthamiana eds1a pnd4 sag101a sag101b (Nb-epss)* to *Xcv* by transient expression of *pAtEDSl FLAG-SIEDS1 (SI), 35S NbSAG10la-GFP (Nb-A). 35S:NbSAG101b-GFP (Nb-B), pAtEDS1:AtEDS1-YFP(At). 35SAtSAG101-YFP (At)* or *35S YFP*(“-”). as indicated Co-expression of FLAG-S/EDS1 and *Nb*SAG101b-GFP is sufficient to suppress *Xcv* growth almost to the level of WT plants No combination with Arabidopsis proteins restored Xcv resistance. Dots of the same color in box plots represent technical replicates (individual extractions of bacteria) in one of three independent experiments (biological replicates). The same letters above bars indicate that differences in means are not statistically significant between genotypes (Tukey’s HSD, α=0 001, n=12). Error bars represent ±SEM. (E) Complementation of *Roq1* cell death triggered by XopQ-myc in *Nb-epss* plants via transient expression of the same protein combinations as in the panel D. Cell death was measured as an increase in conductivity relative to a YFP negative control (all in sample descnption). The experiment was repeated three times with six leaf discs used as technical replicates (same colored dots correspond to replicates in each independent experiment used a biological replicate) Statistical significance of differences between samples v/as assessed using a Nemenyi test with Bonferroni correction for multiple testing (α=0.01, n=18).

We infiltrated WT *N. benthamiana, Nbpad4* and *Nb-pss* mutant lines with *Xcv* (in the presence of *A. tumefaciens*). Whereas WT and *Nbpad4* limited *Xcv* growth, *Nb-pss* was susceptible to *Xcv* (Figure 3B), suggesting that one or both *NbSAG101* genes are essential whereas *NbPAD4* is dispensable for *Roq1* immunity. Supporting this, susceptibility to *Xcv* infection in the *Nbeds1a pad4* double mutant was converted to full resistance after *Agrobacteria*-mediated expression of FLAG-*Sl*EDS1 but not YFP (Figure 3C). This also shows that FLAG-*Sl*EDS1 is functional in *N. benthamiana Roq1*-dependent *Xcv* growth restriction. To test activities of *Nb*SAG101a or *Nb*SAG101b individually in *Roq1* immunity, an *Nbeds1a pad4 sag101a sag101b* quadruple mutant (*Nb-epss*) was selected from a cross between *Nbeds1a* with *Nb-pss* (Ordon et al., 2017; Gantner et al., 2019). Transient co-expression of *Nb*SAG101b-GFP but not *Nb*SAG101a-GFP with functional FLAG-*Sl*EDS1 (Figure 3C) restored resistance to *Xcv* growth in Nb-*epss*, although not completely to the level of WT *N. benthamiana* (Figure 3D). Also, FLAG-*Sl*EDS1 functioned with *Nb*SAG101b-GFP, but not *Nb*SAG101a-GFP, in conferring *Roq1*-dependent cell death, as quantified in a leaf disc ion leakage assay at 3 dpi of XopQ-myc with the protein combinations (Figure 3E). Western blot analysis at 2 dpi showed that *Nb*SAG101a-GFP and *Nb*SAG101b-GFP accumulated to similar levels in these assays (Supplemental Figure 3B). We concluded that *Nb*SAG101b, but not *Nb*SAG101a or *Nb*PAD4, functions together with *Sl*EDS1 or endogenous *Nb*EDS1a in *N. benthamiana Roq1* immunity.

### *At*EDS1 with *At*SAG101 do not restore TNL *Roq1* signaling in *Nb-epss* leaves

In *N. benthamiana* cell death and resistance assays, we tested whether *At*EDS1-*At*SAG101 or the heterologous interacting *Sl*EDS1-*At*SAG101 and non-interacting *At*EDS1-*Nb*SAG101b pairs (Supplemental Figure 2A) could substitute for endogenous *Nb*EDS1a and *Nb*SAG101b in *Roq1* immunity. None of these EDS1-SAG101 combinations mediated *Roq1* restriction of *Xcv* bacterial growth at 6 dpi (Figure 3D) or XopQ-triggered *Roq1* cell death at 3 dpi (Figure 3E) in *Nb-epss*. All tagged proteins accumulated in these assays, as measured on Western blots at 2 dpi (Supplemental Figure 3B). Hence, *At*EDS1 and *At*SAG101, as a homologous pair or together with functional *Nb*SAG101b and *Sl*EDS1, are not functional in *Roq1* signaling. We concluded that the Arabidopsis EDS1-SAG101 heterodimer is inactive or insufficient for signaling in *TNL Roq1* immunity in *N. benthamiana*.

### *At*EDS1 and *At*SAG101 with *At*NRG1.1 or *At*NRG1.2 rescue XopQ-triggered cell death in *Nb-epss*

*SAG101* and *NRG1* were reported to be absent from monocots and several dicot species (*Aquilegia coerulea, Erythranthe guttata)* (Collier et al., 2011; Wagner et al., 2013). Because we additionally did not find *SAG101* in conifers and *Caryophyllales* (Supplemental Table 1), we searched for *NRG1* in these species. Manual reciprocal BLAST searches in nine genomes and transcriptomes of *Caryophyllales* (six *Silene* species, spinach, amaranth and quinoa) failed to identify NRG1 orthologs. Similarly, an OrthoMCL-derived NRG1 orthogroup did not contain conifer sequences, whereas ADR1 orthologs were detected in the examined conifer and *Caryophyllales* species (Supplemental Table 1). The strong *SAG101* and *NRG1* co-occurrence signature combined with *Roq1* dependency on *Nb*SAG101b (Figure 3) and *Nb*NRG1 (Qi et al., 2018) in *N. benthamiana* ETI, prompted us to test whether *At*NRG1.1 or *At*NRG1.2 expressed with *At*EDS1-*At*SAG101 confer *Xcv* resistance and/or XopQ-triggered cell death in *N. benthamiana*.

Previously, tagged *Nb*NRG1, *At*NRG1.1 and *At*NRG1.2 forms or their corresponding CC domains were shown to elicit cell death upon agroinfiltration of *N. benthamiana* leaves (Peart et al., 2005; Collier et al., 2011; Wróblewski et al., 2018; Wu et al., 2018b). Using the quantitative ion leakage assay, we tested whether transiently expressed *At*NRG1.1 or *At*NRG1.2 controlled by a *35S* promoter and either untagged or fused N-or C-terminally to a StrepII-HA (SH) or eGFP epitope tag induce cell death in WT and *eds1a pad4 N. benthamiana* leaves (Supplemental Figure 4). N-and C-terminally eGFP-tagged *At*NRG1.2 produced a strong, and SH-tagged *At*NRG1.2 – a weak, cell death response in both backgrounds at 3 dpi (Supplemental Figure 4A). By contrast, N-and C-terminally eGFP-or SH-tagged *At*NRG1.1, as well as non-tagged *At*NRG1.1 or *At*NRG1.2 forms, did not induce cell death in these two *N. benthamiana* genotypes (Supplemental Figure 4A). Western blot analysis of the expressed proteins at 2 dpi showed that *At*NRG1.1-eGFP and *At*NRG1.2-eGFP accumulated to similar levels as YFP in both backgrounds (Supplemental Figure 4B). All eGFP-tagged *At*NRG1.1 and *At*NRG1.2 forms were detected in the cytoplasm (Supplemental Figure 4C). The data suggest that tagged *At*NRG1.2 but not *At*NRG1.1 induce cell death independently of *Nb*EDS1a, *Nb*PAD4 and XopQ activation of TNL Roq1 in *N. benthamiana*. To avoid possible *At*NRG1 autonomous cell death activity, we used the *At*NRG1.1-SH variant in subsequent TNL *Roq1* immunity assays, because it was clearly detectable (as two bands) on a Western blot (Supplemental Figure 4B) and did not elicit cell death in TNL non-triggered *N. benthamiana* leaves, similar to the untagged *At*NRG1.1 and *At*NRG1.2 proteins (Supplemental Figure 4A).

Agrobacteria-mediated transient expression of *At*NRG1.1, *At*NRG1.2 or *At*NRG1.1-SH together with *At*EDS1-YFP and *At*SAG101-SH (Figure 4, Supplemental Figure 5) produced cell death in *Nb-epss* leaves that was as strong as the WT *N. benthamiana* response to XopQ-myc infiltration (Figure 4A). Without XopQ-myc, none of the three *At*EDS1-*At*SAG101-*At*NRG1 combinations produced ion leakage above the negative control (YFP alone) at 3 dpi (Figure 4A), indicating that the cell death response is XopQ recognition-dependent. Western blot analysis at 2 dpi indicated that *At*NRG1.1-SH, *At*EDS1-YFP, *At*SAG101-SH proteins accumulated to similar levels in XopQ-myc treated and non-treated leaf extracts (Supplemental Figure 5A). These data show that *At*EDS1 and *At*SAG101 coexpressed with either *At*NRG1.1 or *At*NRG1.2 can restore XopQ/*Roq1*-triggered cell death in *Nb-epss* leaves. When *At*SAG101-SH was substituted by SH-*At*PAD4 in the assays (Supplemental Figure 5B), this did not restore *Roq1* cell death (Figure 4B). We concluded that *At*EDS1-*At*SAG101-*At*NRG1 but not *At*EDS1-*At*PAD4-*At*NRG1 reconstitute a TNL cell death signal transduction module in this *Solanaceae* species. Because *At*EDS1 and *At*SAG101 failed to function with endogenous *Nb*NRG1 in triggering *Roq1* cell death (Figure 3E), there appears to be a requirement for molecular compatibility between these immunity components within species or clades.

**Figure 4.**
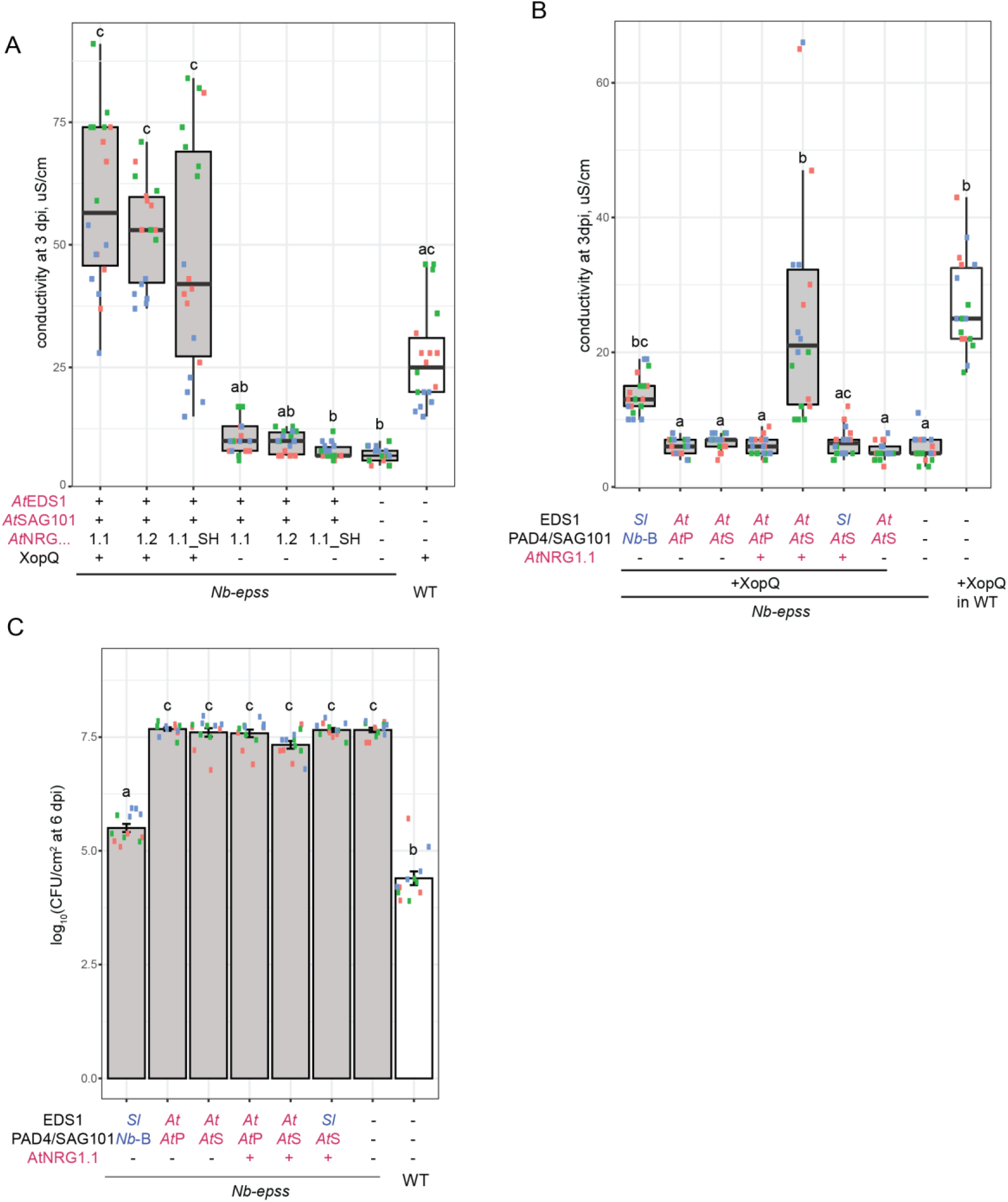
An AfEDS1-AfSAG101-AfNRG1 module rescues *Roq1*-dependent cell death but not resistance to Xcv in *N. benthamiana*. (A) Ion leakage in *N. benthamiana* WT and *eds1a pad4 sag 101a sag101b* (*Nb-epss)* plants transiently expressing combinations of Arabidopsis EDS1-YFP, SAG1D1-SH. ncn-tagged or C-terminally SH-tagged NRG1.1 or NRG1.2 protons in the prosonco of *Xcv* effector XopQ-myc (“-”in the sample description refers to YFP). In box-plots, ion leakage is dGtected m WT pants (white cox) at 3 dpi A XopQ dependent increase in conductivity was observed in *Nb-epss* samples expressing AfEDSI and AJSAG101 with A*t*NRG1.1 or A*t*NRG1 2. The experiment was repeated three times (dots of the same color represent six technical replicates (leaf discs) from cne independent experiment (biological replicate)) Shared letters above the box-whiskers between samples indicate that differences are not statistically significant (using a Nemenyi test with Bonferroni correction for multiple testing. α=0.01, n=18). (B) Ion leakage in *N benthamiana* WT (white) and *Nb-epss* (gray) plants expressing combinations of A*t*EDS1-FLAG with Strepll-HA-A*t*PAD4 (AfP) or A*t*SAG101-SH (A*t*S) and A*t*NRG1 1-SH as indicated. and measured at 3 “-” dpi in sample descriptions indicates YFP Expression of XopQ-myc in WT or FLAG-S,/EDS1(S/)WbSAG101b-GFP *(Nb-B)* in *Nb-epss* leads serve as controls SH-A/PAD4 cannot substitute for A/SAG 101-SH in the reconstitution assay. The expenment was repeated ihree times independently (dots of the same color represent six technical replicates (leaf discs) from cne independent experiment (biological replicato)). Statistical analysis was porformcd with a Nemenyi tost, and row p values were Bonferroni corrected for multiple testing (a-0.01. n_18). C) Complementation of *Nb-vpss* (gray) susceptibility to Xcv growlh by transiently expressing combinations of FLAG-SJEDS1 (SO, AfEDS1-YFP *{At),* Wt>SAG101b-GFP *(Nb-Q),* AfSAGIOI-SH SH-AfPAD4 and AJNRG1 1-SH (as in the panel B). Expression of FLAG SJEDS1/WbSAG101b-GFP partially restores resistance to *Xcv* compared to WT (white) whereas no combination with Arab dopsis protoins reduced Xcvqrowth. The expenment was repeated three times independently (dots of the same color represent four technical replicates (extractions of bactena) within one independent expenment (biological replicate)). Statistical analysis of Xcv titers at 6 dpi used a Tukey’s HSD test showing significant differences between means (a=0.001, N=12)in samples with different letter codes above bars Error bars represent iSEM.

Strikingly, neither the *At*EDS1-*At*SAG101-*At*NRG1.1 nor the *At*EDS1-*At*PAD4-*At*NRG1.1 combination restored *Roq1* resistance to *Xcv* bacterial growth in *Nb-epss* infection assays at 6 dpi, in contrast to the *Sl*EDS1-*Nb*SAG101b pair (Figure 4C).Therefore, the *At*EDS1-*At*SAG101-*At*NRG1.1 cell death module identified here lacks the capacity to limit *Xcv* growth in *N. benthamiana*. We also concluded that native *Nb*EDS1a-*Nb*PAD4 or the trans-clade *At*EDS1-*At*PAD4 pairs do not contribute to *Roq1* restriction of bacteria in these *N. benthamiana* assays (Figure 3B and C, Figure 4C). This contrasts with important roles of *Sl*EDS1-*Sl*PAD4 and *At*EDS1-*At*PAD4 partners in Arabidopsis TNL (*RPP4*) immunity (Figure 2).

### An EP domain α-helical coil surface on *At*SAG101 confers *Roq1* cell death

The above data suggest that the *At*EDS1-*At*SAG101 heterodimer has a distinctive feature that is not shared by *At*EDS1-*At*PAD4 ((Wagner et al., 2013), Supplemental Figure 1B), which enables cooperation with *At*NRG1.1 in XopQ/*Roq1*-dependent cell death in *Nb-epss* plants (Figure 4A-C). Several *At*EDS1 EP-domain residues lining a cavity formed by the heterodimer are essential for *At*EDS1-*At*PAD4 TNL immunity signaling in Arabidopsis (Bhandari et al., 2019). Here, examining EDS1-PAD4 and EDS1-SAG101 evolutionary rate variation across seed plants (Supplemental Figure 1B) highlighted conserved residues on a prominent α-helical coil of *At*PAD4 and *At*SAG101 that spans the length of the EP-domain cavity and, at its base, creates contacts with the *At*EDS1 EP domain ((Wagner et al., 2013), Supplemental Figure 1B). We therefore generated four *At*PAD4-*At*SAG101 chimeric proteins (1 to 4) with decreasing *At*SAG101 contributions to this central EP domain α-helical coil (Figure 5, schematic for chimeras in Figure 5A, SAG101 shown in pink). All four *At*PAD4-*At*SAG101 chimeras contained the complete *At*PAD4 N-terminal lipase-like domain (Figure 5A, green). In *Nb-epss* cell death assays, chimeras 1 to 4 fused N-terminally to a StrepII-YFP tag exhibited a nucleocytoplasmic localization, like YFP-*At*PAD4 (Supplemental Figure 6). We tested the chimeras in the *Nb-epss* TNL *Roq1* cell death reconstitution assay, as before with co-expressed *At*EDS1-YFP, *At*NRG1.1-SH and XopQ-myc (Figure 5B and 5C). Chimeras 1 and 2 mediated XopQ-dependent cell death whereas chimeras 3 and 4 were inactive in quantitative ion leakage assays and macroscopically at 3 dpi (Figure 5B and 5C). All chimeras accumulated to similar levels but less well than YFP-*At*PAD4 or *At*SAG101-YFP full-length proteins at 2 dpi (Supplemental Figure 5C). Comparing the sequences of functional and non-functional chimeras 2 and 3 allowed us to delineate an *At*SAG101 α-helical coil patch responsible for cell death reconstitution to between amino acids 289-308 (Supplemental Figure 5D). These results suggest that a discrete region of the *At*SAG101 EP domain is necessary for conferring *Roq1*-dependent cell death.

**Figure 5.**
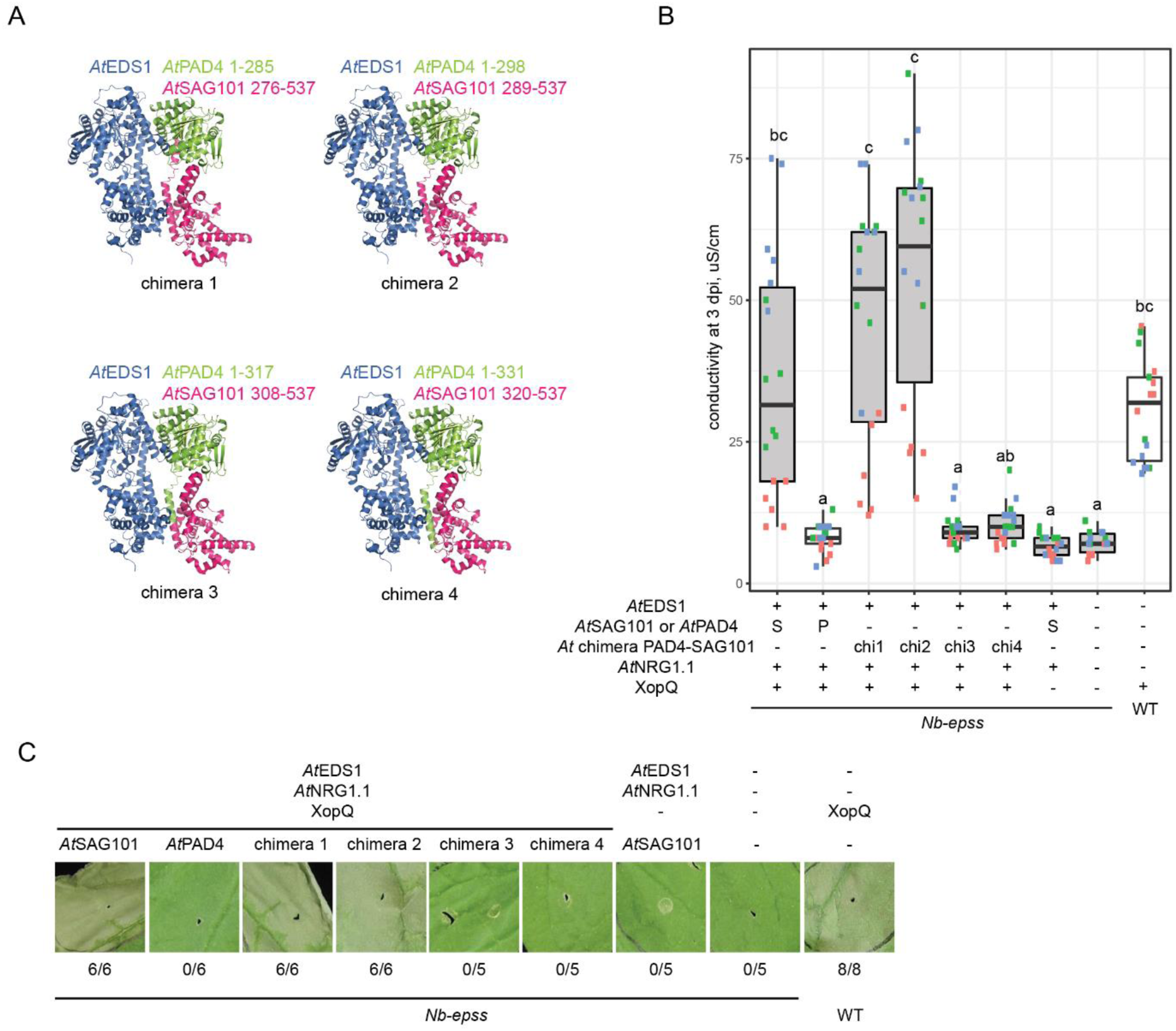
Differences in the EP domains of ArSAGlOl and /UPAD4 determine functionality TNL *Roql* cell death In *N. benthamiana.* (A) Schematic representation of A*t*PAD4-A*t*SAG101 chimeras used in assays shown in panels B and C. The /\fEDS1-A*t*SAG101 crystal structure (PDB ID 4nfu) is used as background with A*t*PAD4 or A*t*SAGIOI portions and amino add positions shown in green and pink, respectively A*t*EDSI is colored blue (B) Ion leakage assay quantifying XopQ-myc triggered cell death in eds*1*a *pad4 sag101a sag101b (Nb-epss)* plants (gray) expressing A*t*PAD4, A*t*SAG101 or chimeras (chi1 to chi4. as indicated) with A*t*EDSI. A*t*NRGI 1 and XopQ Cell death in WT (white) in response to XopQ served as a control The expenment was performed three times (dots of the same color represent six technical replicates (leaf discs) from one independent experiment (biological replicate)) Statistical analysis was performed using a Nemenyi test with Bonferroni correction for multiple testing Samples with different letters above the box-plots have statistically significant differences in conductivity (a=0 (11. n=18) (C) Macroscopic cell death symptoms in WT and *Nb-epss* leaf panels at 3 d after agroinfiltration of the protein combinations shown in panel B In contrast to the ion leakage assays, infiltrated Ieaves were wrapped in aluminum foil for 2 d and photographs taken at 3 dpi. Numbers under each image indicate necrotic/total infiltrated sites observed in three independent experiments. In (B) and (C), “-”in the infiltration scheme refers to addition of YFP expressing strain of *A tumefaciens.*

### TNL (*Roq1*) cell death in *N. benthamiana* requires the *At*EDS1 EP domain

Next we tested whether the *At*EDS1 EP domain is necessary for XopQ-triggered cell death in *Nb-epss* (Figure 6, Supplemental Figure 7). Because the EDS1 EP domain is unstable without its N-terminal lipase-like domain (Wagner et al., 2013), we compared activities of full-length *At*EDS1-FLAG and the FLAG-*At*EDS1 lipase-like domain (amino acids 1-384, (Wagner et al., 2013)), which accumulated to similar levels in *Nb-epss* leaves (Supplemental Figure 7A). The *At*EDS1 lipase-like domain did not confer XopQ-triggered cell death (Figures 6A and 6B), indicating there is a requirement for the *At*EDS1 EP domain in reconstituting *N. benthamiana* TNL (*Roq1*) cell death. We next tested effects of individually mutating two *At*EDS1 EP domain amino acids F419E and H476F which are on the *At*EDS1 EP domain α-helical coil surface closest to the *At*SAG101 patch found to be necessary for *Roq1* cell death (Figure 6C). Mutations at the *Sl*EDS1 position F435, which corresponds to *At*EDS1 F419, impaired *Sl*EDS1 function in *Roq1* cell death (Gantner et al., 2019). Alongside the two *At*EDS1 mutants, we tested two *At*EDS1 variants that are non-functional in Arabidopsis TNL immunity: *At*EDS1^LLIF^with weak EDS1-partner N-terminal binding (Figure 1C, 6C, (Wagner et al., 2013; Cui et al., 2018)), and *At*EDS1^R493A^with impaired EDS1-PAD4 heterodimer signaling (Bhandari et al., 2019). In the *Nb-epss* assays, *At*EDS1^F419E^and *At*EDS1^H476F^failed to confer *Roq1* cell death at 3 dpi whereas *At*EDS1^LLIF^and *At*EDS1^R493A^were functional (Figure 6D). The C-terminally YFP-tagged variants accumulated to similar or higher levels than WT *At*EDS1-YFP in *Nb-epss* leaves at 2 dpi (Supplemental Figure 7B). Put together with the *At*PAD4-*At*SAG101 chimera phenotypes (Figure 5), these data identify aligned parts of the *At*EDS1 and *At*SAG101 EP domains as being necessary for TNL-triggered cell death in *N. benthamiana*. Interestingly, the N-terminal ‘LLIF’ contact and EP domain R493 that are required for *At*EDS1-*At*PAD4 basal and TNL immunity in Arabidopsis (Wagner et al., 2013; Cui et al., 2018; Bhandari et al., 2019) are dispensable for *At*EDS1-*At*SAG101 cooperation with *At*NRG1.1 in the *Nb-epss* TNL (*Roq1*) cell death response.

**Figure 6.**
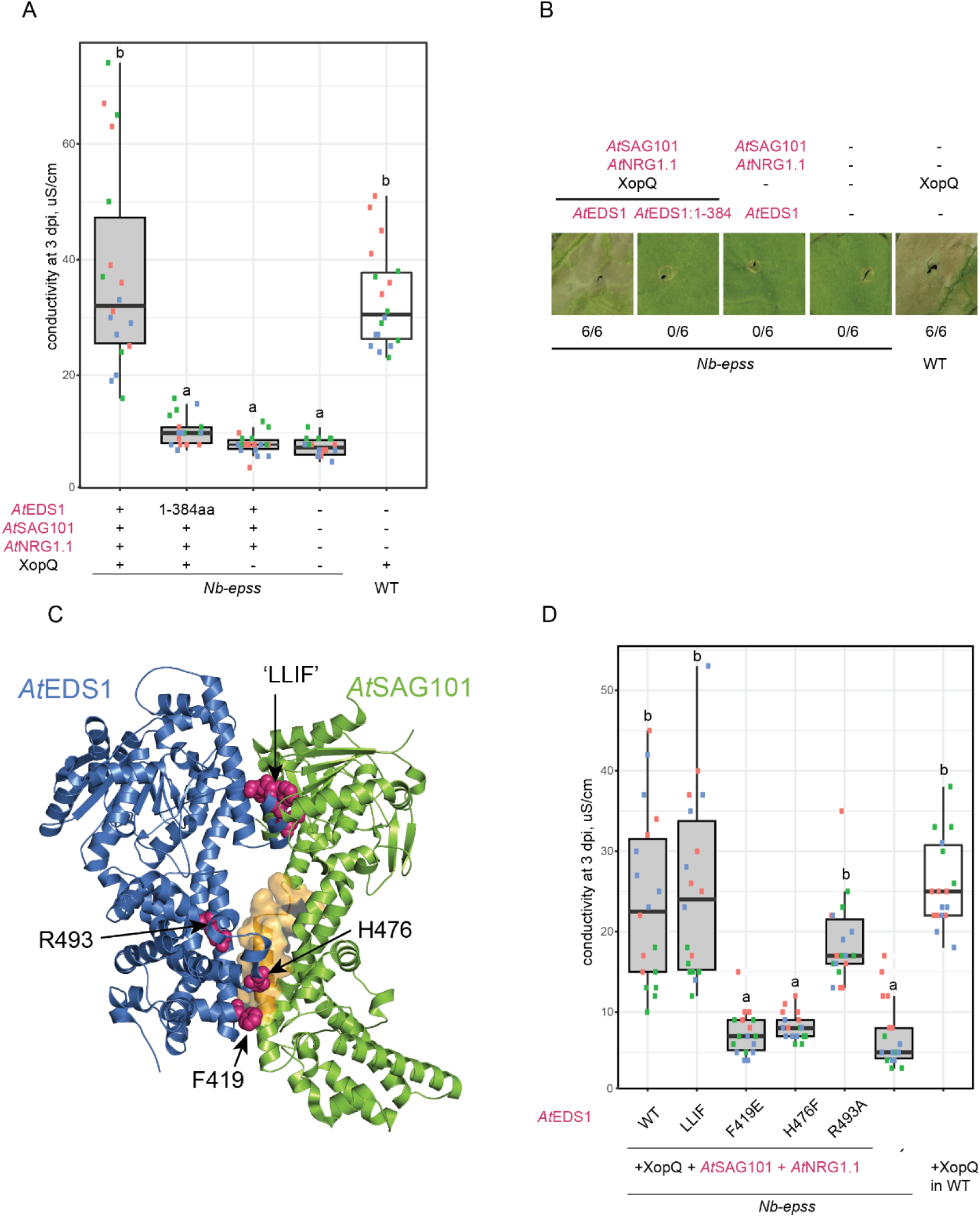
AtEDSI EP domain is essential for N. benthamiana Roq1 cell death. (A) Ion leakage assay for quantifying cell death in *N benthamiana* WT (white) and *eds1a pad4 sag 101 a sag 101b (Nb-epss)* (gray) after infiltration of XopQ-myc with A*t*EDSI-FLAG (A*t*EDSI), FLAG-/VEDS1 lipase-like domain (A(EDS 1:1-384). A*t*SAG101-YFP (A*t*SAG101). AtNRG1.1-SH (A*t*NRG1 1)or YFP (“-w in the sample descriptions). Only full-length A/EDS1-FLAG with AfSAG101-YFP and A/NRG 1 1-SH produces conductivity similar to WT plants infiltrated with XopQ. The experiment was performed three times (dots of the same color represent six technical replicates (leaf discs) in each experiment (biological replicate)). Nemenyi test was applied to test for significance of differences in conductivity (Bonferroni correction for multiple testing. a=0.01 for grouping samples, n=18). (B) Macroscopic cell death symptoms for combinations used in the ion leakage assays in (A). Numbers under each image indicate necroticftotal infiltrated sites observed in three independent experiments (C) AEDS1 ammo acids mutated in the strudure-funclion analysis of A*t*EDSI activity in the cell death reconstitution assay. A*t*EDSI and A/SAG101 are shown as blue and green ribbon diagrams respecbvely Ribbon and sphere depiction of the A/EDS1-A/SAG101 heterodimer crystal structure. Amino acids mutated in this analysis CLLIF’, R493, H476, F419) are displayed as pink spheres. The portion of an AfSAG101 EP domain central a-he(ical coil identified as essential for cell death activity in *Nb-epss* reconstitution assays with chimenc AfPAD4-AfSAG101 proteins is represented as an orange surface. (D) Ion leakage assay quantifying defects of A*t*EDSI-YFP mutants shown in (C) in *Nb-epss* (gray) reconstituted cell death compared to WT *N. benttmriiana* (WT, white) responding to XopQ Co-expression of A/EDS1-YFP (WT). A/EDS 1_R493A-YFP (R493A) or A*t*EDS1-lLIFfAAAA-YFP (LLIF) with A*t*SAG101-SH and AfNRGl 1-SH produced XopQ-myc dependent ion leakage, whereas mutated A/EDS1-YFP (H476F and F416E) were similar to the negative YFP (-) control. Experiment were performed three times independently (dots of the same color represent six technical replicates (leaf discs) from one independent experiment (biological replicate)). Samples with different letters above the box-plots have statistically significant differences in conductivity (o=0.01) after Nemenyi test followed by Bonferroni correction for multiple testing (n=18)

### EDS1 EP domain mutants are impaired in Arabidopsis *RRS1S-RPS4* cell death

The above analysis of the *At*EDS1^F419E^and *At*EDS1^H476F^mutants revealed importance of these EP domain residues with *At*SAG101 and *At*NRG1.1 in *N. benthamiana* TNL cell death (Figure 6D). We examined, whether the same mutations affect Col TNL (*RRS1S-RPS4*) immunity to *Pseudomonas syringae* pv. *tomato* DC3000 (*Pst*) *avrRps4* in Arabidopsis (Figure 7). For this, genomic *At*EDS1^F419E^and *At*EDS1^H476F^(gEDS1-YFP) constructs were transformed into Col *eds1-2* and two independent homozygous transgenic lines expressing EDS1-YFP proteins (#1 and #2) selected for each variant (Figure 7A). The lines were infiltrated with *Pst avrRps4* alongside WT Col, *eds1-2* and functional gEDS1-YFP or signaling defective *At*EDS1^R493A^and *At*EDS1^LLIF^controls (Wagner et al., 2013; Cui et al., 2018; Bhandari et al., 2019). As expected, *At*EDS1^R493A^and AtEDS1^LLIF^plants failed to restrict *Pst avrRps4* growth (Figure 7B). *At*EDS1^F419E^was also fully susceptible but, surprisingly, *At*EDS1^H476F^retained *RRS1S-RPS4* resistance (Figure 7B). We tested the same lines for TNL (*RRS1S-RPS4*) macroscopic cell death at 24 hpi after infiltration of *Pseudomonas fluorescens* 0-1 (*Pf*0-1) delivering AvrRps4 (Heidrich et al., 2011; Sohn et al., 2014). Here, all variants (*At*EDS1^F419E^, *At*EDS1^H476F^, *At*EDS1^R493A^and *At*EDS1^LLIF^) were defective in cell death (Figure 7C). Therefore, *At*EDS1^LLIF^, *At*EDS1^R493A^and *At*EDS1^F419E^failed to limit bacterial growth and induce TNL (*RRS1S-RPS4*) cell death in Arabidopsis, whereas *At*EDS1^LLIF^and *At*EDS1^R493A^but not *At*EDS1^F419E^retain cell death-inducing activity in *N. benthamiana*. More strikingly, *At*EDS1^H476F^was defective in cell death in Arabidopsis and *N. benthamiana* but fully competent in Arabidopsis TNL resistance to bacteria. Together, these data suggest that the same *At*EDS1 EP domain surface lining the *At*EDS1-*At*PAD4 or *At*EDS1-*At*SAG101 cavity controls bacterial TNL resistance and host cell death in Arabidopsis and *N. benthamiana*, but that EDS1-SAG101 and EDS1-PAD4 EP domain signaling functions are different.

**Figure 7.**
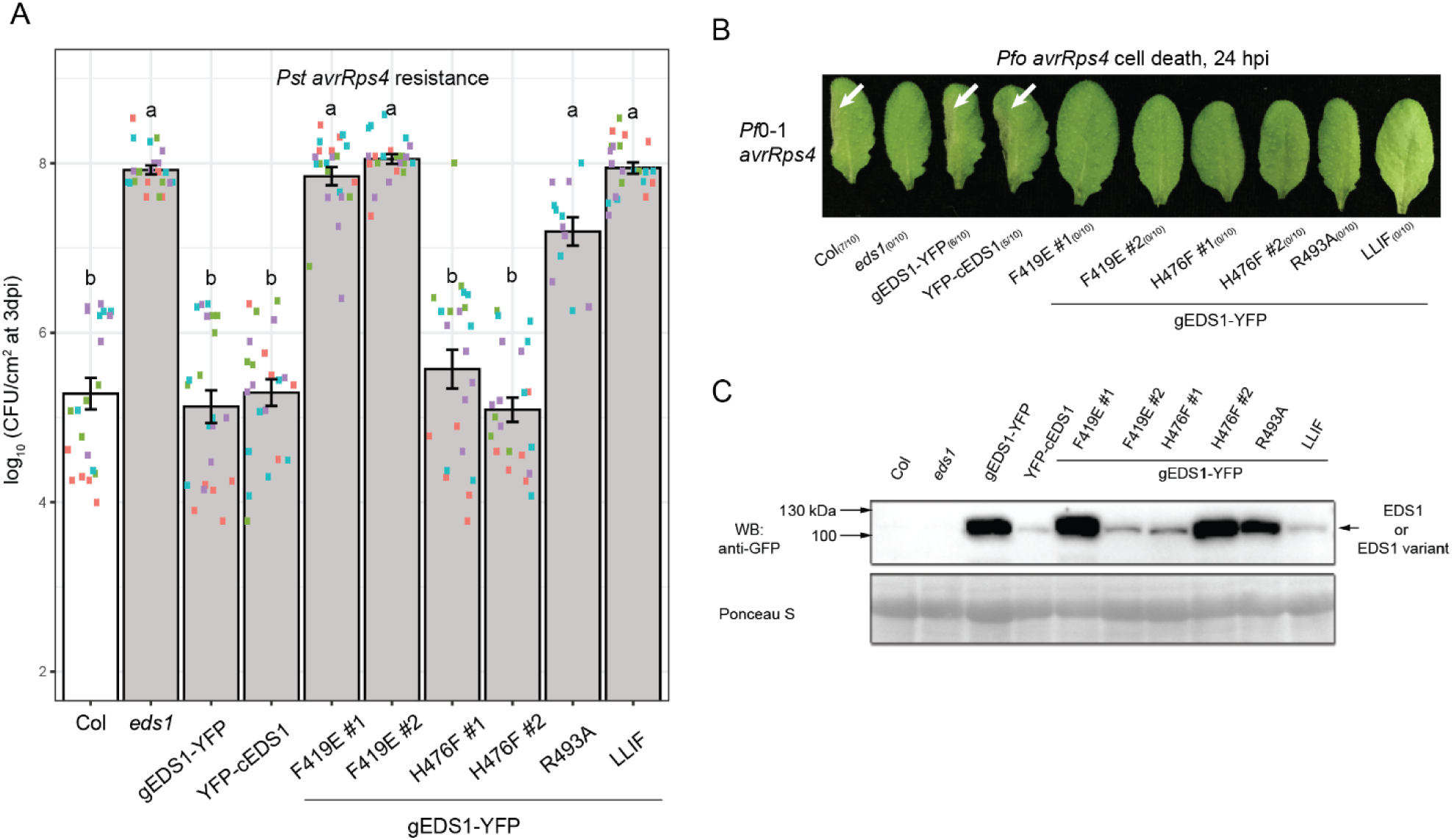
A*t*EDSI variants with mutated lipase-like and EP domains have different defects in Arabidopsis *RRS1S-RPS4* cell death and bacterial growth arrest. (A) *Pst avrRps4* titers at 3 dpi (OD600=0.C005) in leaves of Arabidopsis Col *eds1-2* (gray) independent homozygous transgenic lines expressing Arabidopsis gEDS1-YFP and corresponding mutant variants (F419E, H476F. R493A, ‘LLIF’) under control of native *pEDS1* promoter, as indicated. Responses in WT coding (cEDS1) and genomic (gEDS1) ArEDS1 transgenic (gEDS1-YFP and YFP-cEDS1) lines and Col (white) served as controls Experiments were performed four times (biological replicates) (and twice for the R493A line) each with five technical replicates (extractions of bacteria). Statistical analysis used a Tukey’s HSD test and grouping of genotypes by letters at the significance threshold α=0.001 (n=10-20). Error bars represent ±SEM. (B) Macroscopic cell death of Arabidopsis leaves of the same genotypes as used in (A), visible as tissue collapse at 24 h after *Pf*0-1 *avrRps4* infiltration (OD600=0.2). Experiments were repeated three times with similar results. Numbers in parentheses indicate leaves showing visual tissue collapse/infiltrated leaves in one representative experiment. None of the tested A*t*EDS1-YFP mutant variants developed cell death symptoms observed in Col and cEDS1 or gEDS1 transgenic lines. (C) Western blot analysis of Arabidopsis lines expressing the YFP-tagged WT EDS1 and mutant variants tested in the (A) and (B), prior to pathogen infiltration. The analysis was performed twice independently with similar results Ponceau S staining of membrane shows equal protein loading.

### Different genetic requirements for cell death and bacterial growth restriction in Arabidopsis RRS1/RPS4 immunity

Evidence of *At*EDS1-*At*SAG101-*At*NRG1 activity and *At*EDS1-*At*PAD4-*At*NRG1 inactivity in *N. benthamiana* TNL-triggered cell death (Figure 4, Figure 5) lends support to engagement of distinct *At*SAG101/*At*NRG1 and *At*PAD4/*At*ADR1 immunity branches in Arabidopsis TNL signaling, as proposed by (Wu et al., 2018b).

To test this, we quantified TNL (*RRS1S-RPS4*) bacterial resistance and cell death phenotypes in Arabidopsis Col *EDS1*-family single mutants (*eds1-2, pad4-1* and *sag101-3*), a double mutant *pad4-1 sag101-3* and AvrRps4 non-recognizing mutant *rrs1a rrs1b* (Saucet et al., 2015), together with a CRISPR-associated 9 (Cas9) generated Col *AtNRG1.1 AtNRG1.2* double mutant line *nrg1.1 nrg1.2* (denoted *n2*) (Figure 8, Supplemental Figure 8; mutations introduced in *n2* shown in Supplemental Figure 8A). Different Arabidopsis TNLs exhibited varying genetic dependencies on *NRG1*-and *ADR1*-family genes (Castel et al., 2018; Wu et al., 2018b). Also, NRG1 and ADR1 orthologs share a phylogenetically distinct nucleotide-binding domains (Supplemental Figure 8B, (Collier et al., 2011; Shao et al., 2016)). We therefore also generated a pentuple *nrg1.1 nrg1.2 adr1 adr1-L1 adr1-L2* mutant (denoted *n2a3*) by transforming the *AtNRG1.1/AtNRG1.2* Cas9 mutagenesis construct into an *adr1 adr1-L1 adr1-L2* triple mutant (*a3*) (Supplemental Figure 8A, (Bonardi et al., 2011)).

**Figure 8.**
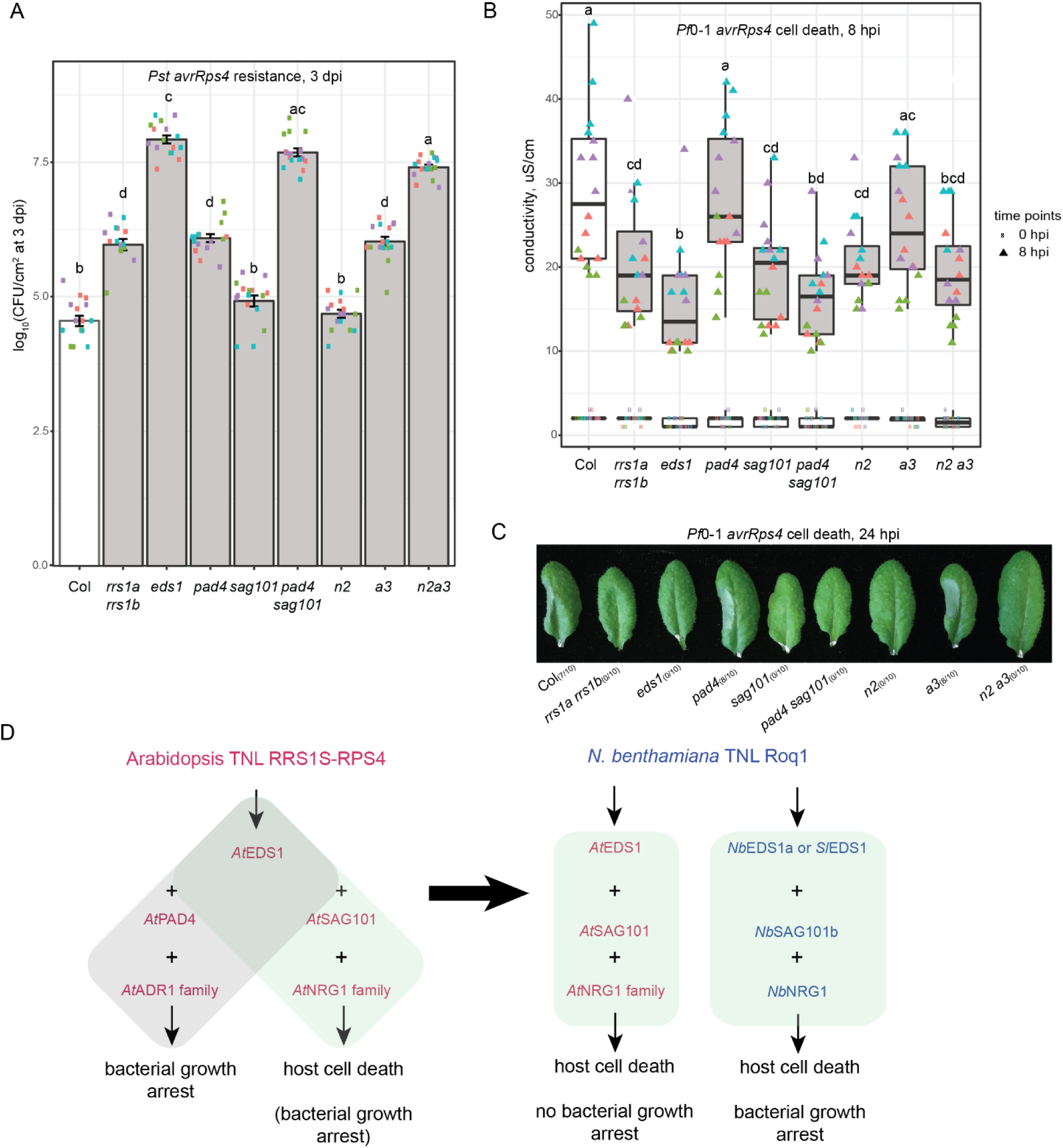
Differential requirements for *EDS1-, NRG1-* and *ADR1*-family genes in Arabidopsis *RRS1S-RPS4* cell death and bacterial growth arrest. (A) *Psl avrRps4* titers at 3 dpi (00600=0.0005) in leaves of Arabidopsis Col (white) and single or combinatorial mutants lines, as indicated. Experiments were performed four times independently (dots with the same colors represent technical replicates (extractions of bactena) within the same experiment (biological replicate)). Letters above bars correspond to statistical grouping after a Tukey’s HSD test (a=0.001. n=16). Error bars represent ±SEM. (B) Ion leakage assay quantifying P/0-1 *avrRps4* (OD600=0.2) cell death (at 8 hpi) in the Arabidopsis genotypes tested in (A). The experiment was repeated four times independently (dots with the same color represent four technical replicates (leaf discs) in one experiment (biological replicate)). Samples with overlapping letter codes above the whisker-boxes do not show statistically significant differences (Tukey’s HSD. a=0.0D1, n=16). (C) Macroscopic cell death symptoms, visible as tissue collapse at 24 h after infiltration of P/0-1 *avrRps4* (OD600=0.2) into leaves of the Arabidopsis lines tested in (A). Numbers in parentheses represent leaves showing tissue collapse/total infiltrated leaves in one experiment. One experiment is shown as representative of three independent experiments. (D) Schematic showing cooperation between EDS1-family proteins and ADR1 or NRG1 helper NLRs in Arabidopsis and *N benthamiana* TNL immune responses tested in this study In Arabidopsis. RRS1S-RPS4 recognition of AvrRps4 in ETI bolsters EDS1-PAD4 ADR1 immune responses leading to restriction of *Pst avrRps4.* A different RRS1S-RPS4 ETI pathway mediated by EDS1-SAG101-NRG1 promotes host cell death but these components are dispensable for limiting bacterial growth if the EDS1-PAD4-ADR1 branch is operational. A complete Arabidopsis TNL immune response requires cooperation between the tv/o branches. In *N. benthamiana,* TNL Roq1-conditioned bactenal (Xcv) growth arrest and cell death are channeled through the EDS1-SAG101-NRG1 signaling module and therefore do not require EDS1-PAD4. Cross-clade transfer of a compatible Arabidopsis EDS1-SAG1D1-NRG1 module is sufficient to signal Roq1 cell death but not resistance to *Xcv* growth. This recapitulates the cell death-promoting functions of AfSAG101. AfNRGl.1 and AfNRGl.2 and their rather weak contributions to bacterial restriction in Arabidopsis.

At the level of *Pst avrRps4* growth at 3 dpi, the *sag101* and *n2* mutants exhibited WT Col resistance (Figure 8A). The *pad4* and *a3* mutants partially restricted bacterial growth, phenocopying *rrs1a rrs1b*, whereas *eds1, pad4 sag101* and *n2a3* mutants were highly susceptible to *Pst avrRps4* (Figure 8A). These data show that *AtADR1*-family genes, like *At*PAD4 (Feys et al., 2005; Wagner et al., 2013), genetically compensate for loss of *AtSAG101* or *AtNRG1* functions in *RRS1S-RPS4* immunity and show that combined loss of the *ADR1-* and *NRG1-* helper NLR family functions, like loss of *PAD4* and *SAG101* together, produce a completely defective TNL/EDS1 bacterial immune response.

We measured TNL (*RRS1S-RPS4*) cell death phenotypes in the same panel of mutants after infiltration with *Pf*0-1 *avrRps4* bacteria by monitoring ion leakage at 8 hpi (Figure 8B) and macroscopic cell death at 24 hpi (Figure 8C). Host cell death was strongly reduced in *eds1* indicating it is *EDS1*-dependent (Heidrich et al., 2011; Sohn et al., 2014). The *pad4* and *a3* mutants exhibited a similar level of tissue collapse and ion leakage as WT Col (Figure 8B and 8C) and therefore are not essential components of TNL (*RRS1S-RPS4*) cell death. The *sag101* and *n2* mutants phenocopied *rrs1a rrs1b* with an intermediate cell death response (Figure 8B and 8C). The *pad4 sag101* double and *n2a3* pentuple mutants phenocopied *eds1-2* (Figures 8B and 8C), indicating complete loss of EDS1-dependent host cell death when combined activities of *AtNRG1*-with *AtADR1*-family, or *AtPAD4* with *AtSAG101* are lost. When compared with the *Pst avrRps4* growth phenotypes (Figure 8A), these demarcations between cell death-competent and cell death-compromised lines (Figure 8B and C) point to a major role of *At*NRG1.1 and *At*NRG1.2 proteins with SAG101 in promoting TNL/EDS1 cell death in Arabidopsis and that this is dispensable for limiting bacterial growth when PAD4 and ADR1-family functions are intact. By contrast, EDS1/PAD4, likely together with ADR1-family proteins, have a major role in limiting bacterial growth in Arabidopsis TNL (*RRS1S-RPS4*) immunity but are dispensable for host cell death.

## Discussion

In dicotyledonous species, activated intracellular TNL receptors converge on the non-NLR lipase-like protein EDS1, which transduces signals to downstream defense and cell death pathways to stop pathogen growth (Wiermer et al., 2005; Adlung et al., 2016; Qi et al., 2018; Gantner et al., 2019). In Arabidopsis, *At*EDS1 functions in a heterodimer with one of its partners, *At*PAD4, to transcriptionally mobilize anti-microbial defense pathways and bolster SA-dependent programs that are important for basal and systemic immunity (Rietz et al., 2011; Wagner et al., 2013; Cui et al., 2017; Bhandari et al., 2019). *At*EDS1-*At*PAD4 mediated signaling is sufficient for basal immunity to bacterial and oomycete pathogens and for ETI initiated by many TNL receptors (Wiermer et al., 2005). The function of *At*EDS1 heterodimers with its second partner, *At*SAG101, was not determined (Rietz et al., 2011; Xu et al., 2015). We show here that *At*EDS1 functions together with *At*SAG101 and *At*NRG1 helper CNL proteins as a coevolved host cell death signaling module in TNL (*RRS1S-RPS4*) ETI (Figure 8D). We provide genetic and molecular evidence that *At*EDS1-*At*SAG101-*At*NRG1 promote TNL-dependent cell death in their native Arabidopsis (Figure 8B and C) and in a solanaceous species, *N. benthamiana* (Figure 4A). In both systems, the cell death activity cannot be substituted by *At*EDS1 with *At*PAD4 (Figure 4B, Figure 5). We establish that *AtSAG101* and *AtNRG1.1/AtNRG1.2* also contribute to Arabidopsis TNL (*RRS1S-RPS4*) restriction of *Pst avrRps4* bacterial growth in the absence of *AtPAD4* and *AtADR1*-family (Figure 8A). By contrast, in an *N. benthamiana* TNL (*Roq1*) reconstitution assay, the *At*EDS1-*At*SAG101-*At*NRG1.1 module confers cell death (Figure 4B) but is inactive or insufficient for limiting *Xcv* bacterial growth (Figure 4C), suggesting there is a functional mismatch or incompatibility between these proteins and *N. benthamiana* immunity factors. Our analysis of EDS1-family evolutionary rate variation (Supplemental Figure 1B) coupled with resistance/cell death phenotyping of targeted *At*EDS1 and *At*SAG101 protein variants (Figure 3D and E, Figure 4 and 5) provide additional evidence that EDS1-SAG101 have coevolved with NRG1 to promote TNL cell death and a structural basis (Figures 5-7) for understanding functionally distinct EDS1-SAG101 and EDS1-PAD4 branches in TNL immunity signaling (Figure 8D).

A motivation for this study was to explore EDS1-family variation between different plant lineages in order to identify constraints that might influence protein functionality between distant clades. For this, we first performed a large-scale phylogenetic analysis of EDS1-family orthologs across 46 seed plant species (Figure 1A, Supplemental Figure 1A, Supplemental Table 1, Supplemental Datasets 5-7). This identified well-supported, phylogenetic groups for EDS1, PAD4 and SAG101 protein-coding sequences in *Brassicaceae* and *Solanaceae*, and for EDS1 and PAD4 in *Poaceae, Pinacea* (conifers) and *Caryophyllales*, which lack *SAG101* genes (Supplemental Table 1). This analysis places origins of the EDS1-family deeper in the evolutionary history of seed plants and not only angiosperms (Wagner et al., 2013). While *EDS1* and *PAD4* are present in the majority of seed plants, *SAG101* has experienced dynamic evolution via loss in flowering plants (Supplemental Table 1). It is unclear whether *SAG101* emerged only in flowering plants or existed earlier in a common ancestor of seed plants. Since *TNL* genes exist in seed plant species without *SAG101* and *NRG1*, as in conifers (Supplemental Table 1, (Meyers et al., 2002)) and in non-seed plants without an entire EDS1-family (Gao et al., 2018), it is possible that some TNLs signal without SAG101 and NRG1. Indeed, TNL Roq1 functioned in effector XopQ-dependent cell death in *Beta vulgaris* (Schultink et al., 2017) which does not have recognizable *SAG101* or *NRG1* genes (Supplemental Table 1). Identification of conserved regions in EDS1, PAD4 and SAG101 (Supplemental Figure 1B) close to the EP-domain interaction surfaces and at the ‘LLIF’ α-helix (Supplemental Figure 1C) promoting EDS1-family hetero-dimerization (Wagner et al., 2013) suggested molecular possibilities for physical interactions between proteins from different taxonomic groups. Testing of EDS1 partner interactions within and between the angiosperm families *Brassicaceae, Solanaceae* and *Poaceae* (Figure 1B, Supplemental Figure 2A) showed conserved within-species or-clade partner associations but certain barriers to EDS1 heterodimer formation between groups.

Two recent studies of TNL and CNL receptor signaling in Arabidopsis show that TNL receptors utilize genetically redundant *ADR1* (*ADR1, ADR1-L1* and *ADR1-L2*) and *NRG1* (*NRG1.1, NRG1.2*) helper NLR families to different extents for immunity (Castel et al., 2018; Wu et al., 2018b). These and earlier reports (Bonardi et al., 2011; Dong et al., 2016) provide evidence that Arabidopsis ADR1 and NRG1 proteins work as parallel branches downstream of TNL activation. Genetic data supported *AtADR1*s and *AtPAD4* operating in the same *EDS1*-controlled pathway to bolster SA and/or other transcriptional defenses, whereas *AtNRG1*s were important for promoting host cell death (Bonardi et al., 2011; Dong et al., 2016; Castel et al., 2018; Wu et al., 2018b). Several tested Arabidopsis TNLs recognizing oomycete pathogen strains, a TNL autoimmune allele of Suppressor of Npr1-1, Constitutive1 (SNC1) and a TNL pair CHS3/CSA1 displayed varying dependence on *AtNRG1* signaling in immunity (Castel et al., 2018; Wu et al., 2018b). By contrast, all so far tested TNL pathogen resistance and cell death responses in *N. benthamiana* signaled via *NbEDS1a, NbNRG1* and *NbSAG101b*, but did not require *NbPAD4* (Adlung et al., 2016; Qi et al., 2018; Gantner et al., 2019). Similarly, we find a dependency of TNL (*Roq1*) immunity and cell death responses to *Xcv* bacteria on *NbEDS1a* and *NbSAG101b* but not *NbPAD4* (Figure 3). Collectively, these data suggest that while there is TNL signaling pathway choice in Arabidopsis, a strong pathway preference exists in *N. benthamiana* for *EDS1* with *NRG1* and *SAG101*.

Further support for a TNL two-branched resistance signaling model in Figure 8D comes from quantifying Arabidopsis TNL (*RRS1S-RPS4*) *Pst avrRps4* growth and *Pf*0-1 *avrRps4* cell death phenotypes in *ADR1*-family triple (*a3*) (Bonardi et al., 2011), double *nrg1.1 nrg1.2* (*n2*) and a combined (pentuple) *n2a3* mutant, alongside *EDS1*-family mutants (Figure 8A-C). Importantly, effects of *pad4* and *sag101* single mutations on Arabidopsis *RRS1S-RPS4* resistance and cell death responses were, respectively, phenocopied by the *a3* (*adr1 triple*) and *n2* (*nrg1.1 nrg1.2*) mutants (Figure 8A-C). Proposed PAD4-ADR1 and SAG101-NRG1 co-functions in bacterial immunity (Figure 8D) is in line with Arabidopsis *NRG1*-like genes regulating *SAG101*-dependent *chs3-2d* autoimmunity (Xu et al., 2015; Wu et al., 2018b) and *ADR1*-like genes regulating *PAD4*-dependent *snc1* autoimmunity (Zhang et al., 2003; Dong et al., 2016). The Arabidopsis TNL (*RRS1S-RPS4*) phenotypic outputs measured here show that each branch contributes to a complete *EDS1*-dependent immune response (Figure 8A and 8B), pointing to synergistic activities and hence a degree of cross-talk between the two immunity arms, as proposed for Arabidopsis *snc1* autoimmunity (Wu et al., 2018b).

Our data further suggest that making a clean distinction between *AtEDS1/AtNRG1/AtSAG101*-controlled cell death and *AtADR1/AtPAD4*-mediated transcriptional promotion of ‘basal’ defenses is not justified, since *AtPAD4* and *AtADR1*s accounted for a small but measurable portion of the *EDS1*-dependent *RRS1S-RPS4* cell death (Figure 8B). *AtSAG101* and *AtNRG1* contributions to Arabidopsis transcriptional reprogramming are not known, but in *N. benthamiana*, TNL (*Roq1*) EDS1-dependent cell death and bacterial resistance were abolished by mutations in *SAG101b* (Figure 3, (Gantner et al., 2019)) and *NRG1* (Qi et al., 2018). Interestingly, *Roq1*-dependent transcriptional reprogramming was almost entirely dependent on *EDS1a* and largerly dependent on *NRG1* (Qi et al., 2018), suggesting that other minor *EDS1*-dependent pathways are at play in *Roq1* immunity. Since Roq1 mediates XopQ-triggered cell death in *Beta vulgaris* (Schultink et al., 2017) which does not have detectable *SAG101* or *NRG1* orthologs (Supplemental Table 1), this TNL might have a capacity to function via a non-SAG101/NRG1 branch. Therefore, it will be of interest to test whether PAD4 and ADR1 are responsible for a set of transcriptional outputs in *N. benthamiana* TNL responses.

It is significant that *SlEDS1*, although not contributing with *Sl*PAD4 or *Nb*PAD4 to TNL (*Roq1*)-triggered *Xcv* resistance or cell death in *N. benthamiana* (Figure 3, (Gantner et al., 2019)), is functional in TNL (*RPP4*) resistance against an oomycete pathogen (*Hpa* Emwa1), when transferred to Arabidopsis (Figure 2B). Thus, *Sl*EDS1-*Sl*PAD4 retains an immunity activity. We presume this function is required for some pathogen encounters in *Solanaceae* hosts and speculate that the *Sl*EDS1-*Sl*PAD4 resistance activity in Arabidopsis reflects a core basal immunity transcriptional reprogramming function that is sufficient for *RPP4* ETI (Cui et al., 2017; Bhandari et al., 2019). Because *At*EDS1-*At*PAD4 heterodimers utilize the same EP domain surface (involving R493) for bacterial resistance conferred by a TNL (*RRS1S-RPS4*) and a CNL receptor RPS2 (Bhandari et al., 2019), it is possible that the maintenance of *EDS1* and *PAD4* genes across seed plants (Supplemental Table 1) also reflects their usage by certain CNLs, which can be masked by compensatory defense pathways. Indeed, the SA immunity branch works in parallel to EDS1/PAD4 in Arabidopsis CNL (RPS2) immunity (Venugopal et al., 2009; Cui et al., 2017; Mine et al., 2018).

In contrast to *Sl*EDS1-*Sl*PAD4 transferable function to Arabidopsis TNL immunity (Figure 2B), the *At*EDS1-*At*SAG101 heterodimer was not active in *N. benthamiana* TNL (*Roq1*) cell death unless co-expressed with *At*NRG1.1 or *At*NRG1.2 (Figure 4A). Also, interaction between *Sl*EDS1 and *At*SAG101 (Supplemental Figure 2A) was insufficient to mediate *Roq1* signaling with otherwise functional *At*NRG1.1 or *Nb*NRG1 proteins (Figures 3D and E, 4B and C). Hence, between-clade barriers exist beyond heterodimer formation for *At*EDS1-*At*SAG101 and *At*EDS1-*At*PAD4 (Figure 1B and C, Supplemental Figure 2A). These findings highlight a requirement for matching Arabidopsis proteins to constitute a functional EDS1-SAG101-NRG1 signal transduction module. The data also point to coevolutionary constraints existing not only on variable NLR receptor complexes (Concepcion et al., 2018; Schultink et al., 2019) but also on more conserved immunity signaling nodes.

The EDS1 structure-guided analyses done here (Figures 5-7) and by (Gantner et al., 2019) show that a conserved EDS1 EP domain signaling surface is necessary for TNL cell death in Arabidopsis and *N. benthamiana*. The same EP domain surface is required for rapid mobilization of transcriptional defenses and restriction of bacterial growth in Arabidopsis *RRS1S-RPS4* ETI (Bhandari et al., 2019). Surprisingly, two *At*EDS1 variants, *At*EDS1^LLIF^and *At*EDS1^R493A^, that are defective in Arabidopsis *RRS1S-RPS4* bacterial resistance and cell death (Figure 7), were functional in the reconstituted *N. benthamiana Roq1* cell death assay (Figure 6D). These difference might be due to a requirement in Arabidopsis for *At*EDS1^LLIF^and *At*EDS1^R493A^(with *At*PAD4) to transcriptionally regulate their own expression and the expression of important immunity components (Cui et al., 2018; Bhandari et al., 2019). That transcriptional role is dispensed with in the *N. benthamiana Roq1* assay, because *At*EDS1, *At*SAG101 and *At*NRG1 proteins are transiently overexpressed. In the future, it will be interesting to examine whether failure of *At*PAD4 to signal in *N. benthamiana* TNL *Roq1* resistance or cell death (Figure 3D and E) is down to the transient assay used or to mismatches with other *N. benthamiana* immunity components. These data reinforce the notion that there are striking functional distinctions between *At*PAD4 and *At*SAG101 in TNL/EDS1 signaling, as indicated by the *At*PAD4–*At*SAG101 chimeras, which identify a specific portion of the *At*SAG101 EP domain conferring cell death activity in *N. benthamiana Roq1* responses (Figure 5).

With identification of an EP domain surface at the EDS1-SAG101 heterodimer cavity that is important for TNL-dependent cell death (Figures 5-7, (Gantner et al., 2019)), it is tempting to speculate that the Arabidopsis EDS1-SAG101 heterodimer forms a complex with *At*NRG1.1 or *At*NRG1.2 to transmit TNL receptor activation to cell death pathways. We have considered whether this model is supported by our data. First, mutation of the *At*EDS1 ‘LLIF’ H α-helix, which strongly reduces *At*EDS1-*At*SAG101 dimerization (Wagner et al., 2013; Cui et al., 2018), did not disable *Roq1* reconstituted cell death in *Nb-epss* plants (Figure 6D). Here, we cannot rule out that partial impairment of EDS1^LLIF^-SAG101 heterodimerization is compensated for by protein overexpression in the *N. benthamiana* transient assays. Second, the subcellular localizations of transiently expressed *At*SAG101 and *At*NRG1 proteins tested in our *N. benthamiana* assays show that *At*SAG101 is mainly nuclear (Supplemental Figure 2B), as observed in Arabidopsis upon transient expression (Feys et al., 2005). By contrast, N-and C-terminally GFP-tagged *At*NRG1.1 and *At*NRG1.2 isoforms are cytoplasmic (Supplemental Figure 5C). A cytoplasmic endomembrane accumulation pattern was also in Arabidopsis stable transgenic lines and *N. benthamiana* transient assays for functional *At*NRG1-mNeonGreen isoforms, which did not obviously change upon TNL activation (Wu et al., 2018b). These data are difficult to reconcile with direct interaction between *At*EDS1-*At*SAG101 and *At*NRG1 underlying co-functions, although there might exist a small overlapping pool of these proteins that confers a cell death activity. In this regard, it is notable that a cytoplasmic AvrRps4 pool was found to elicit *EDS1*-dependent cell death in Arabidopsis *RRS1S-RPS4* immunity (Heidrich et al., 2011). *Nb*NRG1 was reported to interact directly with *Nb*EDS1a (Qi et al., 2018), implying a molecular link between these immunity signaling components.

## Methods

### Plant materials and plant growth conditions

Arabidopsis (*Arabidopsis thaliana* L. Heynh.) Col-0 (Col) mutants *eds1-2, pad4-1, sag101-3, pad4-1 sag101-3, eds1-2 pad4-1 sag101-1* were described previously (Glazebrook et al., 1997; Feys et al., 2005; Bartsch et al., 2006; Wagner et al., 2013; Cui et al., 2018). The mutant *eds1-2 pad4-1 sag101-3* was selected from a segregating F2 population *eds1-2 pad4-1* x *sag101-3* (Cui et al., 2018). The Col *adr1 adr1-L1 adr1-L2* triple mutant was kindly provided by J. Dangl (Bonardi et al., 2011). *Nicotiana benthamiana* mutants *eds1a, eds1a pad4, pad4 sag101a sag101b* are described in (Ordon et al., 2017; Gantner et al., 2019). The quadruple *N. benthamiana eds1a pad4 sag101a sag101b* mutant was selected from a cross between *eds1a* and *pad4 sag101a sag101b* mutants (Gantner et al., 2019). Genotyping was performed with Phire polymerase (F124, Thermo Fisher Scientific) on DNA extracted with the sucrose or Edwards methods (Berendzen et al., 2005). Oligonucleotides for genotyping are provided in the Supplemental Table 3. Arabidopsis homozygous transgenic Col *eds1-2* lines expressing Col coding (c) and genomic (g) *AtEDS1* sequence (pEN *pAtEDS1:YFP-cAtEDS1*, pXCG *pAtEDS1:gAtEDS1-YFP, pAtEDS1:gAtEDS1^LLIF^-YFP, pAtEDS1:gAtEDS1^R493A^-YFP*) are described in (García et al., 2010; Wagner et al., 2013; Cui et al., 2018; Bhandari et al., 2019). Arabidopsis plants for bacterial infiltration assays were grown for 4-5 weeks under a 10 h light/14h dark regime at 22°C/20°C and ∼65% relative humidity. Arabidopsis plants were kept under the same conditions after infiltration. Prior assays, *N. benthamiana* plants were grown for 5-6 weeks under a 16 h light/8 h dark regime at ∼24°C.

### Vectors generation by Gateway cloning

Coding sequences of *EDS1* and *PAD4* with stop codons were amplified from cDNA potato (DM 1-3), barley (cv. Golden Promise) and *Brachypodium distachyon* (BD21-3). Arabidopsis Col genomic and coding *EDS1, PAD4* and *SAG101* sequences were cloned previously (Feys et al., 2005; García et al., 2010; Wagner et al., 2013).Sequences of *AtNRG1.1* (AT5G66900.1), extended *AtNRG1.1* (AT5G66900.2) and *AtNRG1.2* (AT5G66910) were PCR-amplified using genomic DNA of Col as a template from start to stop codons. PCR amplification for all cloning was performed with Phusion (F530, Thermo Fisher Scientific) or PrimeStar HS (R010A, Clontech) polymerases. All sequences were cloned into pENTR/D-TOPO (K240020, Thermo Fisher Scientific) and verified by Sanger sequencing. Sequences of oligonucleotides used for cloning are provided in Supplemental Table 3. Entry clones for *Sl*EDS1 and *Sl*PAD4 from cv. VF36 are described in (Gantner et al., 2019). *AtNRG1.1* and *AtNRG1.2* sequences without stop codons were obtained by site-directed mutagenesis of the pENTR/D-TOPO constructs with stop codons. Recombination into pB7GWF2.0 (Karimi et al., 2002), pB7FWG2.0 (Karimi et al., 2002), pDEST_GAD424 (Mitsuda et al., 2010), pDEST_BTM116 (Mitsuda et al., 2010), pXCSG-GW-StrepII-3xHA (Witte et al., 2004), pXCSG-GW-mYFP (with *At*NRG1.1_Stop and *At*NRG1.2_Stop to generate non-tagged expression constructs; (Witte et al., 2004)), pXCG-GW-3xFLAG, pXCG-GW-mYFP and pENSG-YFP (Witte et al., 2004) as well as custom pENpAtPAD4 StrepII-YFP (Supplemental Dataset 8 with sequence in.gbk format) was performed using LR Clonase II (11791100, Life technologies).

### Vector generation by Golden Gate cloning

Level 0 constructs for coding sequences of *SlEDS1, SlPAD4, AtEDS1, AtPAD4, NbSAG101b* and promoter sequences of *AtEDS1* and *AtPAD4* are described in (Gantner et al., 2018; Gantner et al., 2019). *HvEDS1* and *HvPAD4* from cv. Golden Promise were cloned into level 0 pICH41308. Synthesized (GeneArt, ThermoFisher Scientific) coding sequence of *NbSAG101a* was cloned into the level 0 vector pAGM1287. At level 1, Arabidopsis, tomato and barley *PAD4* coding sequences were cloned into pICH47811 (*pAtPAD4:YFP-xxPAD4-35S_term*), *EDS1* – pICH47802 (*pAtEDS1:3xFLAG-xxEDS1-35S_term*). For level 2 constructs in pAGM4673, the *PAD4* expression module was placed at position 1, *EDS1* – at position 2, and *pNos:BASTA^R^-Nos_term* (pICSL70005)-cassette at position 3. The *35S:NbSAG101a-GFP-35S_term* and *35S:NbSAG101b-GFP-35S_term* expression constructs were cloned into pICH47802. Backbones (pAGM1287, pICH41308, pICH47802, pICH47811), tags (pICSL30005, pICSL30004, pICSL50008) and 0.4 kb CaMV35S promoter (pICH51277) and terminator (pICH41414) modules as well as BASTA^R^expression cassette (pICSL70005) are from the Golden Gate cloning toolkit (Engler et al., 2014). Sequences of oligonucleotides used for cloning are provided in Supplemental Table 3.

### Site-directed mutagenesis and generation of *At*PAD4-*At*SAG101 chimeras

To substitute stop codons for alanine in pENTR/D-TOPO *AtNRG1.1*, pENTR/D-TOPO *AtNRG1.2* and to introduce F419E and H476F mutations in pENTR/D-TOPO *pAtEDS1:gAtEDS1* (García et al., 2010), QuikChange II Site-Directed mutagenesis protocol (#200555, Agilent) was used with hot start polymerases Phusion (F530, ThermoFisher Scientific) or Prime Star (R010A, Takara). pDONR207 *At*PAD4-*At*SAG101 chimeric sequences were generated by overlapping PCR with oligonucleotides in Supplemental Table 3 and LR recombined into pENSG-mYFP (Witte et al., 2004) or a modified pENSG-mYFP with a *CaMV 35S* promoter substituted for a 1,083 bp *AtPAD4* region upstream of start codon (Supplemental Dataset 8 with sequence “pENpAtPAD4 StrepII-mYFP-GW.gbk”). In the expression constructs, *At*PAD4-*At*SAG101 chimeras were N-terminally tagged: *35S:mYFP-chimera* for cell death reconstitution assays or *pAtPAD4:StrepII-mYFP-chimera* for localization assays.

### Yeast-two-hybrid (Y2H) assays

Coding sequences of Arabidopsis (*At*), tomato (*Sl*), potato (*St*), barley (*Hv*) and *Brachypodium distachyon* (*Bd*) *EDS1* and *PAD4* in pENTR/D-TOPO were LR-recombined into gateway-compatible pDEST_GAD424 (Gal4 AD domain) and pDEST_BTM116 (LexA BD domain) (Mitsuda et al., 2010), respectively. The yeast leucine (L), tryptophan (W) and histidine (H) auxotroph strain L40 was used. No 3-amino-1,2,4-triazole (3-AT) was added to SD selection plates without L, W and H. Yeast growth on selection plates –LW and –LWH was recorded at 3 d after transformation.

### Transient expression (agroinfiltration) assays in *N. benthamiana*

*N. benthamiana* plants for agroinfiltration assays were grown under long-day conditions (24°C) for 5-6 weeks. Expression constructs (pAGM4673 *pAtEDS1:3xFLAG-xxEDS1/pAtPAD4:YFP-xxPAD4* (xx stands for donor species *At, Sl* or *Hv*), pICH47811 *pAtPAD4:YFP-AtPAD4*, pICH47802 *pAtEDS1:3xFLAG-SlEDS1,* pICH47802 *35S:NbSAG101a-GFP,* pICH47802 *35S:NbSAG101b-GFP,* pXCG *pAtEDS1:gAtEDS1-YFP* WT or ‘LLIF’, R493A, H476F, F419E variants, pXCG *pAtEDS1:gAtEDS1-3xFLAG*, pXCG *pAtEDS1:gAtEDS1^LLIF^-3xFLAG*, pENS *35S:3xFLAG-cAtEDS1^1-384^, 35S:AtNRG1.1_stop* and *35S:AtNRG1.2_stop* without a tag in pXCSG-mYFP, pXCSG *35S*:*AtNRG1.1-SH,* pXCSG *35S:AtNRG1.2-SH*, pB7WGF2.0 *35S:GFP-AtNRG1.1*, pB7WGF2.0 *35S:GFP-AtNRG1.2*, pB7GWF2.0 *35S:AtNRG1.1-GFP*, pB7GWF2.0 *35S:AtNRG1.2-GFP*, pXCSG *35S:gAtSAG101-SH*, pXCSG *35S:gAtSAG101-YFP*, pICH47811 *pAtPAD4:YFP-AtPAD4*, pXCSG *35S:AtPAD4-YFP*, pENSG *35S:SH-AtPAD4*, pENSG *35S:mYFP-AtPAD4/AtSAG101* chimeras 1 to 4, pENS *pAtPAD4:StrepII-mYFP-AtPAD4/AtSAG101* chimeras 1 to 4, pAM-PAT *35S:YFP*) were electroporated into *Rhizobium radiobacter* (*Agrobacterium tumefaciens*) GV3101 pMP90RK or pMP90. Final OD_600_ for each strain was set to 0.2, each sample contained *A. tumefaciens* C58C1 pCH32 to express *35S:p19* (final OD_600_=0.2 as well). Before syringe infiltration, *A. tumefaciens* was incubated in induction buffer (10 mM MES pH5.6, 10 mM MgCl_2_, 150 nM acetosyringone) for 1-2 h in the dark at room temperature.

### Western blot analysis

To test accumulation of proteins in *N. benthamiana* transient expression assays, four 8 mm leaf discs each were harvested at 2 dpi, ground in liquid nitrogen to powder and boiled in 150 μl 2xLaemmli buffer for 10 min at 95°C. For Arabidopsis lines, four 8 mm leaf discs or 4-5 seedlings per sample were processed in the same manner. The proteins were resolved on 8 or 10% SDS-PAGE (1610156, Bio-Rad) and transferred using the wet transfer method onto a nitrocellulose membrane (10600001, GE Healthcare Life Sciences). For protein detection, primary antibodies (anti-GFP #2956 (Cell Signaling Technology) or 11814460001 (Roche), anti-HA #3724 (Cell Signaling Technology) or11867423001 (Roche), anti-FLAG F7425 (Sigma Aldrich) or F1804 (Sigma Aldrich), anti-myc #2278 (Cell Signaling Technologies)) were used in the dilution 1:5,000 (1xTBST, 3% milk powder, 0.01% NaAz). Secondary HRP-conjugated antibodies (A9044 and A6154 (Sigma Aldrich), sc-2006 and sc-2005 (Santa Cruz)) were applied in the dilution 1:5,000. Detection of the signal was performed with enhanced luminescence assays Clarity and Clarity Max (1705061 and 1705062, Bio-Rad) or SuperSignal West Pico and Femto (34080 and 34095, ThermoFisher Scientific) using ChemiDoc (Bio-Rad). For loading control, membranes were stained with Ponceau S (09276-6X1EA-F, Sigma Aldrich).

### Immunoprecipitation (IP) assays

Five 10 mm leaf discs were collected from *N. benthamiana* leaves at 2-3 days post agroinfitration and ground in liquid nitrogen. All further steps were performed at 4°C if not mentioned otherwise. Soluble fraction was extracted in 5 ml of the buffer containing 50 mM Tris-HCl pH 7.5, 150 mM NaCl, 10% glycerol, 5 mM DTT, 1% Triton X-100 and EDTA-free 1x Plant Protease Inhibitor Cocktail (11873580001, Sigma Aldrich). Debris was removed by 2×15 min centrifugation at 14,000 g. All IPs were performed with 10 μl of anti-FLAG M2 Affiinity Gel slurry (A2220, Sigma Aldrich). After 2.5 h of incubation under constant rotation, beads were washed in 4-5 ml of the extraction buffer and eluted by boiling in 100 μl of 2xLaemmli for 10 min at 95°C.

### *Xanthomonas* infection assays in the presence of *A. tumefaciens*

*Xanthomonas campestris* pv. *vesicatoria* (*Xcv*) (85-10) (Thieme et al., 2005) (*Xanthomonas euvesicatoria*) kindly provided by Ulla Bonas was added to *A. tumefaciens* mixes to a final OD_600_=0.0005. *A. tumefaciens* strains were prepared as for the transient expression assays except without a P19 expressing strain. To ensure equal OD_600_ in all samples, *A. tumefaciens* expressing *p35S:YFP* was used in all experiments as filler. The bacterial mix was syringe-infiltrated into *N. benthamiana* leaves. *A. tumefaciens pAtEDS1:3xFLAG-SlEDS1* complemented *Xcv* susceptibility in *N. benthamiana eds1a* in an OD_600_ range of 0.05 to 0.6. For consistency between the cell death assays, a final *A. tumefaciens* OD_600_ =0.2 was used for each strain. After infiltration, plants were placed in a long-day chamber (16 h light / 8h dark at 25°C/23°C). Bacteria were isolated at 0 dpi (three 8 mm leaf discs served as three technical replicates) and 6 dpi (four 8 mm leaf discs representing four technical replicates), and dilutions were dropped onto NYGA supplemented with Rifampicin 100 mg/l and Streptomycin 150-200 mg/l. In statistical analysis of *Xcv* titers at 6 dpi, results from independent experiments (biological replicates) were combined.

Normality of residuals distribution and homogeneity of variance was assessed visually and by Shapiro-Wilcoxon as well as Levene tests (p>0.05). If both conditions were met, ANOVA was followed by Tukey’s HSD test (α=0.001), otherwise Nemenyi test with Bonferroni correction for multiple testing was applied (α=0.01).

### Cell death assays in *N. benthamiana*

After agroinfiltration, *N. benthamiana* plants were placed under a 16 h light/8 h dark regime at 22°C. Six 8 mm leaf discs from *N. benthamiana* agroinfiltrated leaves were taken at 3 dpi, washed in 10-20 ml of mQ for 30-60 min, transferred to a 24-well plate with 1 ml mQ in each well and incubated at room temperature. Ion leakage was measured at 0 and 6 h with a conductometer Horiba Twin Model B-173. For statistical analysis, results of measurements at 6 h for individual leaf discs (each leaf disc represents a technical replicate) were combined from independent experiments (biological replicates). Data were checked for normality of residuals distribution and homogeneity of variance using visual examination of the plots and Shapiro-Wilcoxon and Levene tests (p>0.05). If both conditions were met, ANOVA was followed by Tukey’s HSD posthoc test (α=0.001). Otherwise, non-parametric Nemenyi test with Bonferroni correction for multiple testing was applied (α=0.01). For visual assessment of cell death symptoms, infiltrated leaves were covered in aluminum foil for 2 d and opened to “dry” the lesions and enhance visual symptoms at 3 dpi.

### *Pseudomonas* infection and cell death assays in Arabidopsis

*Pseudomonas syringae* pv. *tomato* DC3000 (*Pst*) *pVSP61 avrRps4* (Hinsch and Staskawicz, 1996) was syringe-infiltrated into leaves at OD_600_ =0.0005 in 10 mM MgCl_2_. After infiltration, lids were kept on trays for 24 h and then removed. Bacteria were isolated at 0 dpi (6 to 8 5 mm leaf discs making 3-4 technical replicates) and 3 dpi (10 to 12 5 mm leaf discs distributed over 5-6 technical replicates). Dilutions were plated onto NYGA plates supplemented with rifampicin 100 mg/l and kanamycin 25 mg/l. For statistical analysis, bacterial titers from independent experiments (biological replicates) were combined. Normality of residuals distribution and homoscedasticity was checked visually and with formal Shapiro-Wilcoxon and Levene tests (α=0.05). Collected titer data were considered suitable for ANOVA and Tukey’s HSD test (α=0.001). For cell death assays, *Pseudomonas fluorescens Pf*0-1 *pEDV6 avrRps4* (Sohn et al., 2014) was grown at 28°C on King’s B medium (tetracycline 5 mg/l, chloramphenicol 30 mg/l), resuspended at a final OD_600_=0.2 in 10 mM MgCl_2_ and syringe-infiltrated into leaves. Ten leaves (technical replicates) per genotype were infiltrated for each independent experiment (biological replicate). Ion leakage assays were performed at 0 and 8 hpi as described (Heidrich et al., 2011), with an independent experiment considered as a biological replicate. Cell death symptoms visible as collapse of infiltrated areas of leaves were recorded at 24 hpi.

### *Hyaloperonospora arabidopsidis* (*Hpa*) Ewma1 infection assays

Seedlings from segregating (3:1) Arabidopsis T3 transgenic lines coexpressing 3xFLAG-EDS1 and YFP-PAD4 from Arabidopsis or tomato were preselected at 10 d on 1/2 MS plates supplemented with phosphinothricin (10 mg/l). A Col *35S:StrepII-3xHA-YFP* transgenic line and *eds1-2 pad4-1 sag101-1* used as controls were pre-grown on PPT plates alongside the test lines. Ws-2 seedlings were grown on 1/2 MS plates without PPT. After selection, seedlings were transplanted onto soil in Jiffy pots and grown for additional 7 d under a 10 h light /14 h dark, 22°C/20°C regime. *Hpa* Emwa1 spray inoculation (40 conidiospores/μl dH_2_O) was performed as described (Stuttmann et al., 2011). *Hpa* colonization was quantified by counting conidiospores on leaves at 7 dpi. In statistical analysis, counts normalized per mg of fresh weight (five counts used as technical replicates) from independent experiments (biological replicates) were combined. Significance of difference in spore counts was assessed with a non-parametric Nemenyi test and Bonferroni correction for multiple testing (α=0.01).

### Laser scanning fluorescence microscopy

Analysis of protein subcellular localization after transient expression in *N. benthamiana* was performed 2-3 dpi with the exception *At*NRG1.1 and *At*NRG1.2 experiments performed at 1 dpi to avoid quenching of GFP signal due to *At*NRG1.2-triggered cell death. Fluorescence signals in 8 mm leaf discs transiently expressed proteins was recorded on a laser-scanning confocal microscopes LSM780 or LSM700 (Zeiss) and generally under conditions when intensity of only a small fraction of pixels was saturated. Z-stacks were projected with ZEN (Zeiss) or Fiji using maximum intensity or standard deviation methods. Used objectives: 40x (NA=1.3 oil or 1.2 water) and 63x (NA=1.4 oil or 1.2 water).

### Generation of Arabidopsis *nrg1.1 nrg1.2* (*n2*) and *nrg1.1 nrg1.2 adr1 adr1-L1 adr1-L2* (*n2a3*) mutants

Arabidopsis *n2* and *n2a3* mutants were generated using targeted mutagenesis with the Clustered Regularly Interspaced Short Palindromic Repeats (CRISPR) – CRISPR-associated 9 (Cas9) method. Six guide RNAs (Supplemental Table 3) were designed to target the first two exons in *AtNRG1.1* and *AtNRG1.2* using CRISPR-P 2.0 (Liu et al., 2017). *AtNRG1.3* was not targeted because it is likely a pseudogene (Castel et al., 2018; Wu et al., 2018b). Two arrays of three fusions “*pU6:sgRNA_backbone-Pol_III_terminator*” each were synthesized (GeneArt, ThermoFisher Scientific) based on a template from (Peterson et al., 2016) with flanking SbfI/PmeI/SmaI sites for merging via restriction-ligation. The merged single array was further cloned into the pKIR1.0 (Tsutsui and Higashiyama, 2017) at the SbfI restriction site. To generate *n2* and *n2a3* mutant lines, the construct was electroporated into *A. tumefaciens* GV3101 pMP90RK for subsequent floral dip transformation (Logemann et al., 2006) into Col and Col *adr1 adr1-L1 adr1-L2* ((Bonardi et al., 2011)), respectively. T1 plants with active gRNA-Cas9 were preselected with the T7 endonuclease 1 assay (Hyun et al., 2015) or by direct sequencing of PCR products covering the target regions. Absence of the *Cas9* construct in lines homozygous for the *nrg1.1 nrg1.2* double mutation in *n2* and *n2a3* was tested with PCR using oligonucleotides matching the *Hyg* resistance gene in the T-DNA insertion and visually as lack of red fluorescence in the seed coat (Tsutsui and Higashiyama, 2017). One homozygous line free of the mutagenesis construct was selected for *n2* and *n2a3*. Mutations detected in the *AtNRG1.1* and *AtNRG1.2* genes in *n2* and *n2a3* lines are shown in Supplemental Figure 8A. Oligonucleotides used for genotyping of the mutants are listed in the Supplemental Table 3.

### Identification of orthogroups (OGs)

To build OGs from predicted 52 plant proteomes (listed in Supplemental Table 4), results of bidirectional BLASTP search (all versus all, E-value cut-off 1e-3, ncbi-blast-2.2.29+; (Altschul et al., 1990)) were used for orthology inference in orthomcl (v.2.0.9, E-value cut-off is 1e-5) with mcl clustering tool (v. 14-137) (Li et al., 2003).This resulted in 99,696 OGs. OrthoMCL generated OGs for EDS1, PAD4 and SAG101 were further verified with BLASTP (e-value 0.00001) against Arabidopsis (TAIR10). Original SAG101 OG appeared to be contaminated with AT3G01380, while EDS1 and PAD4 OGs contained only respective Arabidopsis hits. To systematically filter for high confidence EDS1-family orthologs, EDS1, PAD4 as well as BLASTP-verified SAG101 sequences were tested for the presence of EP domain (Hidden-Markov Model (HMM) profile for EP domain in Supplemental Dataset 4, hmmsearch--incE 0.00001 in HMMER 3.1b2 (Eddy, 2011), ≥50 amino acid long match). EP domain HMM was obtained with hmmbuild (in HMMER 3.1b2 (Eddy, 2011), default parameters) using MUSCLE multiple sequence alignment for EP domain sequences found by BLASTP (-evalue 0.01) with EP domains of *At*EDS1 (Q9XF23 385-623), *At*PAD4 (Q9S745 300-541), *At*SAG101 (Q4F883 291-537) (Wagner et al., 2013) against proteomes of 32 plant species from algae to *Arabidopsis thaliana* (Supplemental Table 5). Finally, too short (≤400 aa) and too long (≥1200) sequences in OrthoMCL-derived OGs were removed. A full pipeline and scripts to extract EP sequences and build HMM are in the Github repository “Lapin_Kovacova_Sun_et_al” (https://github.com/ParkerLabMPIPZ/Lapin_Kovacova_Sun_et_al). Filtered EDS1-family OGs are referred to as “high confidence orthologs” (Supplemental Datasets 5, 6 and 7). Their counts are given in Supplemental Table 1. BLASTP against TAIR10 did not detect contamination of ADR1 and NRG1 OrthoMCL OGs with other proteins. Their counts are also provided in Supplemental Table 1.

Additional manual searches for EDS1-family and NRG1 orthologs were performed using reciprocal BLASTP with Arabidopsis EDS1, PAD4, SAG101 and NRG1.1 sequences against spinach (*Spinacia oleracea*, bvseq.molgen.mpg.de; v.1.0.1), raspberry (*Rubus occidentalis*, rosaceae.org; v1.0.a1), jujube (*Ziziphus jujube* (Liu et al., 2014)), sesame (*Sesamum indicum*, Sinbase; v1.0) and quinoa (*Chenopodium quinoa* (Zou et al., 2017)). *Nicotiana benthamiana* EDS1-family sequences were obtained with tblasn searches of tomato sequences EDS1-family sequences on solgenomics.net and match sequences from (Ordon et al., 2017; Gantner et al., 2019). To search for EDS1, PAD4, ADR1 and NRG1 orthologs in *Silene* genus (Balounova et al., 2019), BLASTX (-word_size 4-evalue 1e-20) was performed with Arabidopsis amino acid sequences against nonfiltered *de novo* transcriptome assemblies. We considered a gene to be present in the assembly, if Arabidopsis sequence had a significant match with a unique contig.

### Phylogenetic and conservation analyses

Full-length EDS1-family sequences including high-confidence EDS1, PAD4 and SAG101 orthologs (Supplemental Datasets 5, 6, 7) and additional sequences from literature and other databases are provided in the Supplemental Dataset 1. To prepare EDS1-family maximum likelihood (ML) tree, the EDS1-family protein sequence alignment produced with mafft (version mafft-7.221, linsi, 100 iterations; (Katoh et al., 2002; Katoh and Standley, 2013)) was filtered using Gblocks (gap positions <50%, number of contiguous non-conserved positions - 15, minimum length of conserved block – 4; (Castresana, 2000; Talavera and Castresana, 2007)) leaving 101 positions in 12 blocks (Supplemental Dataset 2). The best evolutionary model (JTT+G) was selected with protTest3 (Darriba et al., 2011) based on the BIC criterion. The best ML tree was calculated with RAxML v.8.1.21 (-f a, 1000 bootstraps; (Stamatakis, 2014)). For Bayesian inference of EDS1-family protein phylogeny, we used MrBayes-3.2.6 with the same alignment as used for the ML tree (# generations – 5000000, # runs – 4, aa model – mixture of models, gamma rates; (Huelsenbeck and Ronquist, 2001; Ronquist and Huelsenbeck, 2003)). Annotated phylogenetic trees are available via iTOL (link in the section with accession numbers; (Letunic and Bork, 2016)) For the best nucleotide-binding domain found in Apaf-1, R-proteins and CED-4 (NBARC) domain ML tree, NBARC domain sequences were extracted based on the tabular output of hmmsearch (--incE 0.01, HMMER 3.1b2 (Eddy, 2011); PFAM PF00931.21 pfam.xfam.org) and aligned with mafft (version mafft-7.407, linsi, 1000 max iterations, (Katoh et al., 2002; Katoh and Standley, 2013)). The NBARC domain alignment without editing was supplied to RAxML (v.8.2.10,-f a, 800 bootstraps, LG model with empirical amino acid frequencies proposed via-m PROTGAMMAAUTO; this model was selected as best fitting in protTest3 (Darriba et al., 2011) as well).Gblocks-filtered NBARC domain alignment produced similar topology, but lower bootstrap support values on almost all branches. The annotated NBARC phylogenetic tree is available via iTOL (link in the section with accession numbers; (Letunic and Bork, 2016)).

For calculations of EDS1 family evolutionary conservation rates, amino acid sequences were aligned with mafft (version mafft-7.221-without-extensions, linsi, 100 iterations). Branch lengths of ML phylogenetic trees for EDS1, PAD4 and SAG101 built with RAxML (version standard-RaxML-8.1.21; (Stamatakis, 2014)) were optimized with rate4site package (version - rate4site-3.0.0, default parameters, background optimization with gamma model; (Pupko et al., 2002)). Mapping of the evolutionary rates onto the structure *At*EDS1-*At*SAG101 or homology-based model *At*EDS1-*At*PAD4 (Wagner et al., 2013) was performed in PyMol v2.0.7.

### Positive selection tests for EDS1

Analyses of evolutionary pressure acting on EDS1 sequence as the whole, as well as per site was performed with PAML package (Yang, 2007). The CODEML program of PAML 4.9a (Yang, 2007) was employed to estimate the ratio (ω) of the non-synonymous substitution rate (dN) to the synonymous substitution rate (dS). In all models, the reference tree was an unrooted maximum likelihood phylogenetic tree of EDS1 sequences with optimized branch lengths (CODEML program with the codon model M0 and the site model NS0, as recommended in PAML FAQ doc (page 14, http://abacus.gene.ucl.ac.uk/software/pamlFAQs.pdf)). The equilibrium frequencies of codons were calculated from the nucleotide frequencies (CodonFreq = 2) using jmodeltest-2.1.10 (Guindon et al., 2003; Darriba et al., 2012). Modelling of all models listed in Supplemetary Table 2 was done with the following initial ω values: ω = 0.1, ω = 0.5, ω = 1 and ω = 2. Since multinucleotide mutations can lead to false inference of positive selection (Venkat et al., 2018), we provide alignments at positions with inferred positive selection in *Brassicaceae* EDS1 (Supplemental Figure 1D).

### R packages frequently used in this study

The following R packages were utilized (R core team 2016, bioconductor.org): ggplot2 (http://ggplot2.org; 3.0.0), PMCMRplus (https://CRAN.R-project.org/package=PMCMRplus; 1.0.0), multcompView (https://CRAN.R-project.org/package=multcompView; 0.1-7), bioStrings (2.42.1).

## Accession numbers

Accession numbers of EDS1, PAD4 and SAG101 orthologs used in the study:

*At*EDS1 (AT3G48090.1), *At*PAD4 (AT3G52430.1), *At*SAG101 (AT5G14930.2), *Sl*EDS1 (Solyc06g071280.2.1), *Sl*PAD4 (Solyc02g032850.2.1), *St*EDS1 (PGSC0003DMP400055762), *St*PAD4 (PGSC0003DMP400034509), *Bd*EDS1 (XP_003578076.1), *Bd*PAD4 (XP_003577748.1), *Hv*EDS1 (MLOC_67615.1), *Hv*PAD4 (HORVU4Hr1G043530.1), *Nb*SAG101a (Niben101Scf00271g02011.1), *Nb*SAG101b (Niben101Scf01300g01009.1).

Accession numbers of NBARC-containing proteins used to infer phylogenetic placement of ADR1 and NRG1 NBARC domains: *At*NRG1.1 (Q9FKZ1), *At*NRG1.2 (Q9FKZ0), *Nb*NRG1 (Q4TVR0), *Sl*ADR1 (Solyc04g079420), *Os*ADR1 (LOC_Os12g39620.3), *Hv*ADR1 (A0A287QID5), *At*ADR1 (Q9FW44), *At*ADR1-L1 (Q9SZA7), *At*ADR1-L2 (Q9LZ25), *Sl*Bs4 (Q6T3R3), *Lus*L6 (Q40253), *Nb*Roq1 (A0A290U7), *At*RPP1 (F4J339), *At*RPP4 (F4JNA9), *At*RPS4 (Q9XGM3), *At*RPS2 (Q42484), *Os*RLS1 (Q6Z6E7), *St*Rx (Q9XGF5), AT5G56220 (Q9FH17), *Cc*Bs2 (Q9SNW0), *Sl*NRC1 (A1X877), *Sl*NRC2 (K4CZZ5), *Hv*MLA10 (Q6WWJ4), *At*RPP8 (Q8W4J9), *At*RPM1 (Q39214), *At*ZAR1 (Q38834).

Annotated EDS1 family phylogenetic trees:

ML tree https://itol.embl.de/tree/1953746254251611535639755

Bayesian tree https://itol.embl.de/tree/195374625452181536083186

Annotated ML tree for NBARC domains from selected NLR proteins

https://itol.embl.de/tree/1953746254304461545300543

<u>Content of GitHub repository “Lapin_Kovacova_Sun_et_al”:</u> pipeline and scripts to derive EP domain HMM (sub-directory “EP_domain_HMM”), pipeline and scripts used to filter OrthoMCL EDS1-family OG and obtain high-confidence sequences (sub-directory “high_confidence_OG”). Link - https://github.com/ParkerLabMPIPZ/Lapin_Kovacova_Sun_et_al

## Supplemental Data Files

Supplemental Table 1. Counts of EDS1, PAD4, SAG101, ADR1 and NRG1 orthologs in 52 green plants

Supplemental Table 2. Selection pressure acting on EDS1 sequences in *Poaceae, Solanaceae* and *Brassicaceae*

Supplemental Table 3. Sequences of oligonucleotides used in this study

Supplemental Table 4. Names of 52 green plant species used in the OrthoMCL analysis

Supplemental Table 5. Names of 32 green plant species used to build EP domain Hidden-Markov Model (HMM) profile

Supplemental Dataset 1. Sequences of EDS1-family proteins used for ML and Bayesian phylogeny inference (fasta format)

Supplemental Dataset 2. Gblocks-filtered alignment of EDS1-family sequences used for the phylogenetic analysis with RAxML and MrBayes

Supplemental Dataset 3. Correspondence between EDS1-family sequence names on the phylogenetic trees and in public databases

Supplemental Dataset 4. EP domain Hidden-Markov Model (HMM) profile

Supplemental Dataset 5. Sequences of high-confidence EDS1 orthologs (fasta format)

Supplemental Dataset 6. Sequences of high-confidence PAD4 orthologs (fasta format)

Supplemental Dataset 7. Sequences of high-confidence SAG101 orthologs (fasta format)

Supplemental Dataset 8. Sequence of the custom destination Gateway vector pENpAtPAD4 StrepII-YFP (.gbk format)

## Acknowledgements

We thank Ulla Bonas (Martin-Luther University, Halle) for the strain *Xcv* 85-10 and Artem Pankin (MPIPZ) for helpful discussions. This work was supported by the Max Planck Society and Deutsche Forschungsgemeinschaft (DFG) Grants CRC680 (JEP, DL, VK, AB), CRC670 (JEP, DB), CRC648 (JS), DFG-ANR Trilateral ‘RADAR’ grant (JD), an International Max-Planck Research School (IMPRS) doctoral fellowship (PvB) and a Chinese Scholarship Council PhD fellowship (XS). No conflict of interest declared.

## Author contributions

JEP and DL conceived the project. DL designed, coordinated and performed experiments, generated *n2 a3* mutant, and prepared GitHub repository with input from VK. VK designed and performed orthology inference and phylogenetic analyses. XS, NG, DB, JB and JD performed experiments. JD designed CRISPR-Cas9 mutagenesis constructs and generated the *n2* mutant. DB designed and generated structure-guided mutants. JB selected *Nb-epss* mutant and did EDS1/PAD4 ortholog cloning. JS provided *Nb-pss* mutants, an F2 segregating population to select *Nb-epss* and modules for golden gate cloning, and pENTR/D-TOPO clones of tomato EDS1 and PAD4. AB contributed to orthology analysis and discussions. DL and JEP wrote the manuscript with input from all co-authors.

