## Supplemental Figures and Tables for "A coevolved EDS1-SAG101-NRG1 module mediates cell death signaling by TIR-domain immune receptors"

SUPPLEMENTARY FIGURE 1

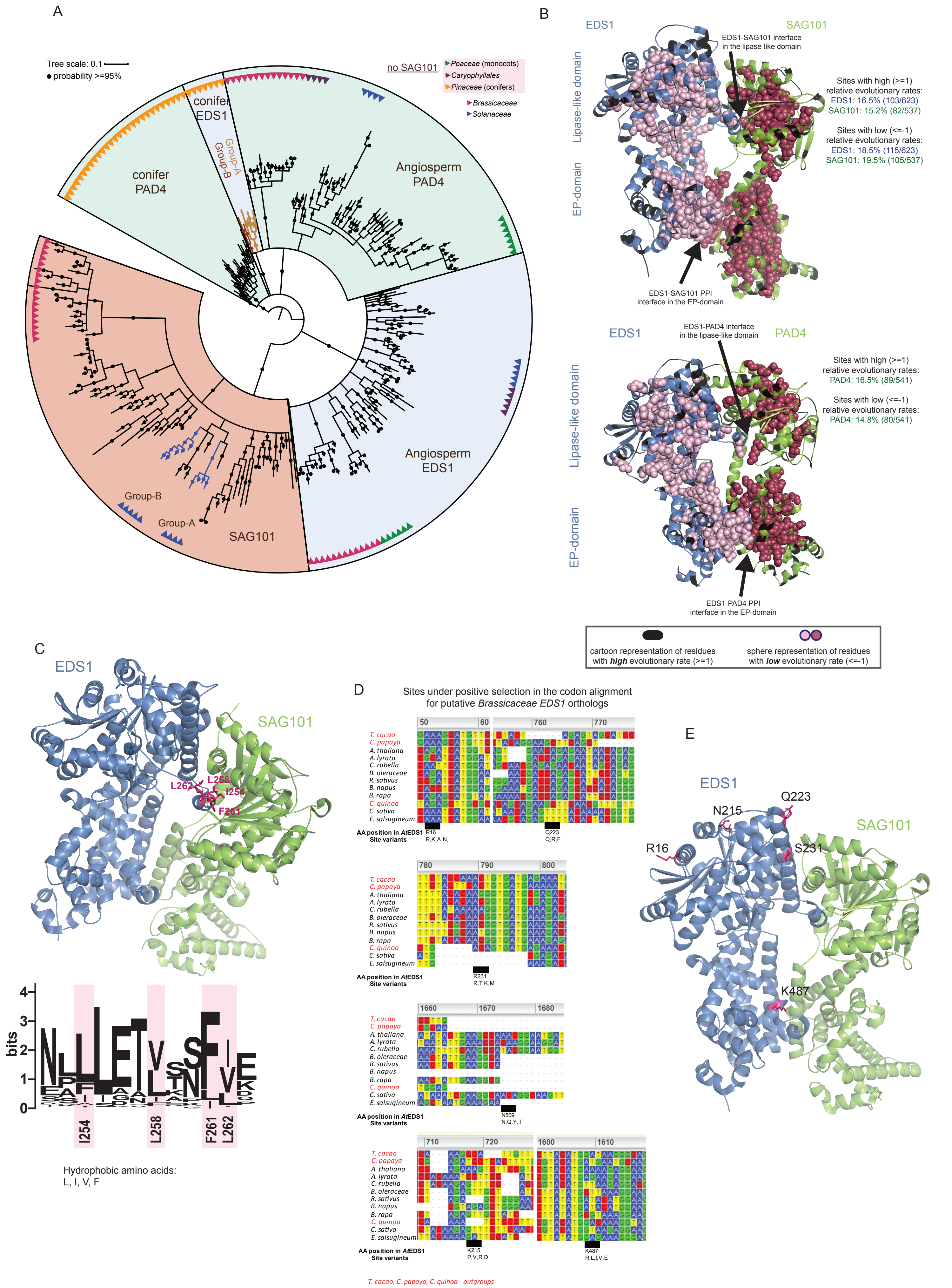

**Phylogenetic and conservation analysis of the EDS1-family protein sequences.**
(A) Phylogenetic tree of the 256 EDS1, PAD4 and SAG101 protein sequences obtained with Bayesian inference of phylogeny was based on the same alignment as the maximum likelihood (ML) tree on Figure 1A. Branches supported with posterior probability ≥95% are indicated with dots. EDS1, PAD4 and SAG101 orthologs form separate branches in flowering plants. Similarly, conifer EDS1 and PAD4 form distinct branches as well. Within conifer EDS1, two well-supported groups of sequences were detected (groups A and B). Two groups of SAG101 orthologs were found within Solanaceae (groups A and B) as well.

(B) Mapping of relative evolutionary rates for EDS1, PAD4 and SAG101 protein sequences on the *At*EDS1-*At*SAG101 structure (PDB 4NFU) or *At*EDS1-*At*PAD4 homology-based model. Black ribbons correspond to fast evolving positions (rates ≥1), while relatively slow evolving amino acids (rates ≤-1) are shown as spheres. 14.8-19.5% of amino acid positions in the EDS1-family proteins are evolving slower than expected based on the ML tree for a full-length protein alignment. These amino acids are hidden inside of the structure and often found on the EP domain interface. Fast evolving amino acids (15.2-16.5%) are predominantly surface exposed.

(C) Representation of *At*EDS1 'LLIF' motif (pink) on the *At*EDS1-*At*SAG101 structure and logo of the 'LLIF' motif alignment based on 78 EDS1 orthologs. Height of the amino acid corresponds to reflects its relative frequency at this position. The 'LLIF' motif is not conserved at the amino acid sequence level, but hydrophobic nature of the site is maintained.

(D) Alignment of *Brassicaceae EDS1* coding sequences at positions with signature of positive selection. Coordinates are relative to *Arabidopsis thaliana* EDS1 (AT3G48090). The alignment provides visual assessment of potential confounding effects caused by gaps and multinucleotide mutations (Venkat et al., 2018). The site N509 is likely a false positive call due to gaps in the alignment. Other detected sites with the exception of R231 contain multinucleotide mutations.

(E) Representation of five amino acid positions under positive selection in *Brassicaceae EDS1* on the structure of *At*EDS1-*At*SAG101. All amino acids are surface-exposed. Four out of five sites (16, 215, 223, 231) are located in the lipase-like domain, while K487 is in the EP domain facing away from the heterodimer cavity. Reference sequence Col AT3G48090 differs as several non-conserved positions from the crystallized *At*EDS1 sequence (PDB 4NFU) including K215N and R231S.

SUPPLEMENTARY FIGURE 2

A

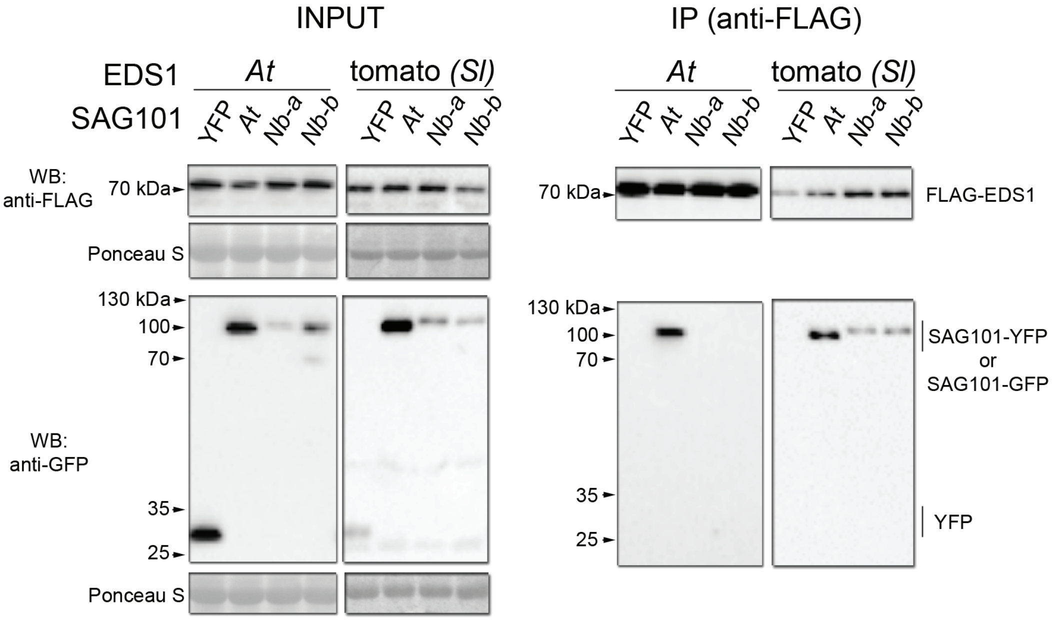


B

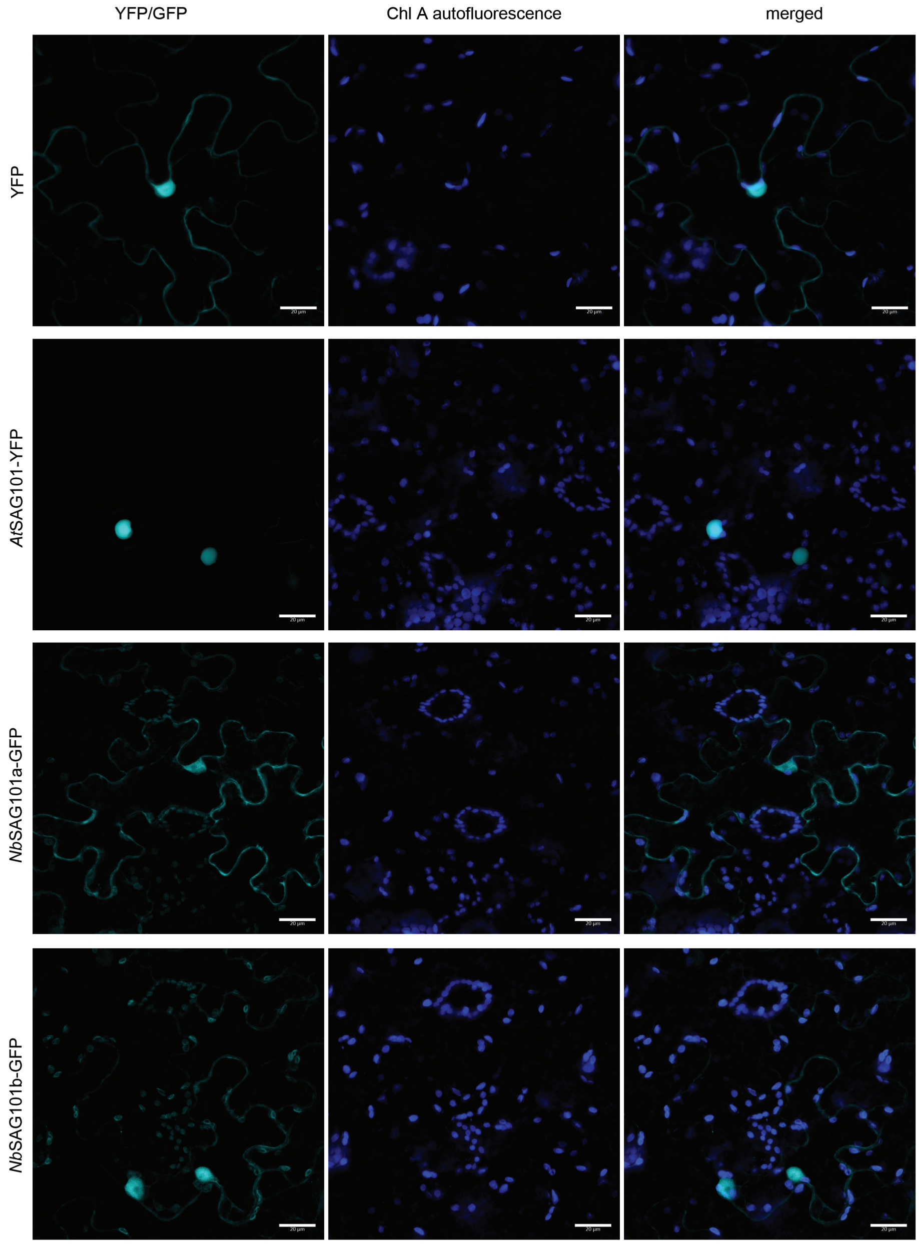


**Arabidopsis EDS1 does not interact in *N. benthamiana* with *Nb*SAG101a and *Nb*SAG101b.**

(A) A coimmunoprecitation (coIP) assay with anti-FLAG antibodies to test *in planta* interactions between Arabidopsis (*At*) *At*EDS1-FLAG (genomic sequence under *pAtEDS1* control) as well as tomato (*Sl*) FLAG-*Sl*EDS1 (*pAtEDS1* promoter control) and coexpressed *At*SAG101-YFP or *Nb*SAG101a-GFP and NbSAG101b-GFP (all *35S* promoter). Proteins were transiently expressed for 2-3 d via agroinfiltration into *Nb-epss* leaves. FLAG-*Sl*EDS1 forms stable complexes with all tested SAG101 GFP/YFP-fusions from *N. benthamiana* and Arabidopsis, while *At*EDS1-FLAG interacted only with *At*SAG101-YFP.

The experiment was performed twice with similar results.

(B) Subcellular localization of SAG101 proteins used the coIP assay in the panel A in *Nb-epss*. *At*SAG101-YFP is found primarily in the nucleus. *Nb*SAG101a-GFP and *Nb*SAG101b-GFP show nucleocytoplasmic localization. Scale bar is 20 µm. Confocal fluorescence imaging was performed in two independent experiments. At least 50 cells were observed per sample.

SUPPLEMENTARY FIGURE 3

A

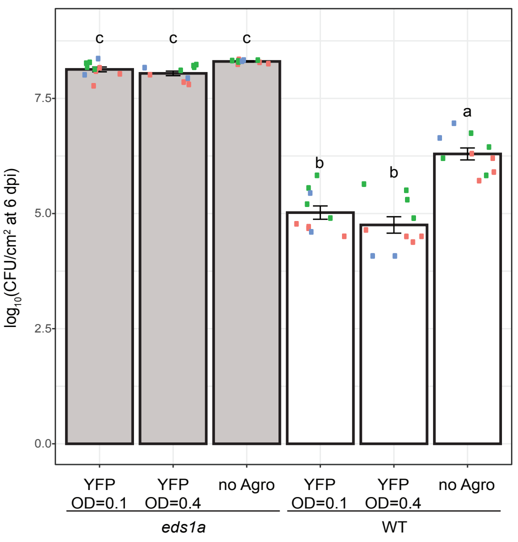


B

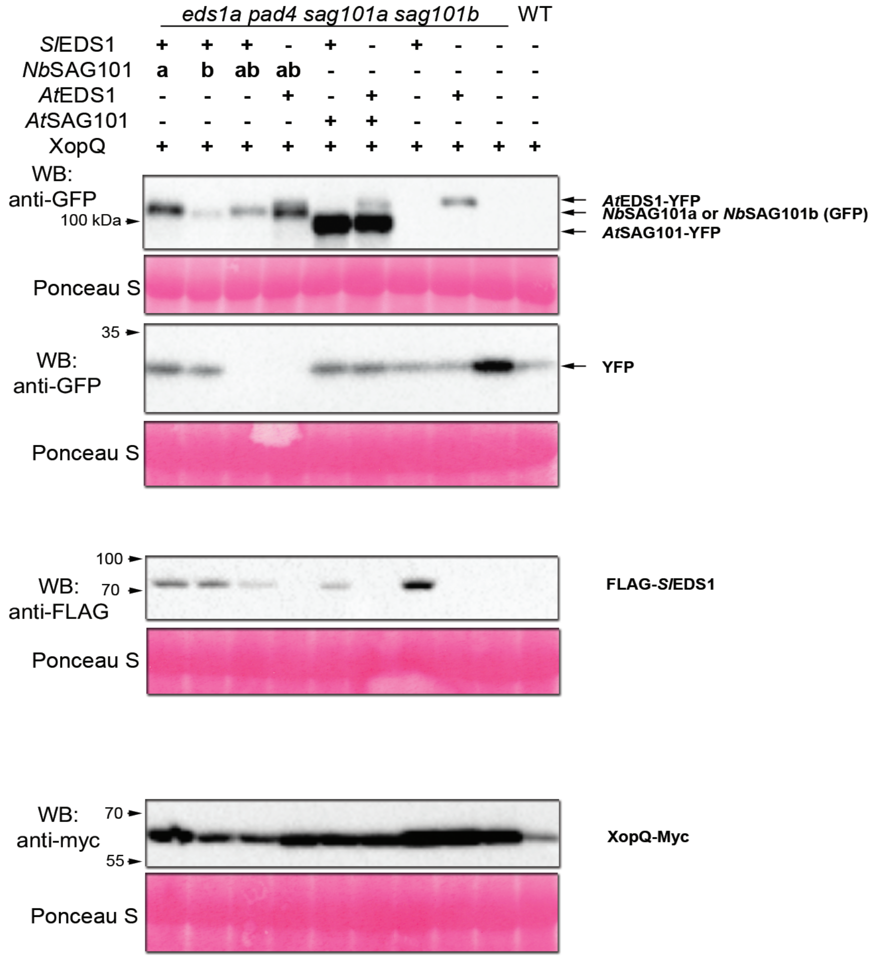


**Effect of Agrobacteria and expression of proteins under test in assays to complement *Roq1* resistance to *Xcv* and XopQ-triggered cell death signaling in *Nb-epss*.**
(A) Effect of *A. tumefaciens* at OD600=0.1 or 0.4 on *in planta Xcv* growth (OD600=0.0005) in *N. benthamiana* WT and *eds1a* plants. Leaves were syringe infiltrated with mixes of the two bacteria or *Xcv* alone. *Xcv* growth assessed as bacterial titer at 6 dpi is not affected by *A. tumefaciens* in the *Nbeds1a* mutant, however reduced *Xcv* growth was observed in the resistant WT compared to the *Xcv*-alone situation. The experiment was performed three times independently (colors of dots represent individual independent experiments used as biological replicates) with four technical replicates (bacterial extractions in one independent experiment) each. Results were analyzed with Nemenyi test using Bonferroni correction for multiple testing, grouping of genotypes is performed at α=0.01 (n=12).

(B) Western blot analysis of protein accumulation in the ion leakage assay to measure complementation of *Roq1* cell death in *Nb-epss* after agroinfiltration of *pAtEDS1:FLAG-SlEDS1, 35S:NbSAG101a-GFP, 35S:NbSAG101b-GFP, pAtEDS1:AtEDS1-YFP* (genomic sequence), *35S:AtSAG101-YFP* or *35S:YFP* (served as a filler to adjust for differences in OD600, labelled "-" in the sample description). Protein samples were collected at 2 dpi when visual cell death symptoms (drying out of lower side of the leaf) were visible. Ponceau S staining of the membrane shows similar loading of samples on the gel. The experiment was performed twice with similar results.

SUPPLEMENTARY FIGURE 4

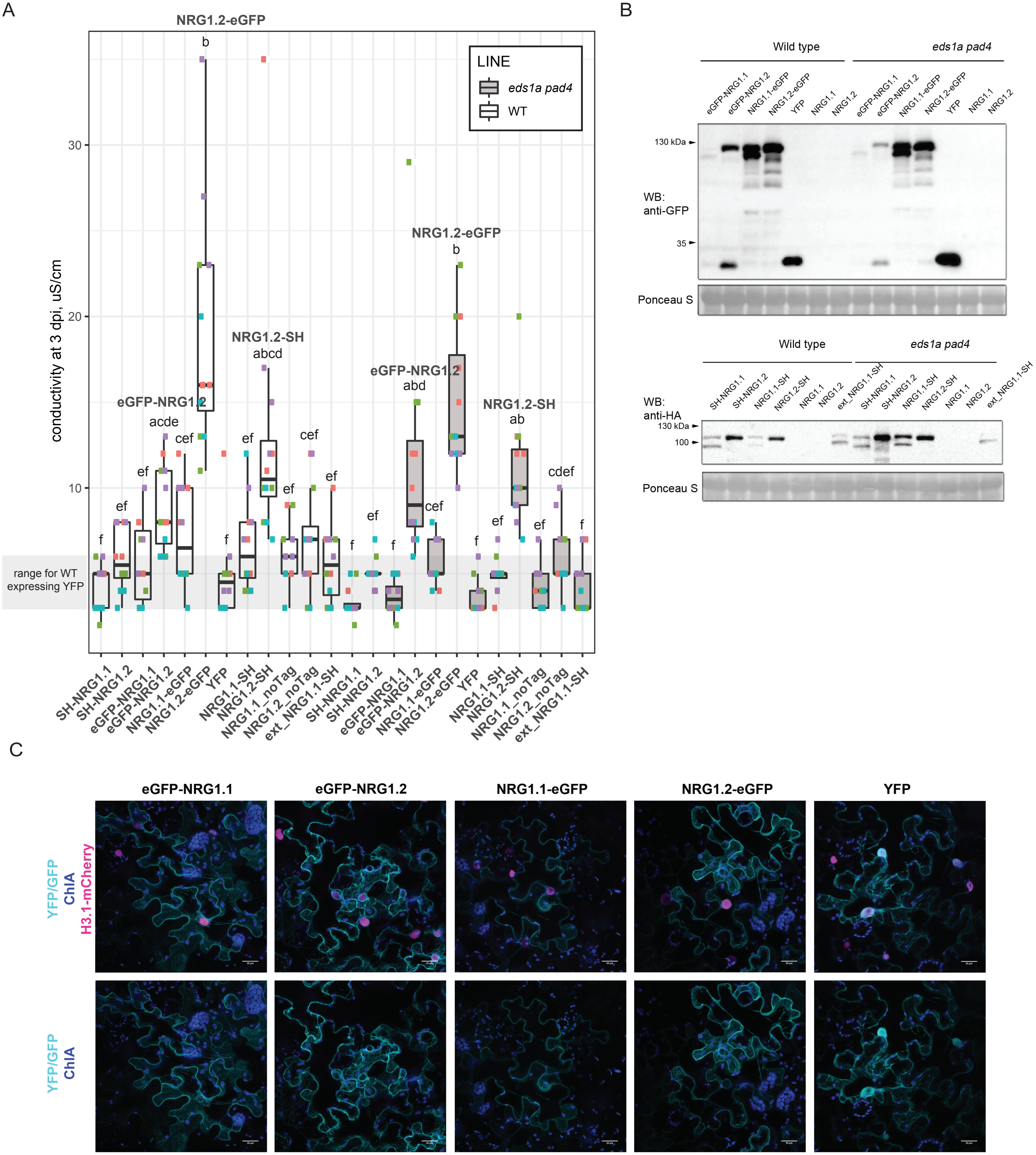


**Initial characterization of N- and C-terminally eGFP or SH tagged *At*NRG1.1 or *At*NRG1.2 after transient expression in *N. benthamiana* under the control of *35S* promoter.**
(A) Transient expression of eGFP-*At*NRG1.2, *At*NRG1.2-eGFP and *At*NRG1.2-SH causes cell death as measured in the ion leakage assay (3dpi) in both WT and *eds1a pad4 N. benthamiana* plants. Infiltration of *A. tumefaciens* to express non-tagged *At*NRG1.1 and *At*NRG1.2 under the control of *35S* promoter did not trigger an increase in conductivity above the negative control YFP. An alternative *At*NRG1.1 gene model (At5g66900.2; ext_NRG1.1-SH on the figure) fused to a C-terminal SH tag did not show symptoms of cell death either. The experiment was performed four times independently (biological replicates) with six leaf discs (technical replicates) in each experiment. Statistical analysis included Tukey's HSD test and subsequent grouping of samples according to their means at α=0.001.

(B) Western blot analysis of expression of tagged and untagged *At*NRG1.1 and *At*NRG1.2 in *N. benthamiana* WT and *eds1a pad4* plants at 2 dpi. eGFP-*At*NRG1.1, SH-*At*NRG1.1, *At*NRG1.1-SH are detected as a double band, whereas *At*NRG1.2 – as a single band independent of the tag or its position. The analysis was repeated three times with similar results. (C) Subcellular localization of eGFP N- and C-terminally tagged *At*NRG1.1 and *At*NRG1.2 in WT N. benthamiana leaves. Histone H3.1-mCherry (At5g10400) was used as marker of nuclear localization. Imaging was performed at 24 hours after *A. tumefaciens* infiltration since ongoing cell death quenched GFP signal in *At*NRG1.2 samples. Microscopy analysis was performed in three independent experiments. At least 50 cells were observed for each sample. Nuclear localization was observed for the tested *At*NRG1.1 and *At*NRG1.2 proteins only occasionally, in the vast majority of cells no GFP signal inside nuclei was detected. Scale bar = 20 µm.

SUPPLEMENTARY FIGURE 5

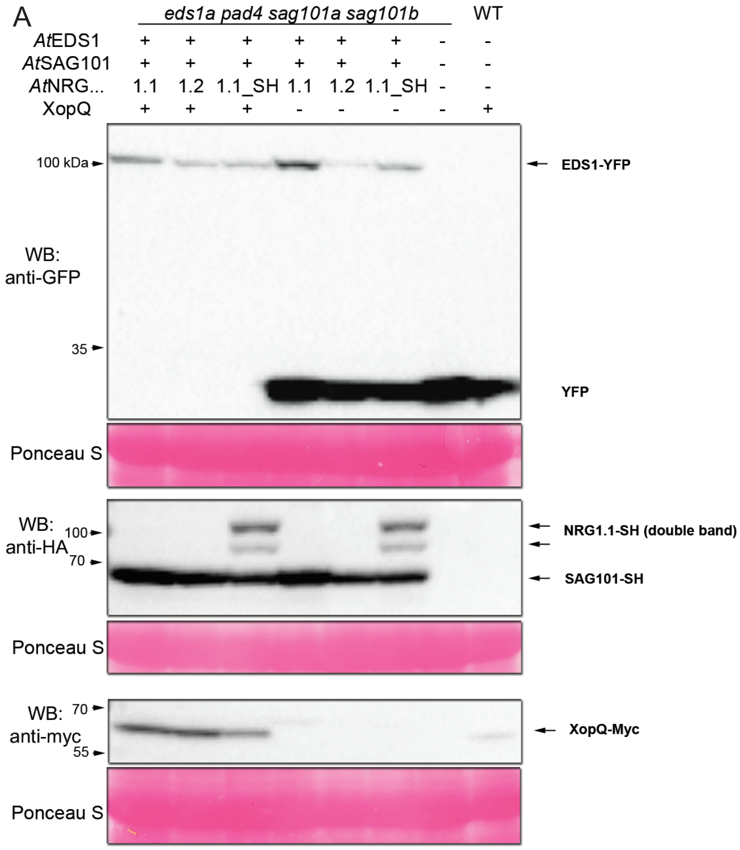


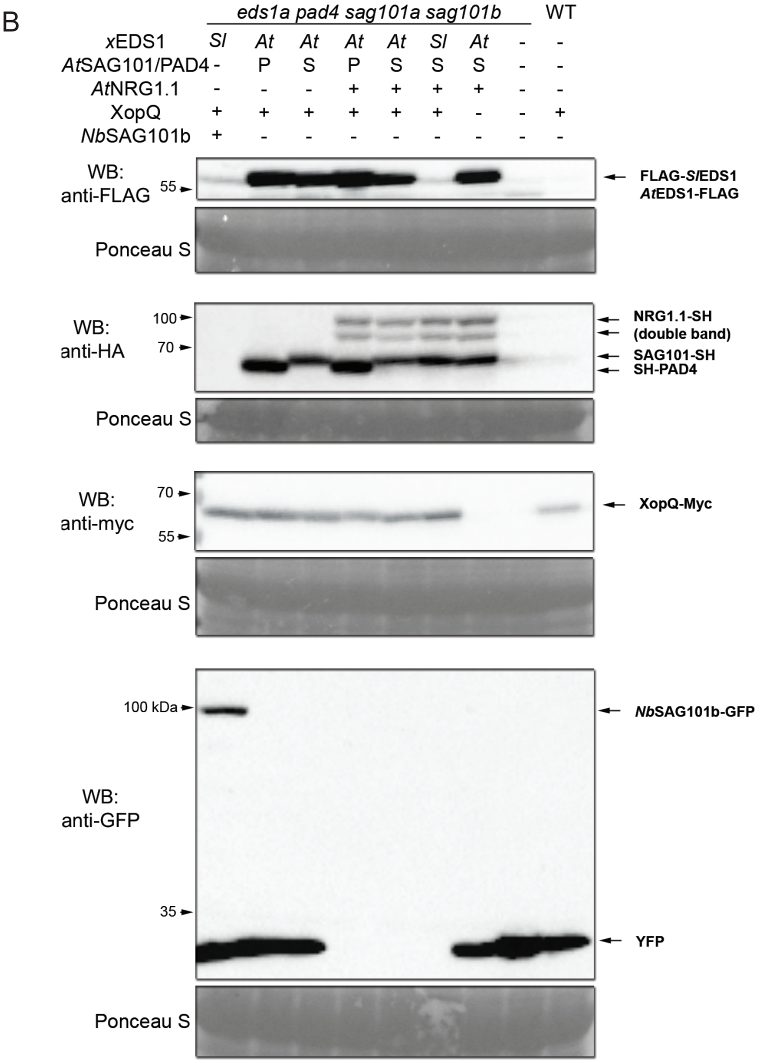


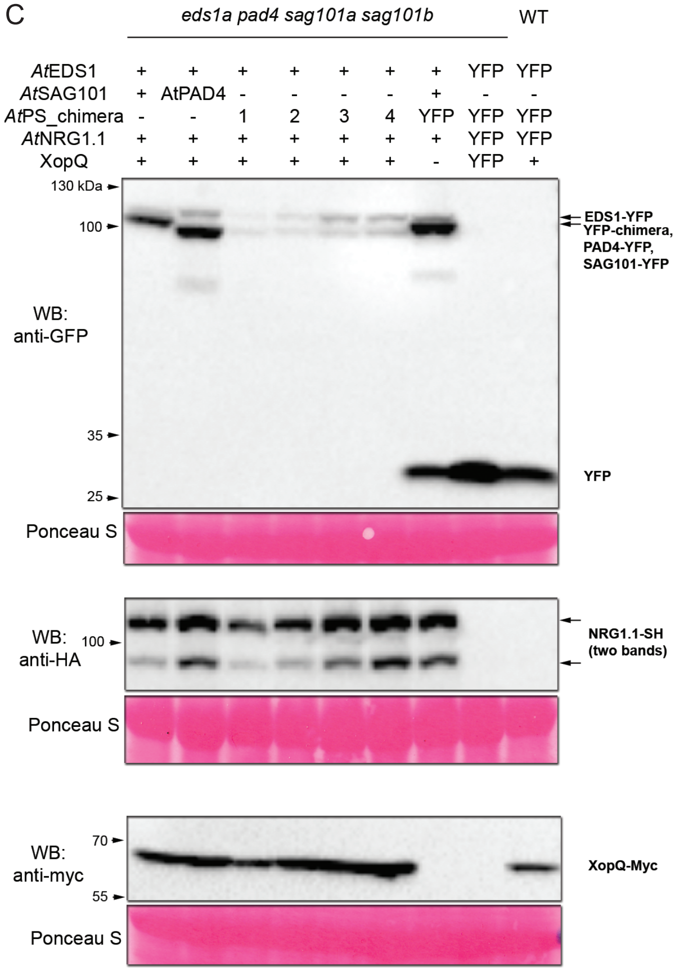


D

```
AtSAG101 276-316     IKGIEKRAEMKNLRFDMMFKKLNDMKISMAYIEWYKKKCKE
AtPAD4 286-326       IETATRIVRAPILRSAELANELASVLPARLEIQWYKDRCDA
align. score         *:   : ..   **   : ::* .:  :   *:***.:*.

chimera 1            IKGIEKRAEMKNLRFDMMFKKLNDMKISMAYIEWYKKKCKE
chimera 2            IETATRIVRAPILRFDMMFKKLNDMKISMAYIEWYKKKCKE
chimera 3            IETATRIVRAPILRSAELANELASVLPARLEIEWYKKKCKE
chimera 4            IETATRIVRAPILRSAELANELASVLPARLEIQWYKDRCDA
```

**Protein accumulation in assays to test *At*SAG101 EP domain functions in the *Roq1* cell death reconstitution assays in *Nb-epss*.**

(A) Expression of proteins in the initial cell death reconstitution assay with Arabidopsis EDS1-SAG101-NRG1 protein module. *At*EDS1-YFP (genomic sequence), *At*SAG101-SH, *At*NRG1.1-SH, *At*NRG1.1 without a tag and nontagged *At*NRG1.2 were coexpressed with XopQ-myc or YFP ("-" in their description, to control for bacterial density at OD600) in *Nb-epss*. As a positive control for the ion leakage assay, XopQ-myc was expressed in WT. All proteins expressed as expected. The analysis was performed for two independent experiments with similar results.

(B) Expression of FLAG-*Sl*EDS1, *Nb*SAG101b-GFP, *At*EDS1-FLAG (genomic sequence), SH-*At*PAD4, *At*SAG101-SH, *At*NRG1.1-SH and XopQ-myc in *Nb-epss* in the reconstitution assay. WT plants expressed XopQ-myc as a control for the ion leakage assay. Samples with "-" in the description contain YFP. All proteins are detected as expected according to the agroinfiltration scheme. Western blot analysis was performed for two independent experiments with similar results.

(C) Expression of *At*EDS1-YFP (genomic sequence), *At*SAG101-YFP, *At*PAD4-YFP, N-terminally StrepII-YFP-tagged *At*PAD4/*At*SAG101 chimeras 1 to 4, *At*NRG1.1-SH, XopQ-myc and YFP in the indicated genotypes according to a scheme above the blot. All proteins are detected in accordance with the infiltration scheme, chimeras are expressed on similar levels with chimeras #1 and #2 showing tendency towards lower expression. Western blot analysis was conducted for samples from two independent experiments with similar results.

(D) Pairwise sequence alignment of *At*SAG101 (276-316 aa, in pink) and *At*PAD4 (286-326 aa, in green) in the area surrounding *At*SAG101 sequence critical to its activity in the cell death reconstitution assay (in bold).

SUPPLEMENTARY FIGURE 6

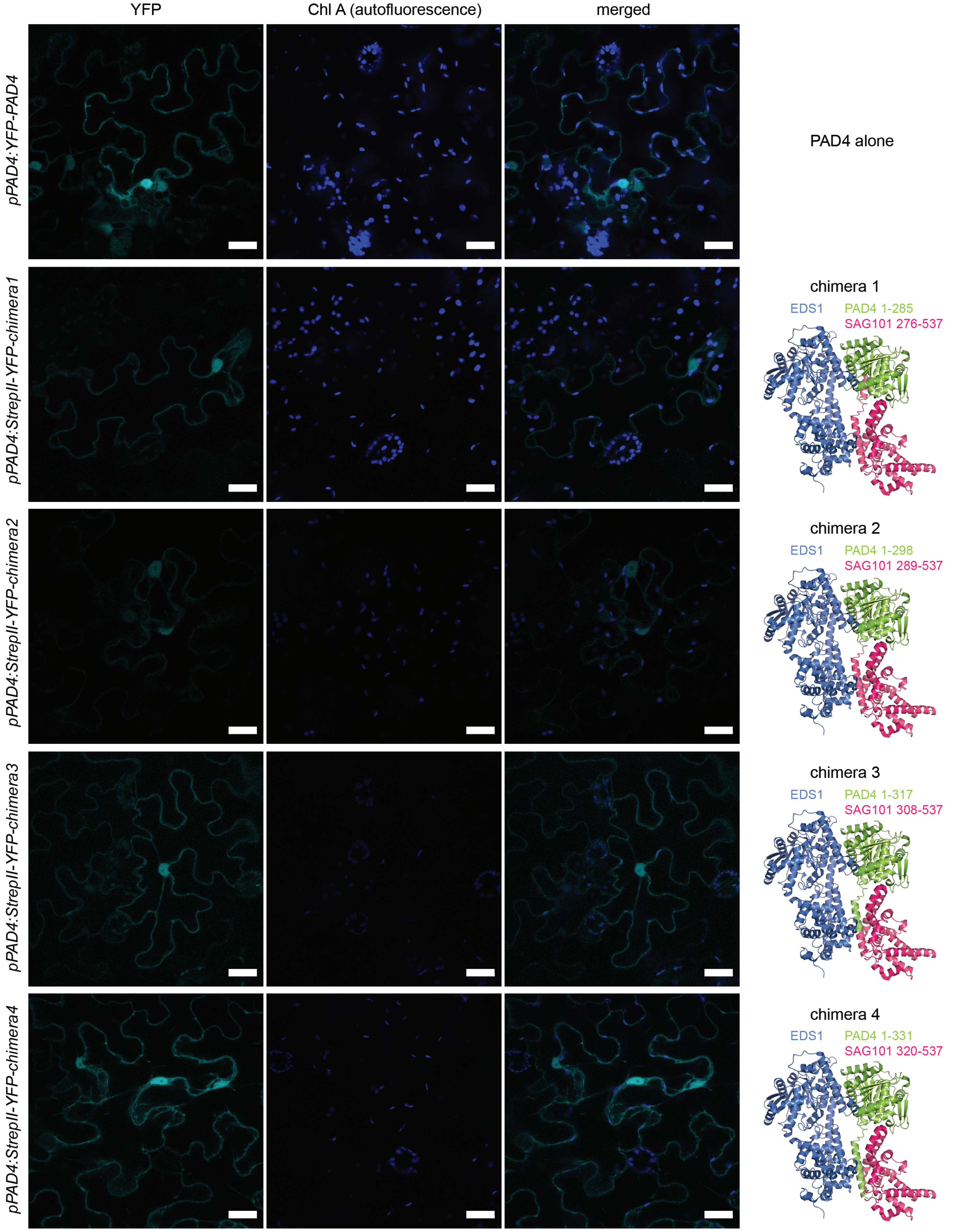


**Subcellular localization of *At*PAD4/*At*SAG101 chimeric proteins in *Nb-epss*.**
The chimeras were expressed under the control of *pAtPAD4* and were N-terminally tagged with StrepII-YFP. All tested chimeric proteins show nucleocytoplasmic localization similar to GFP-*At*PAD4 (under *pAtPAD4*). Imaging was performed at 2 d after *A. tumefaciens* infiltration in two independent experiments. At least 20 cells per sample were observed. Scale bar = 20 µm.

SUPPLEMENTARY FIGURE 7

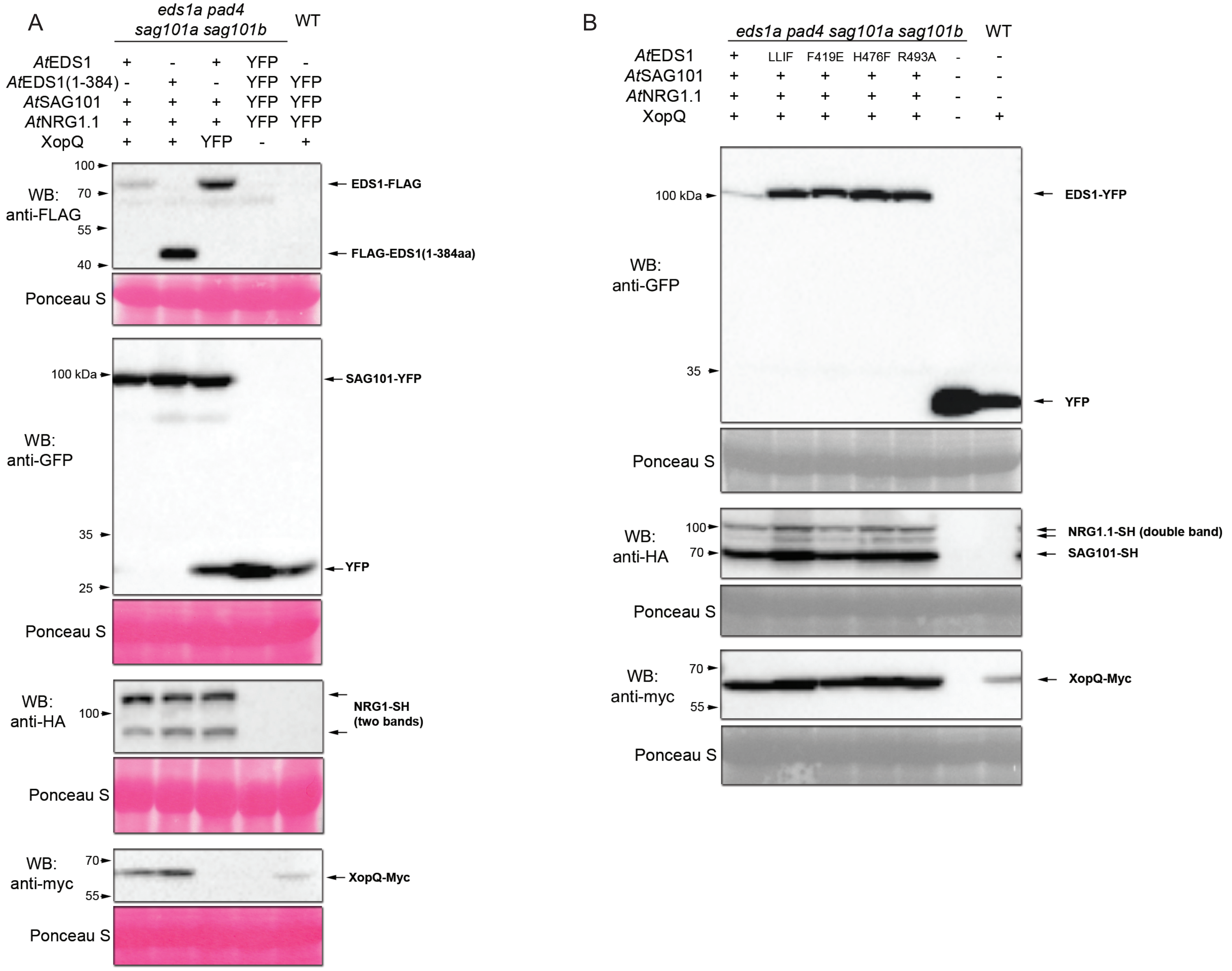


**Protein accumulation in assays to find sequence and structural determinants of *At*EDS1 activity in *Nb-epss Roq1* cell death reconstitution.**
(A) Expression of proteins in assays to test requirement for *At*EDS1 EP domain. Proteins as indicated above the blot were transiently expressed through agroinfiltration of *pAtEDS1:gAtEDS1-FLAG* (“g” for genomic sequence), *pAtEDS1: FLAG-cAtEDS1_1-384aa* (“c” for coding sequence), *35S:AtSAG101-YFP*, *35S:AtNRG1.1-SH*, *35S:XopQ-myc* or *35S:YFP*. All proteins are expressed as expected. Lower ion leakage in the sample #2 expressing a lipase-like domain of *At*EDS1 is not due to lower protein accumulation. The analysis was performed for two independent experiments with similar results.

(B) Protein accumulation in reconstitution assays with structure-guided *At*EDS1 EP domain mutant variants. *A. tumefaciens* carrying *pAtEDS1:gAtEDS1-YFP* or its variants (‘LLIF’, F419E, H476F, R493A), *35S:AtSAG101-SH*, *35S:AtNRG1.1-SH*, *35S:XopQ-myc* or *35S:YFP* (“-” in the sample description) were coinfiltrated according to the sample description above blots. Difference in the expression of *At*EDS1 protein variants does not explain differences observed in the ion leakage assay. The analysis was performed for two independent experiments with similar results.

SUPPLEMENTARY FIGURE 8

A

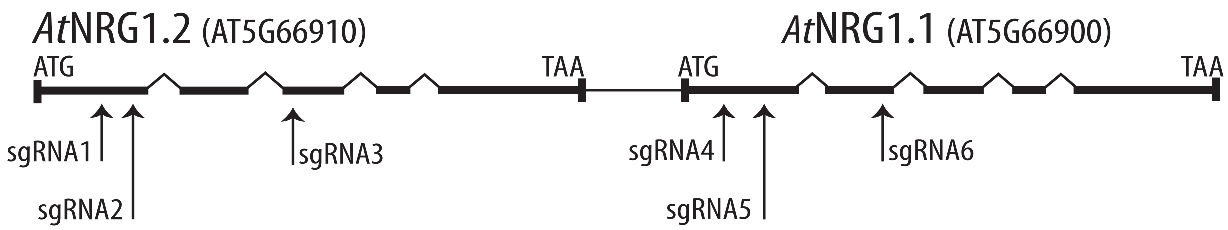


**Target site sgRNA1**

WT: CCG-**CTTGTTTCGAAGCGTTG**
*nrg1.1/1.2 :* CCGACTTGTTTCGAAGCGTTG

WT: CCG**CTTGTTTCGAAGCGTTG**
*nrg1.1/1.2 adr triple:* CCGCTTGTTTCGAAGCGTTG

**Target site sgRNA2**

WT: **GAAACTCCTTGACAATTCCG**TGG
*nrg1.1/1.2 :* GAAACTCCTTGACAATTCCGTGG

WT: **GAAACTCCTTGACAAT-------------------TCCG**TGG
*nrg1.1/1.2 adr triple:* GAAACTCCTTGACAAT-(32 bp insertion)-TCCGTGG

**Target site sgRNA3**

WT: **CCTCTCGTAATTGAAGTAGT**CGG
*nrg1.1/1.2 :* CCTCTCGTAATTGAAGT---CGG

WT: **CCTCTCGTAATTGAAGTA-GT**CGG
*nrg1.1/1.2 adr triple:* CCTCTCGTAATTGAAGTAAGTCGG

**Target site sgRNA4**

WT: CCC**TTGTCCAAGGATCTCGTATC**
*nrg1.1/1.2 :* CCCTT-TCCAAGGATCTCGTATC

WT: CCC**T----------------TGTCCAAGGATCTCGTATC**
*nrg1.1/1.2 adr triple:* CCCTAAAAAAAAGTTCAAAGCTTCCAAGGATCTCGTATC

**Target site sgRNA5**

WT: **GGAAGTTTCCAAGAGTCAAG**-----------------TAA
*nrg1.1/1.2 :* GGAAGTTTCCAAGAGTCAA-(45 bp deletion)-TAA

WT: **GGAAGTTTCCAAGAGTCAAG**TGGTA-----CG
*nrg1.1/1.2 adr triple:* GGAAGTTTCCAAGAGTCATATTTCTATGCATG

**Target site sgRNA6**

WT: CCT**TGTAA-CACGTCTCCTGATGA**
*nrg1.1/1.2 :* CCTTGTAAACACGTCTCCTGATGA

WT: CCT**TGTAACACGTCTCCTGATGA**
*nrg1.1/1.2 adr triple:* CCT**TGT**TTAATTCATCT**-----A**

B

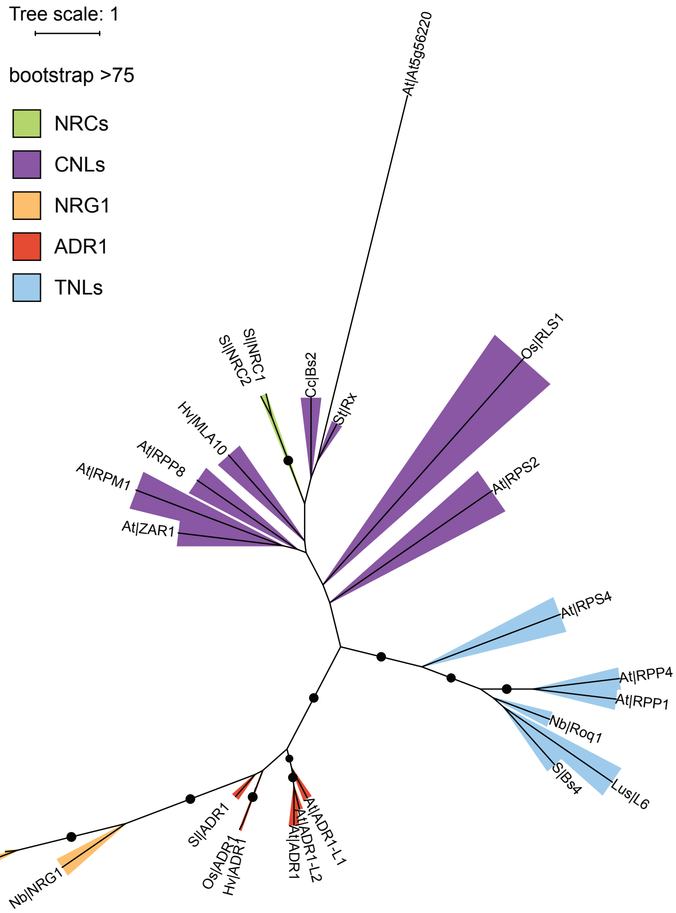


**Mutants for Arabidopsis *NRG1*-family genes generated in this study.**
(A) Schematic of Cas9 knock out mutants of *AtNRG1.1* and *AtNRG1.2* in Col and Col *adr1 adr1-L1 adr1-L2* (*adr triple*) mutant. Guide RNAs were designed to target first two exons of the *AtNRG1.1* and *AtNRG1.2* genes.

(B) Maximum likelihood phylogenetic tree of NBARC domains from selected NLR proteins from flowering plants including helper NLRs (NRC, ADR1, NRG1 orthologs) and Arabidopsis TIR2-NBARC domain protein AT5G56220. ADR1 and NRG1 NBARC domain sequences form a well-supported (bootstrap 96%) group distinct from other NBARC domains, therefore mutations in NRG1.1 and NRG1.2 were introduced in the adr triple mutant as well. Sequence IDs are provided in the section "Accession numbers".

Supplemental Table 1

Count of high-confidence EDS1-, ADR1- and NRG1-family orthologs in 52 green plants

| Species | EDS1 | PAD4 | SAG101 | ADR1 | NRG1 |
|---|---|---|---|---|---|
| *Arabidopsis thaliana* | 2 | 1 | 1 | 4 | 3 |
| *Actinidia chinensis* | 2 | 2 | 1 | 2 | 2 |
| *Amborella trichopoda* | 0 | 0 | 1 | 1 | 0 |
| *Aquilegia coerulea* | 1 | 1 | 0 | 1 | 0 |
| *Arabidopsis halleri* | 2 | 0 | 1 | 3 | 2 |
| *Arabidopsis lyrata* | 1 | 1 | 1 | 3 | 3 |
| *Arachis ipaensis* | 2 | 2 | 1 | 1 | 5 |
| *Beta vulgaris* | 1 | 1 | 0 | 1 | 0 |
| *Boechera stricta* | 1 | 1 | 1 | 3 | 2 |
| *Brachypodium distachyon* | 1 | 1 | 0 | 1 | 0 |
| *Brassica oleracea* | 2 | 4 | 5 | 3 | 1 |
| *Brassica rapa* | 3 | 2 | 3 | 4 | 6 |
| *Capsella grandiflora* | 1 | 1 | 1 | 4 | 2 |
| *Capsella rubella* | 1 | 2 | 1 | 5 | 2 |
| *Capsicum annum* | 1 | 0 | 1 | 1 | 2 |
| *Carica papaya* | 1 | 1 | 1 | 1 | 1 |
| *Castanea mollissima* | 5 | 1 | 8 | 0 | 0 |
| *Citrus clementina* | 4 | 1 | 1 | 1 | 2 |
| *Citrus sinensis* | 1 | 1 | 1 | 1 | 2 |
| *Coccomyxa subellipsoidea* | 0 | 0 | 0 | 0 | 0 |
| *Coffea canephora* | 1 | 1 | 1 | 1 | 1 |
| *Cucumis sativus* | 1 | 1 | 1 | 1 | 2 |
| *Eutrema salsugineum* | 1 | 2 | 3 | 2 | 4 |
| *Fragaria vesca* | 1 | 1 | 1 | 1 | 22 |
| *Gossypium raimondii* | 1 | 1 | 1 | 3 | 1 |
| *Linum usitatissimum* | 2 | 2 | 5 | 2 | 3 |
| *Malus domestica* | 5 | 1 | 13 | 5 | 24 |
| *Medicago truncatula* | 2 | 1 | 7 | 2 | 17 |
| *Mimulus guttatus* | 1 | 1 | 0 | 1 | 0 |
| *Musa acuminata* | 1 | 2 | 0 | 1 | 0 |
| *Oryza sativa* | 1 | 1 | 0 | 1 | 0 |
| *Ostreococcus lucimarinus* | 0 | 0 | 0 | 0 | 0 |
| *Phaseolus vulgaris* | 2 | 2 | 1 | 2 | 8 |
| *Phoenix dactylifera* | 1 | 2 | 0 | 1 | 0 |
| *Physcomitrella patens* | 0 | 0 | 0 | 0 | 0 |
| *Picea abies* | 4 | 16 | 0 | 35 | 0 |
| *Pinus tadea* | 4 | 17 | 0 | 30 | 0 |
| *Populus trichocarpa* | 2 | 2 | 4 | 2 | 5 |
| *Prunus persica* | 2 | 1 | 3 | 1 | 11 |
| *Ricinus communis* | 1 | 1 | 1 | 1 | 3 |
| *Selaginella moellendorffii* | 0 | 0 | 0 | 0 | 0 |
| *Solanum lycopersicum* | 1 | 1 | 2 | 1 | 1 |

| Species | EDS1 | PAD4 | SAG101 | ADR1 | NRG1 |
|---|---|---|---|---|---|
| *Setaria italica* | 1 | 1 | 0 | 1 | 0 |
| *Solanum melongena* | 1 | 1 | 2 | 1 | 1 |
| *Solanum tuberosum* | 1 | 1 | 2 | 1 | 0 |
| *Sorghum bicolor* | 1 | 1 | 0 | 1 | 0 |
| *Spirodela polyrhiza* | 0 | 0 | 0 | 0 | 0 |
| *Thellungiella parvula* | 1 | 1 | 2 | 2 | 3 |
| *Theobroma cacao* | 1 | 2 | 1 | 1 | 1 |
| *Vitis vinifera* | 3 | 1 | 3 | 1 | 14 |
| *Volvox carteri* | 0 | 0 | 0 | 0 | 0 |
| *Zea mays* | 1 | 2 | 0 | 2 | 0 |
| **TOTAL count** | **76** | **89** | **82** | **143** | **156** |

Supplemental Table 2.

Analysis of selection pressure acting on EDS1 in *Brassicaceae*, *Asterids* and *Poaceae*

| **Clade** | **M3:M0 branch model ($2\Delta\ell$, $\chi^2$ p-value)** | **M1:M0 branch model ($2\Delta\ell$, $\chi^2$ p-value)** | **M2a:M1a site model ($2\Delta\ell$, $\chi^2$ p-value)** | **M8:M7 site model ($2\Delta\ell$, $\chi^2$ p-value)** | **M8:M8a site model ($2\Delta\ell$, $\chi^2$ p-value)** | **Branch-site model**<br>**Background:** most recent ancestor (MRA) for clade<br>**Foreground:** external branches | | |
|---|---|---|---|---|---|---|---|---|
| | Variable vs. uniform<br><br>*Is dN/dS variable among sequence sites?* | Variable neutral vs. uniform<br><br>*Is dN/dS variable among sequence sites?* | Positive selection vs. nearly neutral<br><br>More stringent than M8:M7 or M8:M8a<br><br>*Is there evidence for positive selection?* | Diverse selection regimes vs. no positive selection<br><br>*Is there evidence for positive selection?* | Diverse selection regimes vs. diverse & fixed ω1=1<br><br>*Is there evidence for positive selection?* | 0) Background: negative selection<br><br>Foreground: negative selection<br><br>(BEB class 1) | 1) Background: neutral selection<br><br>Foreground: neutral selection<br><br>(BEB class 2) | 2a) Background: negative selection<br><br>Foreground: positive selection<br><br>(BEB class 3)<br><br>2b) Background: neutral selection<br><br>Foreground: positive selection<br><br>(BEB class 4) |
| ***Brassicaceae***<br><br>(N=9 + *C. papaya*, *T. cacao*, *C. quinoa* as outgroups) | **624.9 (p=0)**<br>M0:<br>ω = 0.335<br>M3:<br>ω = 0.063 (*p=0.397*)<br>ω = 0.437 (*p=0.480*)<br>ω = 1.416 (*p=0.123*) | **76.7 (p<0.001)**<br>M1:<br>ω = 0.155 (*p=0.665*)<br>ω = 1.000 (*p=0.335*) | **14.1 (p=0.001)**<br>**BEB, P>95%: K215, K487**<br><br>M2:<br>ω = 0.159 (*p=0.660*)<br>ω = 1.000 (*p=0.325*)<br>ω = 3.311 (*p=0.015*) | **35.3 (p<0.001)**<br><br>**BEB, P>95%: K215, Q223, K487** | **16.5 (p<0.001)** | 421 (62.0%) | 228 (33.6%) | **2a) 17 (2.5%); sites with P>0.95: R16, Q223, N509***<br><br>**2b) 13 (1.9%); site with P>0.95: Q223** |
| ***Asterids***<br><br>(N=9 + *V. vinifera* and *B. vulgaris* as outgroups) | **449.5 (p=0)**<br>M0:<br>ω = 0.232<br>M3:<br>ω = 0.040 (*p=0.410*)<br>ω = 0.308 (*p=0.440*)<br>ω = 0.977 (*p=0.150*) | **85.6 (p<0.001)**<br>M1:<br>ω = 0.128 (*p=0.729*)<br>ω = 1.000 (*p=0.271*) | 0 (p=1)<br><br>M2:<br>ω = 0.128 (*p=0.729*)<br>ω = 1.000 (*p=0.173*)<br>ω = 1.000 (*p=0.098*) | **8.95 (p=0.006)**<br><br>**BEB, P>95%:** none | 1.9 (p=0.08) | 496 (76.8%) | 150 (23.2%) | **0 (0%)** |
| ***Poaceae***<br><br>(N=7 + *M. acuminate* and *A. comosus* as outgroups) | **1277.3 (p=0)**<br>M0:<br>ω = 1<br>M3:<br>ω = 0.000 (*p=0.231*)<br>ω = 0.156 (*p=0.561*)<br>ω = 0.610 (*p=0.207*) | **1075.1 (p=0)**<br>M1:<br>ω = 0.115 (*p=0.810*)<br>ω = 1.000 (*p=0.190*) | 0 (p=1)<br><br>M2:<br>ω = 0.115 (*p=0.810*)<br>ω = 1.000 (*p=0.118*)<br>ω = 1.000 (*p=0.072*) | 0.1 (p=0.965) | 0.1 (p=0.82) | 520 (88.4%) | 66 (11.3%) | **0 (0%)** |

* N509: 5 out of 9 sequences contain gaps at this codon.

Supplemental Table 3

List of oligonucleotides used in this study

| Name | Sequence 5'->3' | Use, comments |
|---|---|---|
| **nDL483** | CTCAACAGCCAGATTTTGCTC | LP for genotyping adr1-1 SAIL_842_B05 |
| **nDL484** | TCCTCGTCAATATCATGCCTC | RP for genotyping adr1-1 SAIL_842_B05 |
| **nDL485** | CGTGCTTGTTGCTTTAGGAAG | LP for genotyping adr1-L1 SAIL_302_C06 |
| **nDL486** | AAGTTCCCTCAGCTCCTCAAG | RP for genotyping adr1-L1 SAIL_302_C06 |
| **nDL487** | ATTTCGCTCCGGACTCTAAAG | LP for genotyping adr1-L2-4=SALK_126422 |
| **nDL488** | ATCGGTTGTCACCATCTCAAC | RP for genotyping adr1-L2-4=SALK_126422 |
| **nDL421** | gttttccttggtcattgcgc | Together with nDL596 detects insertion2 in NRG1.1 in n2a3 mutant background.<br>In combination with nDL597 – WT band only |
| **nDL596** | ccactATGCATAGAAATAtgactcttgg | Together with nDL421, genotyping of nrg1.1 in n2a3 |
| **nDL597** | GATTTCTCGTACCACTTG | Together with nDL421, nrg1.1 in n2a3 |
| **nDL577** | acataccttcgatctctgggt | nrg1.2 in n2a3 |
| **nDL578** | tcaataagaagctcgaccgtt | nrg1.2 in n2a3 |
| **JAD049** | ttcagtgatcccctttttcg | Together with JAD050 or JAD051 in step1 or 2, respectively, of nested PCR for T7E1 assay to detect nrg1.1 mutagenesis events |
| **JAD050** | GTGCATTGTAACCGTTGTGC | Together with JAD049 for T7E1 assays |
| **JAD051** | CCctgccaaaggaaactaca | Together with JAD049 for T7E1 assays; together with nDL421 to detect insertion1 in NRG1.1 in n2a3 mutant |
| **JAD052** | CCAGCACAACGGTTACAATG | Together with JAD053 or JAD054 in step1 or 2, respectively, of nested PCR for T7E1 assay to detect nrg1.1 mutagenesis events |
| **JAD053** | ATGAGCCCATGTCCAAAAAG | Together with JAD052 |
| **JAD054** | CTTTCCACCTGACCTTTCCA | Together with JAD052 |
| **JAD055** | GTTGCTGGAGCTTTGGTCTC | Together with JAD056 |
| **JAD056** | AATGTAATTGCTCCGCAACC | Together with JAD055 or JAD057 in step1 or 2, respectively, of nested PCR for T7E1 assay to detect nrg1.2 mutagenesis events |
| **JAD057** | GAGTCATTGCTTCCGGTGAT | Together with JAD056 |
| **JAD058** | AGGGATCCGAGTTCTTGCTT | Together with JAD059 or JAD060 in step1 or 2, respectively, of nested PCR for T7E1 assay to detect nrg1.2 mutagenesis events |
| **JAD059** | TCTTCGCGCTTATTTGTCct | Together with JAD058 |
| **JAD060** | GAAGCTAGGCTGCAGACGTT | Together with JAD058 |
| **AC251** | CACCATGCCGATGGACACCCCG | Together with AC252, TOPO cloning of HvEDS1 from cDNA |
| **AC252** | TTACGAAGGCACAAGTCTCGCG | Together with AC251, TOPO cloning of HvEDS1 from cDNA |
| **AC286** | caccatggaggacgcagaagaaaat | Together with AC254, TOPO cloning of HvPAD4 from cDNA |
| **AC254** | CTAAAACAACGACGCTGCTCTGGCT | Together with AC286, TOPO cloning of HvPAD4 from cDNA |
| **AC288** | caccATGCCGATGGACACTCCG | Together with AC289, TOPO cloning of BdEDS1 from cDNA |
| **AC289** | TTACGGAGGCATAAGCCTC | Together with AC288, TOPO cloning of BdEDS1 from cDNA |
| **AC290** | caccATGGAAGACGCAGAAGAAATTTC | Together with AC291, TOPO cloning of BdPAD4 from cDNA |
| **AC291** | TCAGAACTGAAATAATGATGATGTC | Together with AC290, TOPO cloning of BdPAD4 from cDNA |
| **StcEDS1_pE_UF** | caccATGGTGAAAATTGGAGAAGGAAT | Together with StcEDS1_pE_DR, TOPO cloning of StEDS1 from cDNA |
| **StcEDS1_pE_DR** | CTAAGGAGTTATTTTCCTTGATACC | Together with StcEDS1_pE_UF, TOPO cloning of StEDS1 from cDNA |
| **StPAD4_1_UF** | caccATGGAATCGGAAGCTTCATCGT | Together with StPAD4_1_DR, TOPO cloning of StEDS1 from cDNA |
| **StPAD4_1_DR** | TCAAGGAAACTGAGGTTGGAGC | Together with StPAD4_1_UF, TOPO cloning of StEDS1 from cDNA |
| **AC336** | CGAAAACTGGGCTCGTTACT | Detection of eds1a mutation in N. benthamiana, together with AC337 |
| **AC337** | CCACCAGTTTCCCATTTCCA | Detection of eds1a mutation in N. benthamiana, together with AC336 |
| **AC338** | TTATGATGGCTGCAGGTTCG | Detection of pad4 mutation in N. benthamiana, together with AC339 |
| **AC339** | ACGGAAGAAATTGTTTGTGCA | Detection of pad4 mutation in N. benthamiana, together with AC338 |
| **JS1114** | AAGTTGGTGGTGAGCTCAGA | Detection of sag101a-1 mutation in N. benthamiana, together with AC382 |
| **AC382** | GCAAGCTGAAACGCATGTAG | Detection of sag101a-1 mutation in N. benthamiana, together with JS1114 |
| **AC369** | CAAGTTCATGAATATCCACCAG | Detection of WT SAG101a in N. benthamiana, together with AC382 |
| **JS1120** | AAGTTGGTGGTGAGCTCAGA | Detection of sag101b-1 mutation in N. benthamiana, together with JS1121 |
| **JS1121** | GCCTCGTTAGTCAATCAGCTC | Detection of sag101b-1 mutation in N. benthamiana, together with JS1120 |
| **nDL372** | CACCATGAACGATTGGGCTAGTTTGGG | TOPO cloning of NRG1.1 AT5G66900.1, together with nDL373 |
| **nDL373** | TTAAAACATTTGAAGCAAGTTC | together with nDL372 |
| **nDL374** | CACCATGGTCGTGGTCGATTGGC | TOPO cloning of NRG1.2 AT5G66910.1, together with nDL375 |

| Name | Sequence 5’->3’ | Use, comments |
|---|---|---|
| **nDL375** | TTAAAACGTTAGAAGCAACTT | together with nDL374 |
| **nDL478** | GAACTTGCTTCAAATGTTTGCAAAGGGT GGGCGCGCCGA | Site-directed mutagenesis Stop->Ala in pENTR/D-TOPO AT5G66900.1 NRG1.1. Together with nDL479 |
| **nDL479** | TCGGCGCGCCCACCCTTTGCAAACATTT GAAGCAAGTTC | Together with nDL478 |
| **nDL480** | GAAGTTGCTTCTAACGTTTGCAAAGGGT GGGCGCGCCGACC | Site-directed mutagenesis Stop->Ala in pENTR/D-TOPO AT5G66900.1 NRG1.2. Together with nDL481 |
| **nDL481** | GGTCGGCGCGCCCACCCTTTGCAAACGT TAGAAGCAACTTC | Together with nDL480 |
| **nDL489** | caccatgcaattcgtctccttc | TOPO cloning of ext_NRG1.1 AT5G66900.2, together with nDL490 |
| **nDL490** | AAACATTTGAAGCAAGTTCAGGTTATG | Together with nDL489 |
| **nPVB031** | tacaGAAGACaaaATGCCGATGGACACCCC G | Golden gate cloning of HvEDS1 into level 0, incl. BbsI, CDS1 overhang, together with nPVB032 |
| **nPVB032** | atgtGAAGACttGAATACGAAGGCGGCGCA G | Golden gate cloning of HvEDS1 into level 0, mutates GAAGAC to GAATAC removes internal BbsI RS, together with nPVB031 |
| **nPVB033** | tacaGAAGACaaATTCCCGCCCACCTGGTC | Golden gate cloning of HvEDS1 into level 0, mutates GAAGAC to GAATAC removes internal BbsI RS, together with nPVB034 |
| **nPVB034** | tacaGAAGACaaAAGCCTACGAAGGCACA AGTCTCG | Together with nPVB033 |
| **nPVB070** | tacaGAAGACaaaATGGAGGACGCAGAAG AAAATTCCATGTTCGAaACCAGCCATG | Golden gate cloning of HvPAD4, together with nPVB071 |
| **nPVB071** | atgtGAAGACttGAATACGTCGGCGCCGCT CGCCACCGC | together with nPVB070 |
| **nPVB037** | tacaGAAGACaaATTCGCGCCGGTCGGC | Golden gate cloning of HvPAD4; together with nPVB038 |
| **nPVB038** | tacaGAAGACaaAAGCCTAAAACAACGACG CTG | Together with nPVB037 |
| **DB115** | CAAACTACCATCGAtATTTAAAGAACGAA G | Site-directed mutagenesis of AtEDS1 H476F, together with DB116 |
| **DB116** | CTTCGTTCTTTAAATaTCGATGGTAGTTT G | together with DB115 |
| **D167** | GAAGAGAATGACgagAAAGCAAACGTC | Site-directed mutagenesis of AtEDS1 F419E, together with D168 |
| **D168** | GACGTTTGCTTTctcGTCATTCTCTTC | together with D167 |
| **CLR017** | AGCTGCGCCGATGGTTTCTACAA | Detection of pKIR1.0 T-DNA in CRISPR mutants, together with CLR018 |
| **CLR018** | ATCGCCTCGCTCCAGTCAATG | Together with CLR018 |
| **gRNA2** | GGAAGTTTCCAAGAGTCAAG | Guide RNA for CRISPR-Cas9 mutagenesis of *AtNRG1.1* |
| **gRNA12** | TCATCAGGAGACGTGTTACA | Guide RNA for CRISPR-Cas9 mutagenesis of *AtNRG1.1* |
| **gRNA14** | GATACGAGATCCTTGGACAA | Guide RNA for CRISPR-Cas9 mutagenesis of *AtNRG1.1* |
| **gRNA3** | CAACGCTTCGAAACAAGCGG | Guide RNA for CRISPR-Cas9 mutagenesis of *AtNRG1.2* |
| **gRNA4** | GAAACTCCTTGACAATTCCG | Guide RNA for CRISPR-Cas9 mutagenesis of *AtNRG1.2* |
| **gRNA15** | CCTCTCGTAATTGAAGTAGT | Guide RNA for CRISPR-Cas9 mutagenesis of *AtNRG1.2* |
| **Y527** | CGTTTCTCTATTCCTTTGATACATTCTTTG ACTAAAACGC | Cloning PAD4-SAG101 chimera 1, with Y528 |
| **Y528** | GCGTTTTAGTCAAAGAATGTATCAAAGGA ATAGAGAAACG | Cloning PAD4-SAG101 chimera 1, with Y527 |
| **Y529** | CTTAAACATCATATCAAATCGCAGAATAG GAGCCCGAACA | Cloning PAD4-SAG101 chimera 2, with Y530 |
| **Y530** | TGTTCGGGCTCCTATTCTGCGATTTGATA TGATGTTTAAG | Cloning PAD4-SAG101 chimera 2, with Y529 |
| **Y531** | GCATTTCTTTTTGTACCACTCAATCTCaA GTCTTGCTGGC | Cloning PAD4-SAG101 chimera 3, with Y532 |
| **Y532** | GCCAGCAAGACTtGAGATTGAGTGGTAC AAAAAGAAATGC | Cloning PAD4-SAG101 chimera 3, with Y531 |
| **Y533** | TGAACCGATCGTAGTAACCTAGCTGCTC TTCTGATGCA | Cloning PAD4-SAG101 chimera 4, with Y534 |
| **Y534** | TGCATCAGAAGAGCAGCTAGGTTACTAC GATCGGTTCA | Cloning PAD4-SAG101 chimera 4, with Y533 |

Supplemental Table 4

List of species used to infer plant protein orthogroups

| Species | Downloaded on | Database | NCBI tax ID |
|---|---|---|---|
| ***Arabidopsis thaliana*** | 29. Jul 14 | TAIR10 | 3702 |
| ***Actinidia chinensis*** | 21. Nov 14 | http://bioinfo.bti.cornell.edu/cgi-bin/kiwi/home.cgi | 3625 |
| ***Amborella trichopoda*** | 29. Jul 14 | Phytozome | 13333 |
| ***Aquilegia coerulea*** | 29. Jul 14 | Phytozome | 218851 |
| ***Arabidopsis halleri*** | 29. Jul 14 | Phytozome | 81970 |
| ***Arabidopsis lyrata*** | 29. Jul 14 | Phytozome | 59689 |
| ***Arachis ipaensis*** | 25. Nov 14 | PeanutBase | 130454 |
| ***Beta vulgaris*** | 21. Nov 14 | http://bvseq.boku.ac.at/index.shtml | 161934 |
| ***Boechera stricta*** | 29. Jul 14 | Phytozome | 72658 |
| ***Brachypodium distachyon*** | 29. Jul 14 | Phytozome | 15368 |
| ***Brassica oleracea*** | 28. Jan 15 | Brassica; http://brassicadb.org/brad/downloadOverview.php | 3712 |
| ***Brassica rapa*** | 29. Jul 14 | Phytozome | 3711 |
| ***Capsella grandiflora*** | 29. Jul 14 | Phytozome | 264402 |
| ***Capsella rubella*** | 29. Jul 14 | Phytozome | 81985 |
| ***Capsicum annum*** | 20. Nov 14 | http://peppersequence.genomics.cn/page/species/download.jsp | 4072 |
| ***Carica papaya*** | 29. Jul 14 | Phytozome | 3649 |
| ***Castanea mollissima*** | 20. Jun 14 | https://www.hardwoodgenomics.org/Genome-assembly/1962958 | 60419 |
| ***Citrus clementina*** | 29. Jul 14 | Phytozome | 85681 |
| ***Citrus sinensis*** | 29. Jul 14 | Phytozome | 2711 |
| ***Coccomyxa subellipsoidea*** | 29. Jul 14 | Phytozome | 248742 |
| ***Coffea canephora*** | 20. Nov 14 | http://coffee-genome.org/coffeacanephora | 49390 |
| ***Cucumis sativus*** | 29. Jul 14 | Phytozome | 3659 |
| ***Eutrema salsugineum*** | 29. Jul 14 | Phytozome | 72664 |
| ***Fragaria vesca*** | 29. Jul 14 | Phytozome | 57918 |
| ***Gossypium raimondii*** | 29. Jul 14 | Phytozome | 29730 |
| ***Linum usitatissimum*** | 29. Jul 14 | Phytozome | 4006 |
| ***Malus domestica*** | 29. Jul 14 | Phytozome | 3750 |
| ***Medicago truncatula*** | 29. Jul 14 | Phytozome | 3880 |
| ***Mimulus guttatus*** | 21. Nov 14 | Phytozome | 4155 |
| ***Musa acuminata*** | 11. Dec 2014 | Phytozome | 4641 |
| ***Oryza sativa*** | 29. Jul 14 | Phytozome | 4530 |
| ***Ostreococcus lucimarinus*** | 29. Jul 14 | Phytozome | 242159 |
| ***Phaseolus vulgaris*** | 29. Jul 14 | Phytozome | 3885 |
| ***Phoenix dactylifera*** | 11. Dec 2014 | http://qatar-weill.cornell.edu/research/research-highlights/date-palm-research-program/date-palm-draft-sequence | 42345 |
| ***Physcomitrella patens*** | 29. Jul 14 | Phytozome | 3218 |
| ***Picea abies*** | 30.10.2014 | Congenie; http://congenie.org/start | 3329 |

| Species | Downloaded on | Database | NCBI tax ID |
|---|---|---|---|
| ***Pinus tadea*** | 12. Nov 14 | https://pinerefseq.faculty.ucdavis.edu/ | 3352 |
| ***Populus trichocarpa*** | 29. Jul 14 | Phytozome | 3694 |
| ***Prunus persica*** | 29. Jul 14 | Phytozome | 3760 |
| ***Ricinus communis*** | 29. Jul 14 | Phytozome | 3988 |
| ***Selaginella moellendorffii*** | 29. Jul 14 | Phytozome | 88036 |
| ***Solanum lycopersicum*** | 29. Jul 14 | Phytozome | 4081 |
| ***Setaria italica*** | 29. Jul 14 | Phytozome | 4555 |
| ***Solanum melongena*** | 21. Nov 14 | http://eggplant.kazusa.or.jp/ | 4111 |
| ***Solanum tuberosum*** | 29. Jul 14 | Phytozome | 4113 |
| ***Sorghum bicolor*** | 29. Jul 14 | Phytozome | 4558 |
| ***Spirodela polyrhiza*** | 29. Jul 14 | Phytozome | 29656 |
| ***Thellungiella parvula*** | 28. Jan 15 | http://thellungiella.org/data/ | 98039 |
| ***Theobroma cacao*** | 29. Jul 14 | Phytozome | 3641 |
| ***Vitis vinifera*** | 29. Jul 14 | Phytozome | 29760 |
| ***Volvox carteri*** | 29. Jul 14 | Phytozome | 3067 |
| ***Zea mays*** | 29. Jul 14 | Phytozome | 4577 |

3

Supplemental Table 5

List of species used to prepare EP domain HMM profile

| **Species** | **Downloaded on** | **Database** |
|---|---|---|
| ***Physcomitrella patens*** | 07. Aug 2018 | Phytozome.org via BioMart |
| ***Sphagnum fallax*** | 07. Aug 2018 | Phytozome.org via BioMart |
| ***Marchantia polymorpha*** | 07. Aug 2018 | Phytozome.org via BioMart |
| ***Selaginella moellendorffii*** | 07. Aug 2018 | Phytozome.org via BioMart |
| ***Ananas comosus*** | 07. Aug 2018 | Phytozome.org via BioMart |
| ***Musa acuminata*** | 07. Aug 2018 | Phytozome.org via BioMart |
| ***Amborella trichopoda*** | 07. Aug 2018 | Phytozome.org via BioMart |
| ***Oryza sativa*** | 07. Aug 2018 | Phytozome.org via BioMart |
| ***Brachypodium distachyon*** | 07. Aug 2018 | Phytozome.org via BioMart |
| ***Zea mays*** | 07. Aug 2018 | Phytozome.org via BioMart |
| ***Sorghum bicolor*** | 07. Aug 2018 | Phytozome.org via BioMart |
| ***Aquilegia coerulea*** | 07. Aug 2018 | Phytozome.org via BioMart |
| ***Amaranthus hypochondriacus*** | 07. Aug 2018 | Phytozome.org via BioMart |
| ***Kalanchoe laxiflora*** | 07. Aug 2018 | Phytozome.org via BioMart |
| ***Daucus carota*** | 07. Aug 2018 | Phytozome.org via BioMart |
| ***Solanum tuberosum*** | 07. Aug 2018 | Phytozome.org via BioMart |
| ***Solanum lycopersicum*** | 07. Aug 2018 | Phytozome.org via BioMart |
| ***Eucalyptus grandis*** | 07. Aug 2018 | Phytozome.org via BioMart |
| ***Vitis vinifera* Genoscope** | 07. Aug 2018 | Phytozome.org via BioMart |
| ***Linum usitatissimum*** | 07. Aug 2018 | Phytozome.org via BioMart |
| ***Populus trichocarpa*** | 07. Aug 2018 | Phytozome.org via BioMart |
| ***Citrus clementina*** | 07. Aug 2018 | Phytozome.org via BioMart |
| ***Theobroma cacao*** | 07. Aug 2018 | Phytozome.org via BioMart |
| ***Carica papaya*** | 07. Aug 2018 | Phytozome.org via BioMart |
| ***Arabidopsis thaliana*** | 07. Aug 2018 | Phytozome.org via BioMart |
| ***Capsella rubella*** | 07. Aug 2018 | Phytozome.org via BioMart |
| ***Cucumis sativus*** | 07. Aug 2018 | Phytozome.org via BioMart |
| ***Fragaria vesca*** | 07. Aug 2018 | Phytozome.org via BioMart |
| ***Glycine max*** | 07. Aug 2018 | Phytozome.org via BioMart |
| ***Volvox carteri*** | 07. Aug 2018 | Phytozome.org via BioMart |
| ***Micromonas sp.* RCC299** | 07. Aug 2018 | Phytozome.org via BioMart |
| ***Chlamydomonas reinhardtii*** | 07. Aug 2018 | Phytozome.org via BioMart |
